# Structure-based virtual screening reveals new metabolite interactions and modulators of human glucose-6-phosphate isomerase

**DOI:** 10.64898/2026.10.06.757072

**Authors:** Yiming Yang Jónatansdóttir, Knut Teigen, Jens G. Hjörleifsson

## Abstract

Exploring the ability of metabolic enzymes to interact with diverse small molecules can provide insights into their ligand recognition and activity regulation. Glucose-6-phosphate isomerase (GPI) is an enzyme that catalyzes the second step of glycolysis – a critical node that links glucose metabolism to other cytosolic pathways, such as pentose phosphate and hexosamine biosynthetic pathways. Despite its importance in cellular metabolism, little is known about the broad ligandability of GPI. Here, we performed a structure-based virtual screening using molecular docking to identify new modulators predicted to engage with the human GPI active site. Enzyme kinetic and binding studies confirmed eleven compounds as active-site binders of GPI, exhibiting inhibitory activities and binding affinities in upper micromolar to sub-millimolar range. The binding of several ligands to GPI active site was further supported by X-ray crystallographic data. In addition, *in vitro* assays were performed to evaluate the modulatory effects of the GPI hits on cellular glycolysis and viability. The identified inhibitors included the mono-phosphorylated metabolite of ribavirin, which inhibited glycolytic flux and cell viability; two compounds with phosphate bioisosteres; and endogenous phosphate-containing metabolites from the pentose phosphate, hexosamine, nucleotide, and mevalonate pathways. Collectively, these findings expanded the repertoire of human GPI ligands and revealed previously unrecognized GPI-metabolite interactions that may contribute to modulation of glycolytic flux.

## Introduction

Glycolysis enzymes play central roles in cellular metabolism by regulating metabolic flux required for energy generation and biosynthetic pathways. Dysregulation of these enzymes is implicated in a range of diseases, including cancer, metabolic disorders, and neurodegenerative conditions. Human glucose-6-phosphate isomerase (GPI; EC5.3.1.9) is one such enzyme, whose role extends beyond its canonical enzymatic function, making it a target of considerable diagnostic and therapeutic interest. Elevated GPI expression is associated with tumorigenesis and metastasis in multiple cancer types [1–3], while deficiencies in GPI activity can lead to hemolytic anemia and neurological disorders [3–5]. The canonical role of GPI in glycolysis involves the reversible isomerization of glucose-6-phosphate (G6P) to fructose-6-phosphate (F6P), a step that bridges glycolysis to pentose phosphate pathway (PPP) and hexosamine biosynthesis (HBP). GPI also exhibit moonlighting functions that include neuroleukin (NLK) [6], autocrine motility factor (AMF) [7], and cell differentiation mediator [8].

Glycolysis is elevated in many tumor cells, supplying the energy and biosynthetic precursors for unrestrained growth, conferring protection against oxidative stress, while shaping a tumor microenvironment that promotes immune evasion and invasion into surrounding tissues. GPI is ubiquitously overexpressed across 14 cancer classes [1], where it accelerates glycolytic flux in concert with other pathway enzymes and drives tumor cell migration [9], depending on its cellular localization. When secreted by cancer cells, GPI/AMF interacts with a cell-surface receptor – AMF receptor or glycoprotein 78 (AMFR/gp78) [10]. This interaction activates pro-migratory signaling pathways that induce cytoskeletal rearrangement, which enhances cell motility, and extracellular matrix (ECM) remodeling, promoting invasive behavior [11].

GPI is catalytically active as a homodimer, with each monomer containing an identical active site, wherein polar and charged residues facilitate substrate binding and catalysis. The active site is currently the only known ligandable regulatory region of GPI. High-resolution crystallographic studies have revealed residues relevant for ligand binding, with Thr214, Thr215, Glu358, His389, and Lys519 being the key players [12], stabilizing ligand through hydrogen bonds and electrostatic interactions. The catalytic mechanism of GPI involves proton transfers that facilitate isomerization of G6P to F6P, proceeding through a cis-enediol intermediate – an open-chain sugar phosphate [13]. Kinetic studies have characterized the ligand specificity of mammalian GPI, demonstrating a strong preference for phosphorylated compounds, particularly linear sugars comprising 4-6 carbons [14]. Beyond catalytic inhibition, emerging evidence suggests that active-site blockage can influence the AMF cytokine function of GPI [15]. Inhibitors such as erythrose-4-phosphate (E4P) and 6-phosphogluconate (6-PGA) have been shown to suppress not only isomerase activity but also AMF-mediated signaling, presumably by interfering with interactions between extracellular GPI/AMF and its cognate receptor – AMFR/gp78 [10]. Such functional overlap highlights the potential of GPI active-site inhibition in modulating both enzymatic and cytokine functions involved in malignant growth and metastasis.

In this study, we report new active-site inhibitors of human GPI, expanding the chemical space beyond known inhibitors and new possibilities for ligand optimization. In our previous work, an experimental screen of 800 metabolite compounds against human GPI identified a few weak binders with limited potential for inhibitor design [16]. To explore a broader chemical space and improve hit identification, we employed virtual screening using Schrödinger’s Glide docking [17,18] to evaluate the binding potential of diverse small molecules within the GPI active site and prioritize candidates for experimental validation (Fig. 1). Compound libraries were retrieved from several chemical databases, including Zinc15/20, Enamine, ChEMBL, PubChem, Mcule, and eMolecules. The screening libraries were designed to cover two distinct regions of chemical space, comprising drug-like and hydrophilic compounds. Experimental validation of the hits was achieved with enzyme activity assay and binding studies using microscale thermophoresis (MST) and differential scanning fluorometry (DSF), followed by X-ray crystallography to elucidate their binding modes. The docking study revealed eleven new competitive inhibitors of human GPI, including an active metabolite of ribavirin, two inhibitors with phosphate bioisosteres, and phosphorylated endogenous metabolites from the pentose phosphate, nucleotide, hexosamine, and mevalonate metabolism. Together, these findings provide new insights into GPI ligand recognition and highlight the potential of targeting this enzyme through both metabolite-inspired and phosphate-substitution strategies.

**Figure 1:**
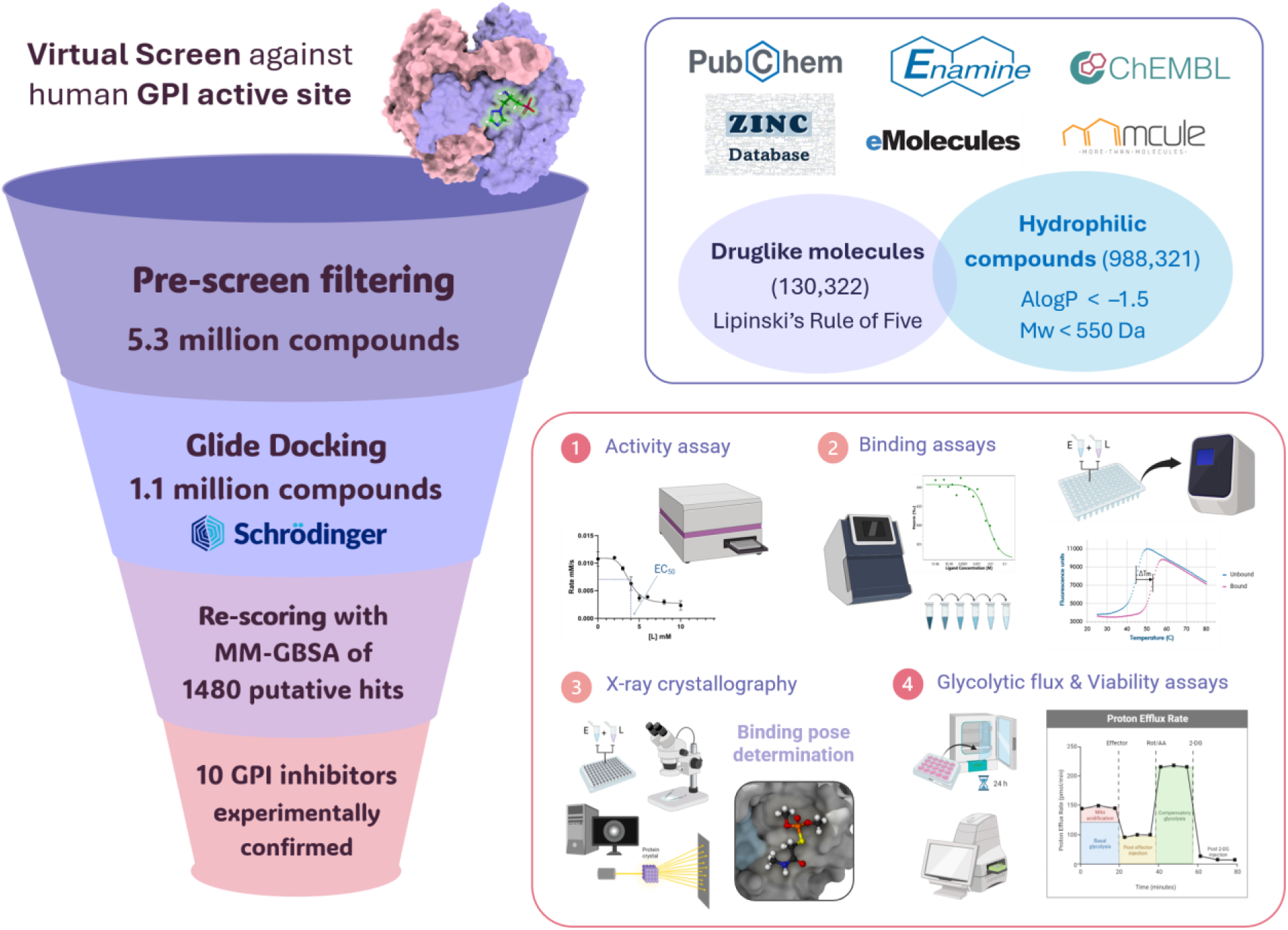
Structure-based virtual screening (SBVS) workflow for the identification of human glucose-6-phosphate isomerase (GPI) inhibitors. Compound libraries were filtered according to user-defined physicochemical properties and docked into the GPI active site using Schrödinger Glide. Top-ranked compounds were subsequently re-scored using Prime MM-GBSA and manually curated for experimental validation. Selected candidates were evaluated by enzymatic inhibition, ligand-binding, and structural studies, followed by glycolytic flux and cell viability assays.

## Results

### New GPI inhibitors identified through targeted chemical space exploration

To identify new binders of the GPI active site, two virtual screens were performed, each designed to explore a distinct chemical space (Fig.1). The first screen focused on drug-like compounds, using Lipinski’s rule of five as selection criteria to enrich for molecules with physicochemical properties of oral drug-like compounds. The second screen targeted hydrophilic space, where the libraries were filtered using AlogP values between −6.0 and −1.5 and molecular weights (Mw) of 100 to 550 Da, reflecting the polarity and size of known GPI active-site binders, respectively. For hit selection, Glide docking and Prime MM-GBSA scores of known GPI inhibitors were used as reference values, with the least negative scores serving as cutoff values (see Methods). Compounds were prioritized if their predicted scores met or exceeded these criteria. The Emodel scoring function was employed for ranking the docked ligands, as it showed better enrichment performance than the GlideScore function.

In the first screen, which was confined to drug-like chemical space, 130,322 compounds were docked into the GPI active site. Based on commercial availability, a total of 12 drug-like compounds were selected for experimental evaluation (Suppl. Table S1). Despite showing good docking scores, only one compound in the drug-like docking screen, dimethylallyl pyrophosphate (DMAPP), exhibited a dose-dependent response in the MST binding assay, with an estimated binding affinity (K_d_) of 7.4 mM. Moreover, co-crystallization of GPI and DMAPP revealed partial electron density for the ligand in the active site, with the pyrophosphate group aligning with the observed density (shown in later section). However, its lack of enzymatic inhibition, even after prolonged pre-incubation time, indicates slow binding kinetics and rapid dissociation when substrate is present. All the remaining drug-like compounds did not cause apparent changes in GPI activity and MST signal within the tested concentration range (0.01 – 40 mM) and showed no electron density in crystals soaked and/or co-crystallized with the ligand. Together, these observations indicate that these compounds are likely false-positive hits. The Glide scoring functions (Gscore and Emodel) showed a tendency to favor those with higher molecular weight, resulting in artificially high docking scores for larger compounds despite missing charges and hydrogen-bonding partners critical for ligand binding.

The lack of activity among the drug-like compounds further suggests that this chemical space may be poorly suited for targeting GPI active site, which has evolved to accommodate charged, phosphate-based ligands. However, not all phosphate-containing compounds are capable of binding GPI. Of the tested drug-like compounds, two contained phosphate groups, but only DMAPP exhibited binding, whereas the second compound carrying a pyridine scaffold with hydrophobic substituents showed no detectable interaction (Suppl. Table S1). This suggests that the presence of a phosphate moiety is not always sufficient for recognition by the GPI active site, and the overall polarity and charge distribution of the ligands are critical for binding. To improve hit rate, the screening strategy was shifted toward more hydrophilic, phosphate-enriched chemical space that better matches the physicochemical properties of the GPI active site. In this attempt, 988,321 hydrophilic compounds were screened and over a thousand showed favorable docking scores. These compounds were clustered and representatives were re-ranked by prime MM-GBSA analysis, and curated based on synthetic trackability, reducing to 193 candidates, of which 20 were purchased for experimental validation (Suppl. Table S2). Among these, ten compounds were confirmed as true hits, exhibiting dose-dependent responses in both activity and/or binding assays (Fig. 2-5). However, there was no significant correlation observed between docking/MM-GBSA scores and experimental inhibitory potency (Suppl. Fig. S1). The kinetic data, binding affinities, and docking scores for all identified GPI hits in this study are summarized in Table 1.

**Figure 2:**
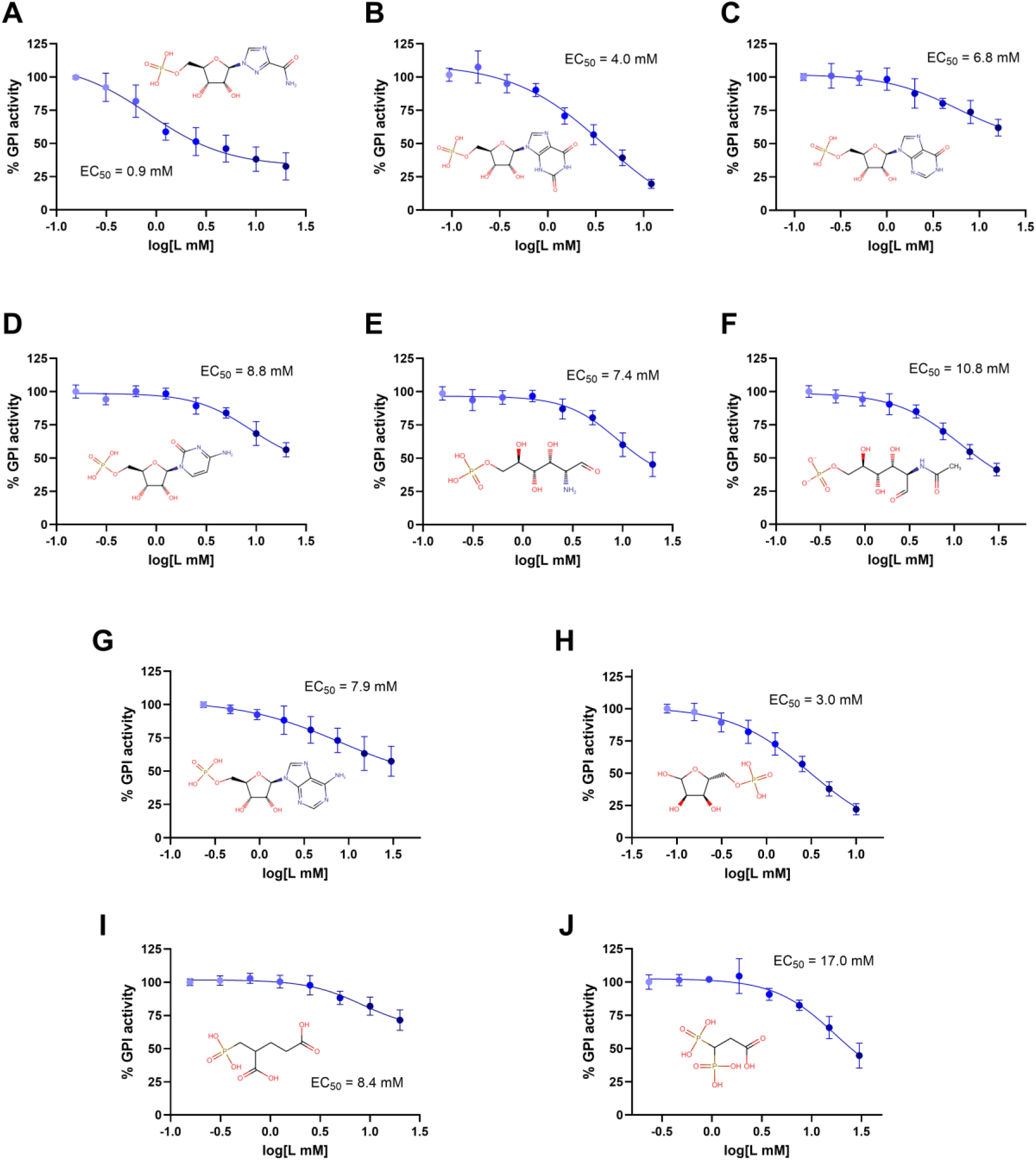
Inhibition of human glucose-6-phosphate isomerase (GPI) by compounds identified from virtual screening. (A) RMP. (B) XMP. (C) IMP. (D) CMP. (E) GlcN6P. (F) NAG6P. (G) AMP. (H) R5P. (I) 2PMPA. (J) 33DP. Dose-response curves obtained using enzyme-coupled assay. Each data point represents mean ± error bar denoting SD of two independent duplicates measured on two days. Data were fitted using the log(inhibitor) vs. response – Variable slope (four parameters) and EC_50_ values were determined in Prism GraphPad.

**Table 1:** Summary of biochemical, biophysical, structural, and computational characterization of selected GPI ligands, including potency (EC₅₀, K_i_), binding affinity (K_d_) determined by MST and DSF assays, mode of action (MOA), qualitative structural validation (electron density) by X-ray crystallography, and Glide Emodel scores for the best-ranked poses from Glide SP docking. The EC_50_ and K_d v_alues are expressed as mean ± standard deviation of two independent duplicates or triplicates, and K_i_ values as mean ± standard deviation of two replicates.

| Compound | $EC_{50}$ (mM) | MST - $K_d$ (mM) | $K_i$ (mM) | MOA | DSF - $K_d$ (mM) | Electron density | Docking scores (kJ/mol) |
| --- | --- | --- | --- | --- | --- | --- | --- |
| **Ribavirin 5'-monophosphate (RMP)** | 0.9 ± 0.1 | 0.9 ± 0.1 | 1.2 ± 0.3 | Competitive | 2.6 ± 0.1 | Full ligand | -147.14 |
| **Xanthosine 5'-monophosphate (XMP)** | 4.0 ± 2.7 | 0.4 ± 0.1 | 1.0 ± 0.1 | Competitive | N.A. | Partial densities | -152.91 |
| **Inosine 5'-monophosphate (IMP)** | 6.8 ± 2.8 | 1.4 ± 0.6 | 6.9 ± 0.7 | Competitive | 9.0 ± 0.1 | Partial densities | -146.17 |
| **Cytidine 5'-monophosphate (CMP)** | 8.8 ± 1.1 | 1.3 ± 0.4 | 13.4 ± 3.1 | Competitive | 10.8 ± 0.2 | Full ligand | -151.07 |
| **D-Glucosamine 6-phosphate (GlcN6P)** | 7.4 ± 0.8 | 1.5 ± 0.3 | 2.4 ± 0.3 | Mixed | 7.5 ± 0.1 | Full ligand | -118.04 |
| **N-Acetyl-D-glucosamine 6-phosphate (NAG6P)** | 10.8 ± 2.4 | 4.6 ± 0.9 | 4.8 ± 0.4 | Competitive | 8.8 ± 0.2 | Full ligand | -148.83 |
| **Adenosine 5'-monophosphate (AMP)\*** | 7.9 ± 2.9 | 19.0 ± 2.3 | 24.2 ± 3.8 | Competitive | N.A.*** | Partial densities | -151.65 |
| **D-Ribose-5-phosphate (R5P)** | 3.0 ± 0.3 | 3.1 ± 0.7 | 0.6 ± 0.2 | Competitive | 7.2 ± 0.1 | N.D. | -117.24 |
| **2-Phosphonomethyl pentanedioic acid (2PMPA)** | 8.4 ± 1.8 | 7.8 ± 0.7 | 5.6 ± 1.3 | Competitive | N.A.** | N.D. | -125.28 |
| **3,3-diphosphono-propanoic acid (33DP)** | 17.0 ± 3.5 | 16.0 ± 3.0 | 20.5 ± 3.1 | Mixed | N.A | N.D. | -121.44 |
| **Dimethylallyl pyrophosphate (DMAPP)** | N.A. | $7.4 \pm 0.8$ | N.A. | N.D. | N.A.*** | Partial densities | -110.83 |
| **Methylene diphosphonic acid (MDP)** | $7.8 \pm 3.1$ | $9.1 \pm 0.3$ | N.D. | N.D. | N.D. | N.A. | - |
| **Phosphonoacetic acid (PAE)** | N.A. | $5.9 \pm 0.4$ | N.D. | N.D. | N.D. | N.A. | - |

\* Moderate inhibition on coupling enzyme – G6PDH; N.D (not determined); N.A. (not available).
DSF results: \*\*upshift and \*\*\*downshift in $T_m \geq 0.5$ °C with respect to control, at least one of the tested concentrations (supplement data).

### Identified GPI hits act as competitive inhibitors

Michaelis-Menten kinetic measurements were carried out in gluconeogenesis direction (F6P to G6P) to determine the mechanism of action for the identified ligands (Fig. 3 & 4). Seven ligands displayed purely or near-competitive inhibition, while three compounds (XMP, GlcN6P, and 33DP) behaved as competitive inhibitors at lower concentrations but showed mixed-mode inhibition at higher concentrations. Overall, the ligands exhibited dissociation constants (K_d_) and inhibitory constants (K_i_) falling within high-micromolar (µM) to sub-millimolar (mM) range, indicating moderate-to-weak binding affinities. D-ribose-5-phosphate (R5P) emerged as the strongest inhibitor with K_i_ of ∼0.6 mM, followed by xanthosine monophosphate (XMP) and ribavirin 5’-monophosphate (RMP) with K_i_ values of 1.0 and 1.2 mM, respectively. Glucosamine-6-phosphate (GlcN6P; K_i_ = 2.4 mM) and N-acetyl-D-glucosamine-6-phosphate (NAG6P; K_i_ = 4.8 mM) displayed moderate inhibitory activity. Weaker inhibition was observed for 2-phosphonomethyl pentanedioic acid (2PMPA; Ki = 5.6 mM), inosine 5′-monophosphate (IMP; Ki = 6.9 mM), and cytidine 5′-monophosphate (CMP; Ki = 13.4 mM). Compared to the ligands above, 3,3-diphosphono-propanoic acid (33DP; K_i_ = 20.5 mM) and adenosine 5’-monophosphate (AMP; K_i_ = 24.2 mM) showed the weakest inhibition.

**Figure 3:**
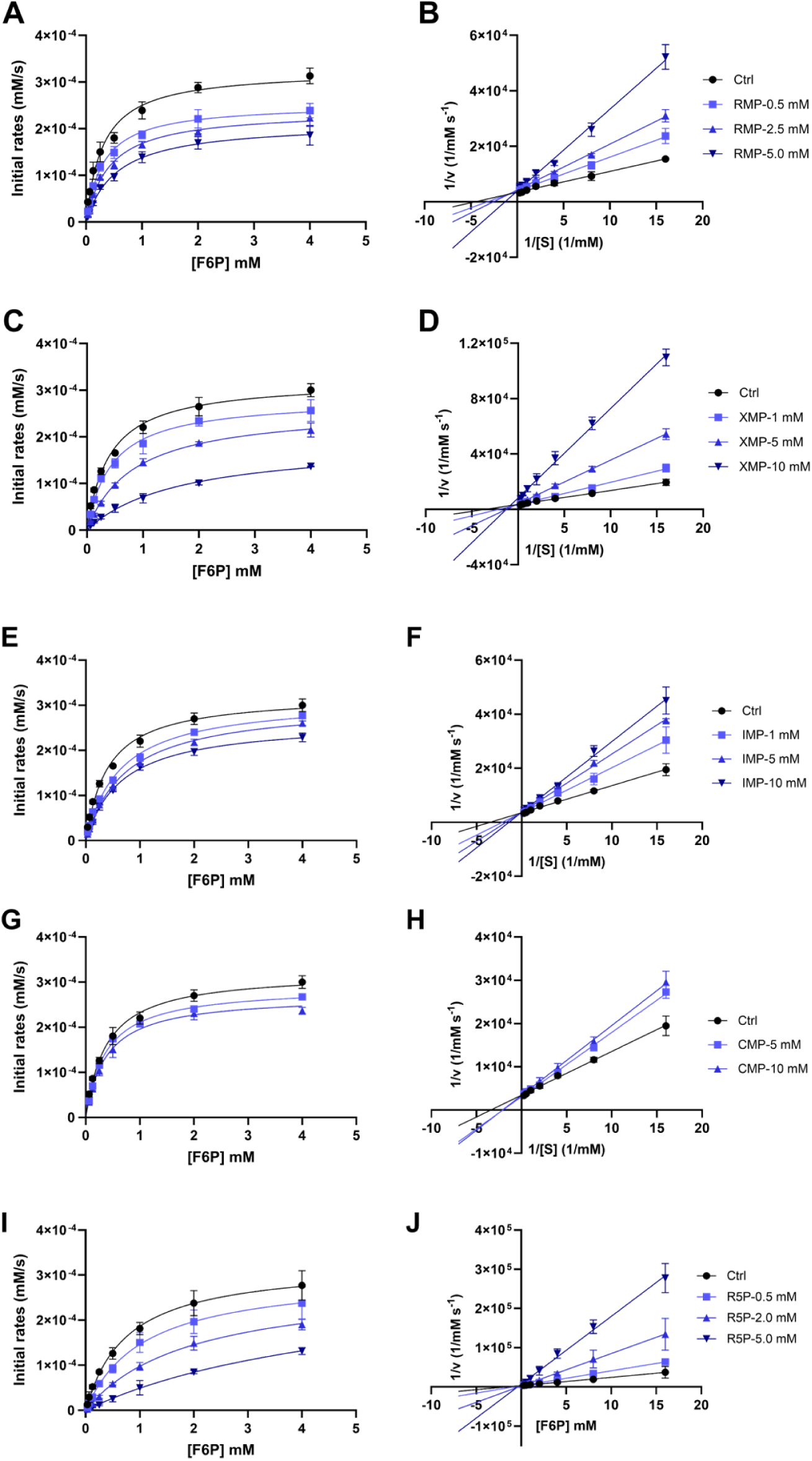
Michaelis–Menten kinetics and Lineweaver–Burk analyses of human glucose-6-phosphate isomerase (GPI) inhibition by (A, B) RMP, (C, D) XMP, (E, F) IMP, (G, H) CMP. (I, J) R5P. GPI activity was measured over a range of eight substrate concentrations (0.05–4 mM F6P). Data points represent the mean of duplicate measurements, and error bars indicate the standard deviation (± SD). Data were fitted using Michaelis–Menten nonlinear regression and linear regression for Lineweaver-Burk in GraphPad Prism.

**Figure 4:**
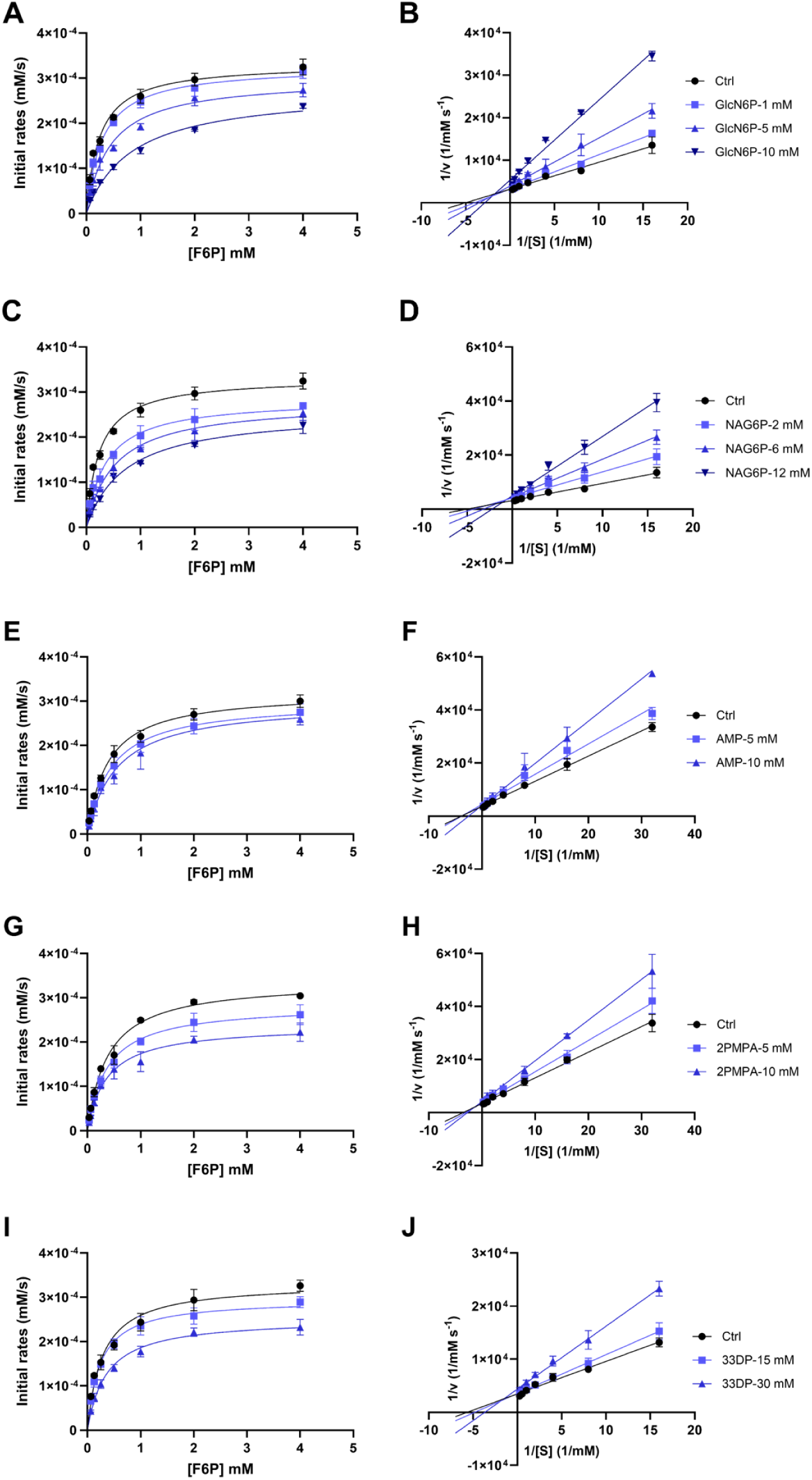
Michaelis–Menten kinetics and Lineweaver–Burk analyses of human glucose-6-phosphate isomerase (GPI) inhibition by (A, B) GlcN6P. (C, D) NAG6P. (E, F) AMP. (G, H) 2PMPA. (I, J) 33DP. GPI activity was measured over a range of eight substrate concentrations (0.05–4 mM F6P). Data points represent the mean of duplicate measurements, and error bars indicate the standard deviation (± SD). Data were fitted using Michaelis–Menten nonlinear regression and linear regression for Lineweaver–Burk in GraphPad Prism.

**Figure 5:**
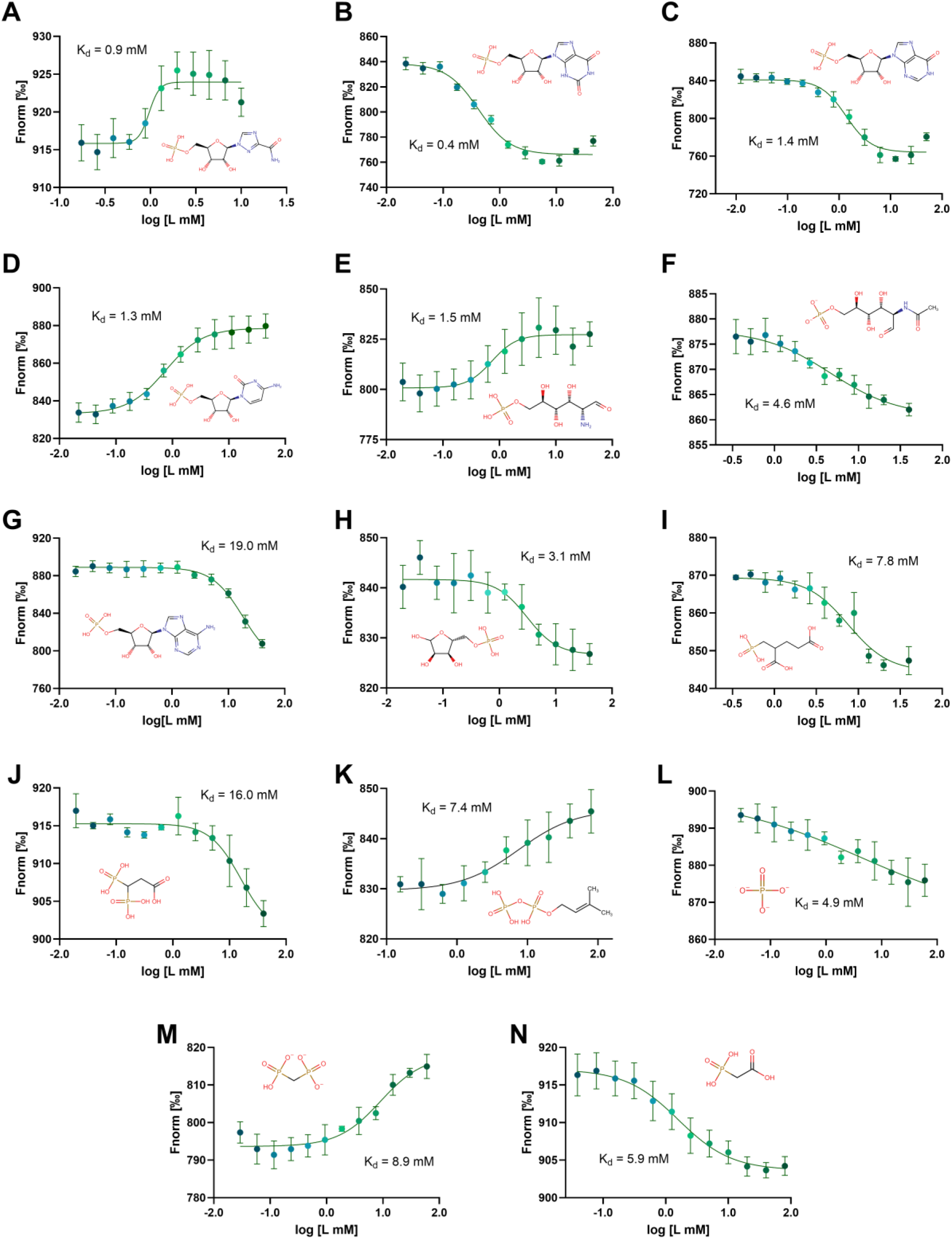
Binding of identified hits to human glucose-6-phosphate isomerase (GPI). Dose-response curves obtained using label-free microscale thermophoresis (MST). (A) RMP. (B) XMP. (C) IMP. (D) CMP. (E) GlcN6P. (F) NAG6P. (G) AMP. (H) R5P. (I) 2PMPA. (J) 33DP. (K) DMAPP. (L) Inorganic phosphate (PO₄). (M) MDP. (N) PAE. Compounds were titrated against GPI by a 2/3-fold serial dilution starting at 30-45 mM for metabolite hits and 60 mM for phosphate and bioisosteres (PO_4_, MDP and PAE). Data were fitted using the Sigmoidal dose-response (variable slope) in Prism GraphPad. Each data point represents mean ± SD of two independent triplicates or duplicates.

### Binding of the GPI ligands confirmed by orthogonal assays

The binding affinities (K_d_) of the compounds were primarily determined using label-free MST assay (Fig. 5). For most compounds, the MST-derived K_d_ were similar to their K_i_ values, except for R5P, IMP and CMP (Table 1). The estimated K_d_ of R5P was 5-fold higher than its Ki value, whereas IMP and CMP exhibited lower K_d_ relative to their K_i_ values. As weak interactions require high ligand concentrations that can introduce assay variability, the resulting affinity estimates may be subject to greater uncertainty, contributing to differences in K_d_ and K_i_ values. In addition, the discrepancy observed for IMP and CMP may be attributed to ligand-induced changes in the target’s photobleaching behavior in the MST measurement. In both cases, the photobleaching rate decreased with increasing ligand concentration. MST measurement repeated, following protein denaturation, suggested that these changes in target’s fluorescent properties were ligand-induced rather than binding-specific, and may have contributed to an apparently lower K_d_ values. Besides the hits identified from hydrophilic library screening, two putative phosphate mimics, methylene diphosphonate (MDP) and phosphonoacetate (PAE), and inorganic phosphate were assessed for their binding to GPI using the MST assay (Fig. 5L-N). MDP and PAE exhibited binding affinities of 9.1 mM and 5.9 mM, respectively, in the same range as inorganic phosphate (K_d_ = 4.9 mM).

To further validate their binding, DSF thermal shift assays were performed at 2-3 ligand concentrations in triplicate for all ligands, including MDP and PAE (Suppl. Fig. S2 & S3). An apparent shift in melting temperature (T_m_) relative to the control was observed for all ligands except XMP, MDP and PAE. Seven ligands caused an increase in T_m_, whereas two ligands (AMP and DMAPP) resulted in T_m_ decrease. The T_m_ shifts observed for the majority of ligands were concentration-dependent, reflecting ligand-induced changes in GPI thermal stability. However, three ligands did not produce a consistent dose-dependent response. This variability is consistent with previous observation that DSF response of GPI can differ between ligands, with even strong binders such as 6-phosphogluconate (6PG) exhibiting atypical bell-shaped dose– response patterns [16]. In addition, binding affinities of five ligands were determined by 8-point titration DSF experiment (Table 1 & Suppl. Fig. S4). All five compounds induced a dose-dependent increase in the T_m_ of GPI, indicating thermal stabilization upon ligand binding.

### Crystallographic data revealed electron densities for new GPI ligands

X-ray crystallography was employed to fully confirm the binding of the ligands to GPI and elucidate their binding modes. Eight ligands were successfully co-crystallized with GPI and all of them were shown to be active-site binders (Fig. 6). Among them, RMP, CMP, GlcN6P, and NAG6P showed well-defined electron densities, and the co-crystal structures were deposited. The remaining four ligands – XMP, IMP, AMP, and DMAPP exhibited only partial occupancy in the density maps. For XMP, IMP, and AMP, only the ribose phosphate moiety could be confidently modeled into the electron density, whereas the nucleobases remained poorly defined. Similarly, for DMAPP, only the pyrophosphate moiety fitted into the density map, while the isoprenoid chain lacked well-defined electron density. Nevertheless, the DMAPP-and XMP-bound structures were of sufficient quality for PDB deposition. Crystal structures of human GPI in complex with the selected ligands were solved to resolutions ranging from 1.6 to 2.1 Å. Data collection and refinement statistics are summarized in Table 2.

**Figure 6:**
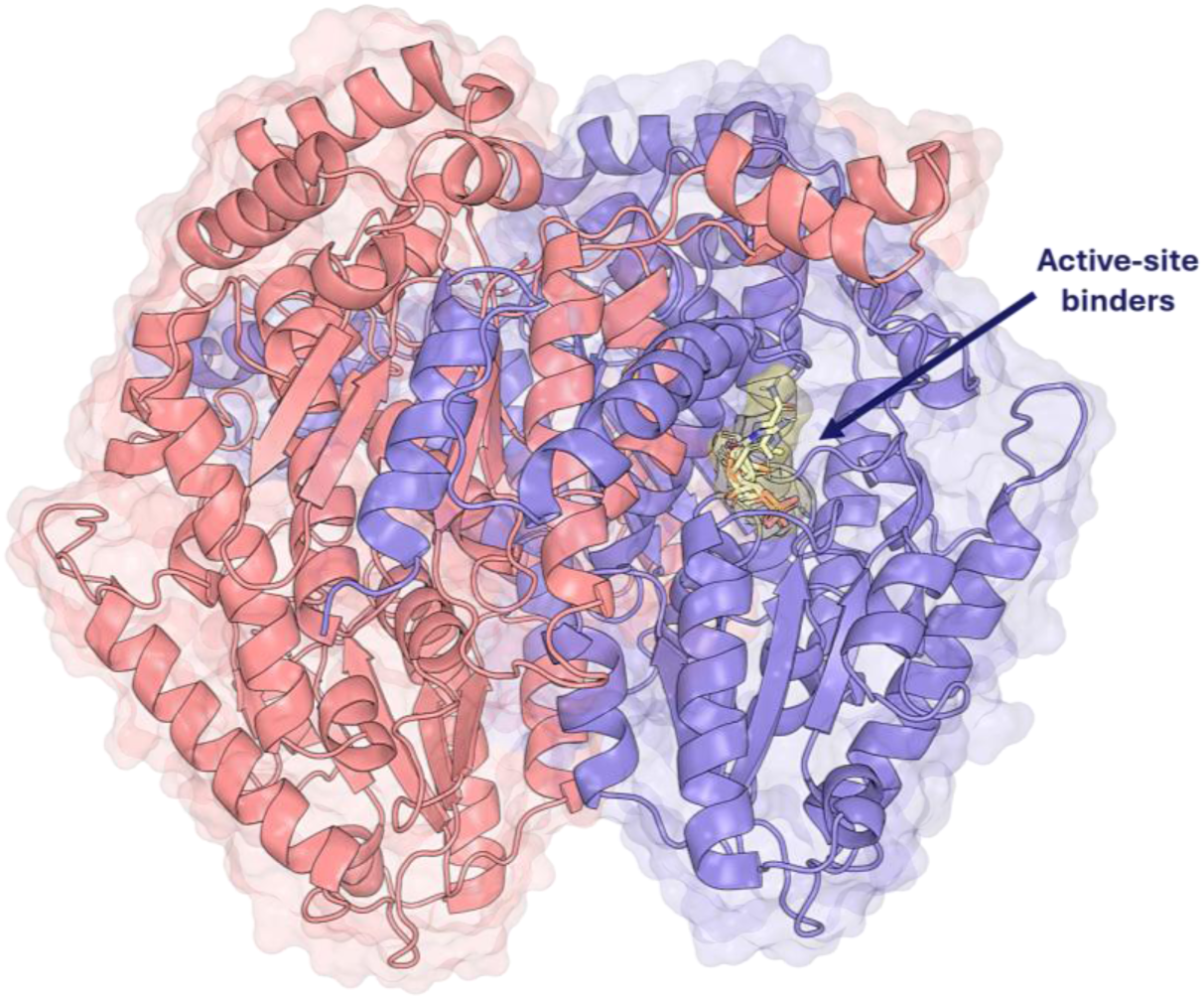
Overall view of the human GPI homodimer in complex with six active-site ligands identified by structure-based virtual screening (SBVS). Bound ligands (highlighted yellow) are shown in one of the subunits (purple), indicated by the arrow. The figure was generated with human GPI crystal structure (PDB: 32BF) using PyMOL. Structures of GPI-ligand complexes were obtained by co-crystallization; at 25 °C in a crystallization solution containing 22% (w/v) PEG 3350 and 58 mM HEPES (pH 7.0).

**Table 2:** X-ray diffraction data collection and refinement statistics for human GPI-ligand complexes.

| Structure | GPI-RMP<br>PDB: 32BF | GPI-CMP<br>PDB: 32JZ | GPI-GlcN6P<br>PDB: 32BE | GPI-NAG6P<br>PDB: 32JQ | GPI-DMAPP<br>PDB: 32XF | GPI-XMP<br>PDB: 32YR |
| --- | --- | --- | --- | --- | --- | --- |
| **Space group** | P 21 21 21 | P 21 21 21 | P 21 21 21 | P 21 21 21 | P 21 21 21 | P 21 21 21 |
| **Resolution range (Å)** | 49.39 – 1.66<br>(1.69 – 1.66) | 49.35 – 1.95<br>(1.98 – 1.95) | 49.60 – 2.14<br>(2.18 – 2.14) | 49.30 – 1.89<br>(1.92 – 1.89) | 48.36 – 2.14<br>(2.18 – 2.14) | 49.67 – 1.89<br>(1.92 – 1.89) |
| **Total reflections** | 3778051<br>(187942) | 2237462<br>(105648) | 1670223<br>(82123) | 2437041<br>(115343) | 1747693<br>(85891) | 2420090<br>(92354) |
| **Unique reflections** | 271513<br>(13077) | 161610<br>(7848) | 122656<br>(5983) | 176050<br>(8153) | 129618<br>(6378) | 178113<br>(8726) |
| **Multiplicity** | 13.9 (14.4) | 13.8 (13.5) | 13.6 (13.7) | 13.8 (14.1) | 13.5 (13.5) | 13.6 (10.6) |
| **Completeness (%)** | 98.8 (96.7) | 99.9 (99.0) | 100.0 (100.0) | 99.6 (93.6) | 100.0 (100.0) | 100.0 (100.0) |
| **Mean(I)/sig(I)** | 11.6 (1.4) | 8.1 (1.3) | 7.8 (2.0) | 9.2 (1.7) | 10.0 (2.4) | 11.3 (1.8) |
| **Wilson B-factor (Å<sup>2</sup>)** | 17.3 | 23.9 | 24.5 | 22.3 | 16.8 | 22.6 |
| **R<sub>merge</sub>** | 0.163 (2.524) | 0.250 (2.409) | 0.263 (1.667) | 0.208 (1.848) | 0.211 (1.163) | 0.146 (1.257) |
| **CC<sub>1/2</sub>** | 0.999 (0.676) | 0.998 (0.593) | 0.997 (0.837) | 0.998 (0.729) | 0.998 (0.755) | 0.999 (0.738) |
| **R<sub>work</sub>** | 0.186 | 0.187 | 0.189 | 0.191 | 0.215 | 0.166 |
| **R<sub>free</sub>** | 0.214 | 0.231 | 0.229 | 0.229 | 0.252 | 0.201 |
| **Ramachandran outliers (%)** | 0 | 0 | 0 | 0 | 0 | 0 |
| **Sidechain outliers (%)** | 0.1 | 0.1 | 0.3 | 0.2 | 0.2 | 0.1 |
| **RSRZ outliers (%)** | 1.8 | 1.1 | 1.6 | 2.7 | 2.7 | 0.9 |
| **Clash score** | 2.46 | 3.22 | 2.65 | 4.02 | 3.37 | 2.92 |

The binding poses of the ligands in complex with human GPI are depicted in Figure 7. Comparison of the docked and crystallographic ligand poses showed that GlcN6P was reproduced most accurately by SP Glide, with an RMSD of 2.4 Å, followed by NAG6P at 3.5 Å. In contrast, RMP, XMP, CMP, and DMAPP showed larger deviations, with RMSD values exceeding 4.7 Å (Suppl. Table S3). Their binding modes in the docked and crystal structures are shown in Suppl. Fig. S5 and S6, respectively. Although the overall RMSD values varied between ligands, the phosphoryl groups showed consistent alignment between the docked and crystallographic poses, maintaining a similar orientation and interactions within the active site. The main differences were observed in the backbone.

**Figure 7:**
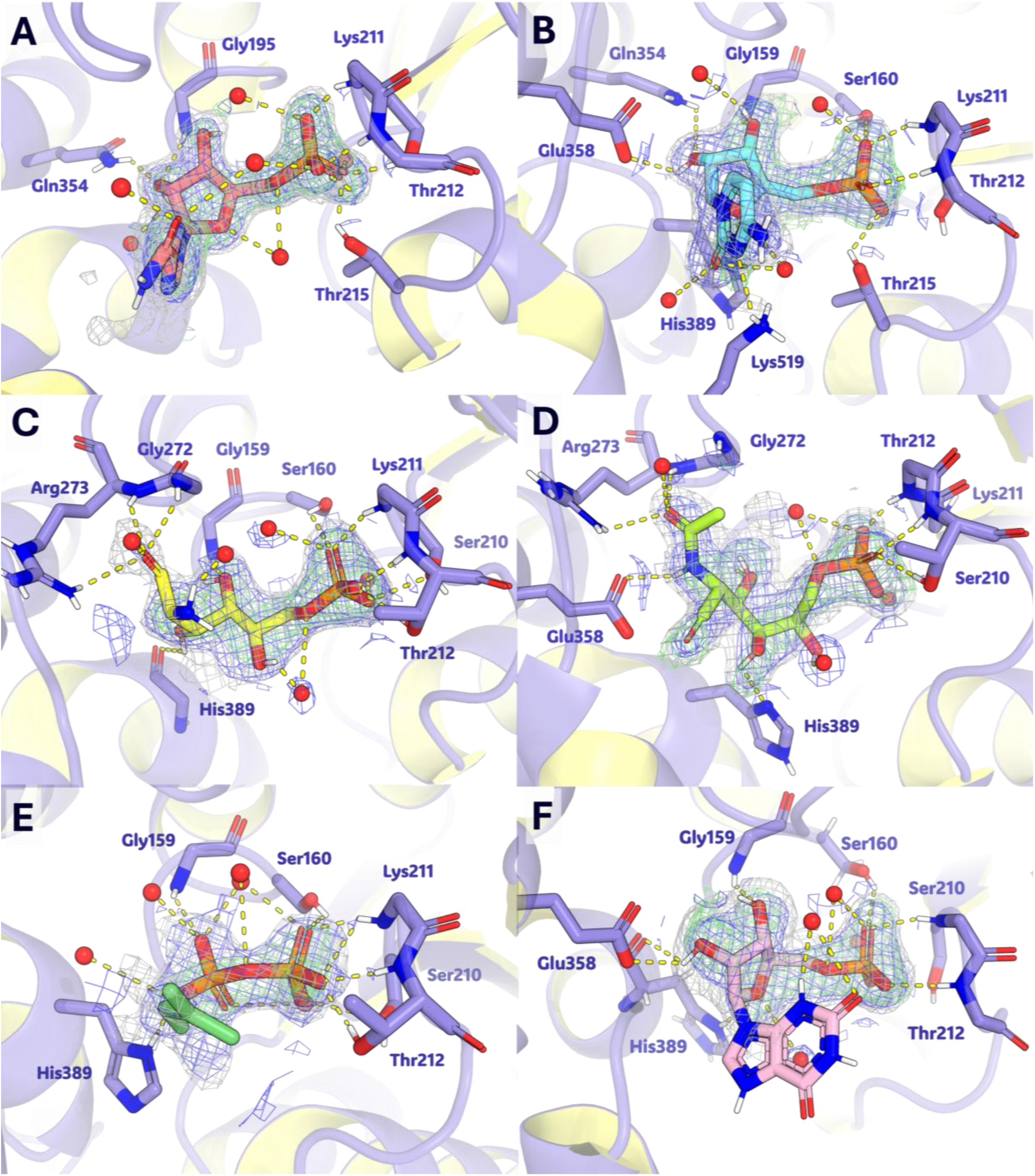
Close-up view of ligands bound in the active site of human GPI) isomerase. GPI-ligand co-crystals were grown at 25 °C in 22% (w/v) PEG 3350 and 58 mM HEPES (pH 7.0). (A) RMP in chain A; refined to 1.66 Å (PDB: 32BF). (B) CMP in chain D; refined to 1.95 Å (PDB: 32JZ). (C) GlcN6P in chain D; refined to 2.14 Å (PDB: 32BE). (D) NAG6P in chain C; refined to 1.89 Å (PDB: 32JQ). (E) DMAPP in chain C; refined to 2.14 Å (PDB: 32XF). (F) XMP in chain D; refined to 1.89 Å (PDB: 32YR). The 2mFo-DFc electron density maps are shown as blue mesh contoured at 1.0 σ, difference omit maps as green mesh at 3.0 σ, and polder maps as gray mesh at 3.0 σ. The yellow dashed lines indicate polar interactions between the ligands and GPI active-site residues. The figures were created using PyMOL.

The crystal structures of GPI-ligand complexes each contained two homodimers in the asymmetric unit (four monomers in total). In the GPI complexes with RMP (PDB: 32BF) and GlcN6P (PDB: 32BE), clear electron densities were observed in all four subunits. By contrast, the GPI-XMP complex (PDB: 32YR) showed electron density for the ribose phosphate moiety of XMP in one monomer of each dimer, whereas only small weak density (not modeled) corresponding to the phosphoryl group was found in the second monomer. In comparison, the complexes with CMP (PDB: 32JZ), NAG6P (PDB: 32JQ), and DMAPP (PDB: 32XF) showed ligand electron densities in three of the four monomers, whereas the remaining monomer had no visible ligand density. This occupancy pattern was observed across all three complexes, suggesting that the absence of ligand in one monomer might be due to crystal packing effects, differences in local crystal environments, or incomplete ligand occupancy rather than part-of-site/cooperative binding behavior. Notably, in one monomer of the GPI-NAG6P complex, the observed ligand density was best modeled by the closed-ring form of NAG6P (Suppl. Fig. S7) instead of the open-chain form seen in other two subunits (Fig. 7D). The cyclic conformation may arise from ring-chain equilibration of NAG6P in solution, followed by stabilization of the cyclic form upon binding to GPI.

Partial ligand occupancy of GPI subunits has been reported for weak inhibitors in an earlier study [16], in which maleate (MAE) displayed half-of-sites binding, occupying only two of the four subunits in the asymmetric unit, whereas phosphoenolpyruvate (PEP) exhibited asymmetric binding accompanied by local conformational change in the active-site loop (βc–α11). In the present study, subtle local conformational changes were observed in the α13 region adjacent to the active site, without evidence of half-of-sites or asymmetric binding (Suppl. Fig. S8). Nevertheless, similar to previous structural studies of human and other mammalian GPI in complexed with phosphorylated ligands, the current structural data showed that all ligands adopt highly conserved interaction patterns within the GPI active site. The ligand backbone is stabilized through hydrogen bonds with conserved active-site residues, including Gly159, Arg273, Glu358, and His389. Despite structural differences among the ligands, the phosphoryl group is consistently anchored by a conserved network of hydrogen bonds and electrostatic interactions involving Ser160, Lys211, Thr212, and Thr215. The strong preference for dianionic groups with phosphate-like geometry highlights the importance of charge complementarity within the active site, providing a rationale for the poor binding of compounds lacking these features.

### Identified GPI inhibitors partially reduced HEK-293 and U2OS glycolytic flux

To investigate whether the biochemical activity of the identified ligands translated into *in vitro* cell culture assays, three compounds – RMP, XMP, and GlcN6P were evaluated for their effects on cell glycolytic flux and viability. In addition, the prodrug form of RMP – ribavirin was included for comparison with its active metabolite. Two glycolytic inhibitors – 2-deoxyglucose (2-DG) and 6-phosphogluconate (6PG) were utilized as control compounds, whereas the mitochondrial inhibitors – rotenone and antimycin A (Rot/AA) were employed to examine the dual inhibition of glycolysis and oxidative phosphorylation (OXPHOS). The compounds were tested in two human cell lines – human embryonic kidney (HEK-293) and human osteosarcoma (U2OS) cells. The glycolytic flux of both cell lines was measured using the Seahorse XFe96 Flux Analyzer (Agilent Technologies, USA). The cells were treated with the selected compounds during the flux measurement, and glycolytic rate was quantified as the proton efflux rate (PER), calculated from real-time extracellular acidification rate (ECAR) measurements after correcting for the contribution of mitochondrial respiration.

The glycolytic PER (glycoPER) of HEK-293 and U2OS cells, measured after test compound and Rot/AA injection, are shown in Figures 8 and 9. A clear concentration-dependent decrease in glycoPER was observed in HEK-293 cells treated with XMP and GlcN6P, suggesting inhibition of glycolytic flux. The inhibition remained partial, with glycoPER reduced by only approximately 50% at the highest tested concentration of GlcN6P (20 mM). RMP also reduced glycoPER at 5 mM; however, no clear dose-dependent effect was observed over the tested concentration range (1–5 mM). In contrast, ribavirin, the non-phosphorylated prodrug, showed almost no effect on glycoPER at 5 mM and only minimal reduction at 10 mM. The effects of these compounds were further examined in U2OS cells to assess whether the glycolytic response was consistent across different cell types. All tested compounds led to decrease of glycoPER in U2OS cells, with greatest reduction reaching approximately 40–50% at the highest tested concentrations. Among the compounds, XMP, RMP, and ribavirin displayed an apparent concentration-dependent decrease in glycoPER. Compared to ribavirin, RMP showed stronger inhibitory effects, reducing glycoPER by 40% at 5 mM in U2OS, whereas ribavirin required higher concentration to elicit similar effects, indicating that the phosphorylated form may exhibit greater inhibition on glycolytic flux than its non-phosphorylated counterpart. GlcN6P also significantly reduced glycoPER in U2OS cells, although the dose-dependent effect was less evident compared with XMP and RMP. Similar to HEK-293 cells, the glycoPER reduction in U2OS remained partial, indicating incomplete inhibition of glycolytic activity.

**Figure 8:**
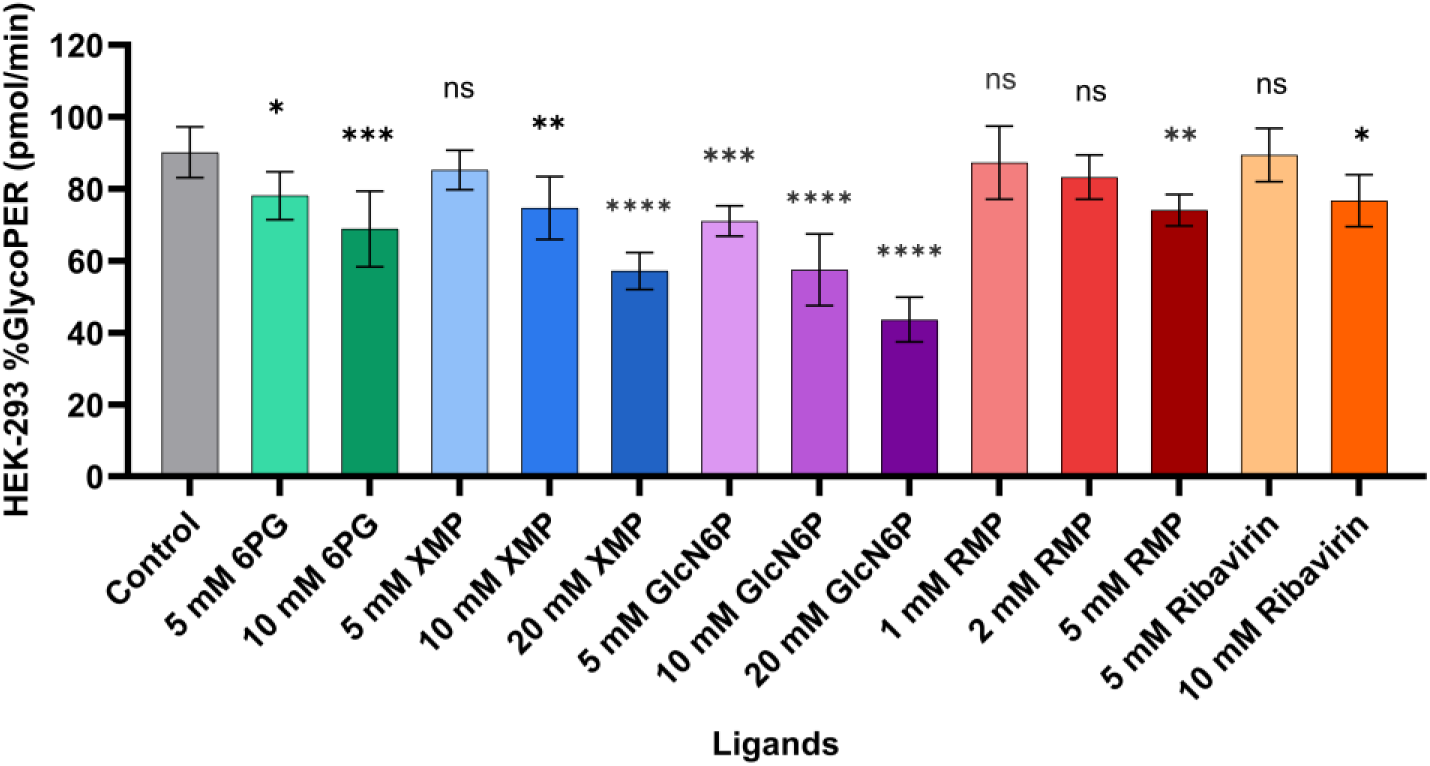
Percentage (%) glycolytic proton efflux rate (glycoPER) of HEK-293 cells after treatment with the indicated compounds: XF-assay medium as control; 6-phosphogluconate (6PG) as positive control, XMP, GlcN6P, RMP, and ribavirin. GlycoPER was measured following effector compound and rotenone/antimycin A (Rot/AA) injections and normalized to the basal glycoPER measured before compound injection. Bars represent the mean ± SD of five replicates (n = 5) per condition. Statistical P-values: ns (P ≥ 0.05), * (P = 0.01–0.05), ** (P = 0.001–0.01), *** (P = 0.0001–0.001), and **** (P < 0.0001), were determined by one-way ANOVA using GraphPad Prism.

**Figure 9:**
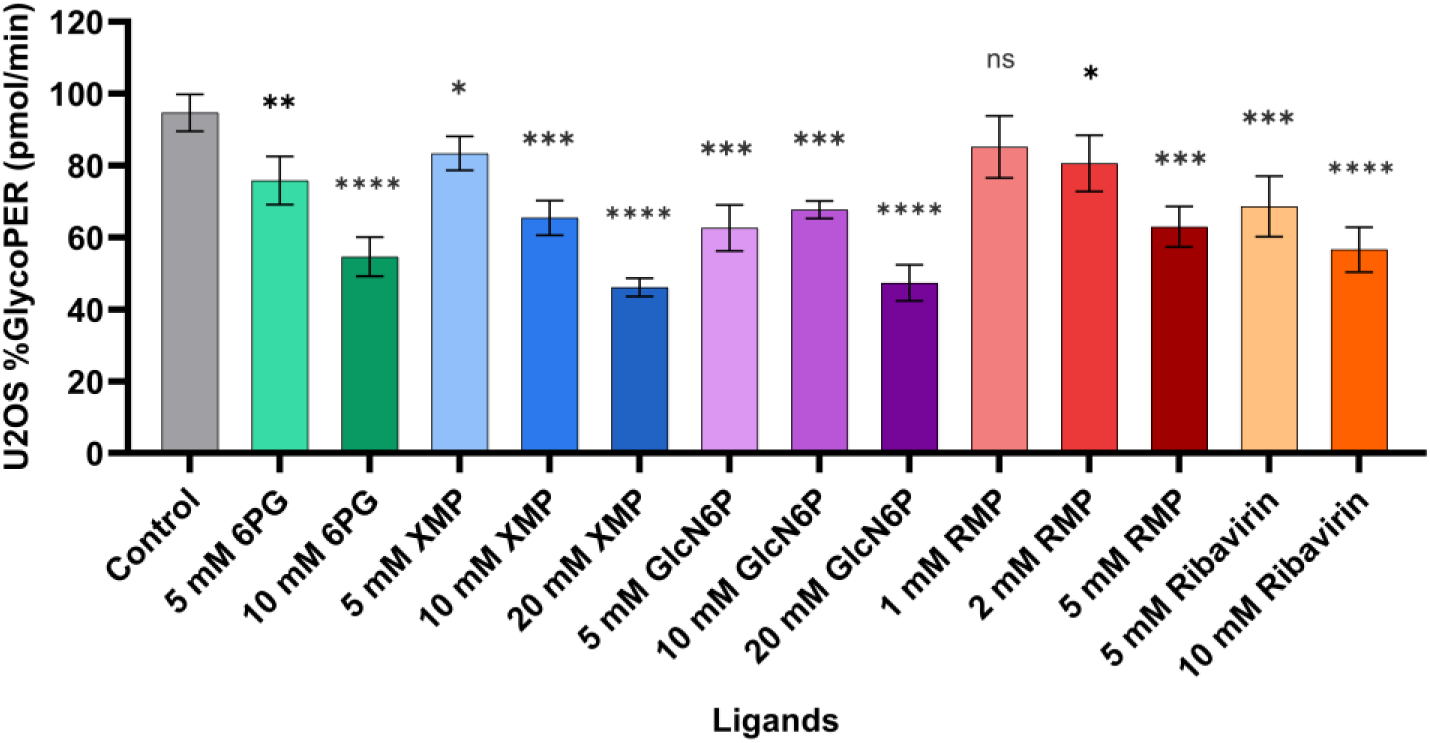
Percentage (%) glycolytic proton efflux rate (glycoPER) of U2OS cells following treatment with the indicated test compounds: XF-assay medium as control; 6-phosphogluconate (6PG) as positive control, XMP, GlcN6P, RMP, and ribavirin. GlycoPER was measured after effector compound and rotenone/antimycin A (Rot/AA) injections and normalized to the basal glycoPER measured before compound injection. Bars represent the mean ± SD of five replicates (n = 5) per condition. Statistical P-values: ns (P ≥ 0.05), * (P = 0.01–0.05), ** (P = 0.001–0.01), *** (P = 0.0001–0.001), and **** (P < 0.0001), were determined by one-way ANOVA using GraphPad Prism.

### GPI inhibitors have limited effects on HEK293 and U2OS cell growth

To determine whether the selected compounds affected cell growth, the viability of both HEK-293 and U2OS cells was assessed before and after 24 h, 48 h, and 72 h of treatment using PrestoBlue assay. The glycolysis inhibitor 2-DG and the mitochondrial inhibitors Rot/AA were included for comparison. The viability of HEK-293 and U2OS cells measured after 72 h of compound treatment are shown in figures 10 and 11. After the 72-h treatment, XMP and GlcN6P had minimal effects on the viability of both HEK-293 and U2OS cells. In contrast, both RMP and ribavirin resulted in dose-dependent decrease in viability, with significantly greater inhibitory effects observed for ribavirin at equivalent concentrations. Ribavirin reduced U2OS viability to approximately 60%, 35%, and below 20% at 2 mM, 5 mM, and 10 mM, respectively, while the corresponding reductions in HEK-293 cells were slightly less pronounced. Overall, the observed glycoPER decrease did not translate into reduced cell viability. Both XMP and GlcN6P significantly reduced glycoPER but had minimal impact on cell growth, suggesting that partial glycolytic inhibition by these metabolites was insufficient to impair long-term cell growth. On the contrary, ribavirin and RMP exhibited greater cytotoxic effects despite weak inhibition on the glycolytic flux.

**Figure 10:**
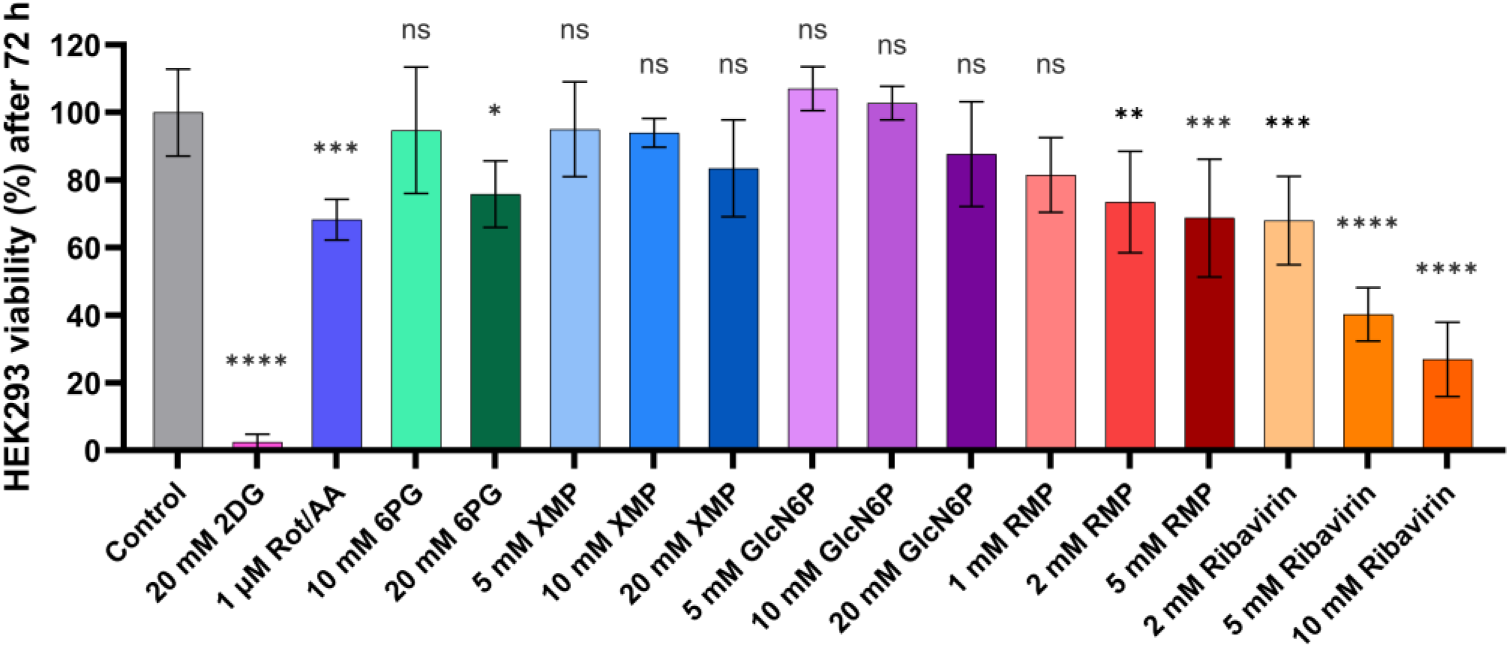
Percentage (%) cell viability of HEK-293 after 72-hour compound treatment. Relative viability of HEK-293 cells following 72 hours of treatment with the indicated compounds: 2-DG, Rot/AA, 6PG, XMP, GlcN6P, RMP, and ribavirin. Cell viability was determined using the PrestoBlue assay and normalized to the pretreatment values for each condition. Bars represent the mean ± SD of six replicates (n = 6) per condition. Statistical significance was determined by one-way ANOVA using GraphPad Prism. ns, P ≥ 0.05; *P < 0.05; **P < 0.01; ***P < 0.001; ****P < 0.0001

**Figure 11:**
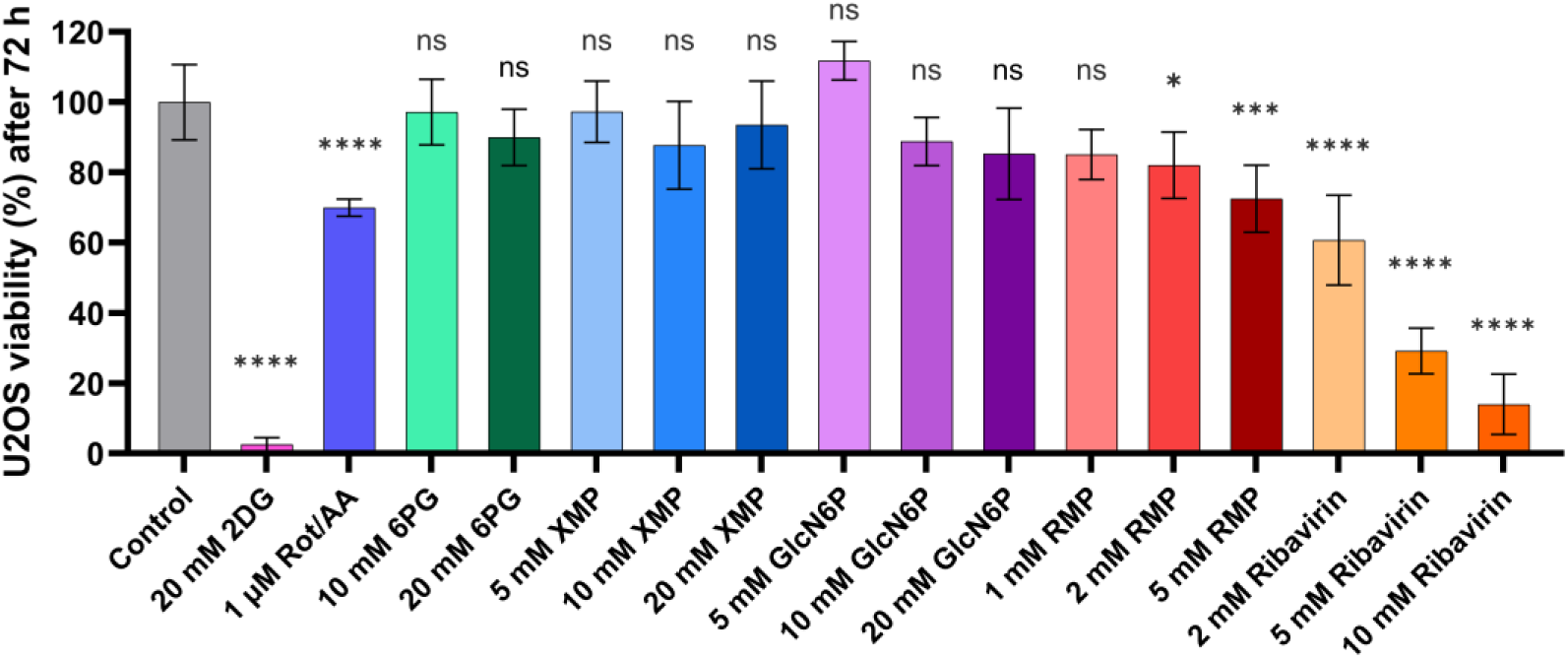
Percentage (%) cell viability of U2OS after 72-hour compound treatment. Relative viability of U2OS cells after 72 hours of treatment with test compounds: 2-DG, Rot/AA, 6PG, XMP, GlcN6P, RMP, and ribavirin. Cell viability was determined using the PrestoBlue assay and normalized to the pretreatment values for each condition. Bars represent the mean ± SD of six replicates (n = 6) per condition. Statistical significance was determined by one-way ANOVA using GraphPad Prism. ns, P ≥ 0.05; *P < 0.05; **P < 0.01; ***P < 0.001; ****P < 0.0001

To examine if dual inhibition of glycolysis and mitochondrial respiration further impaired cell growth, HEK-293 and U2OS cells were treated with 1 µM Rot/AA alone or in combination with GlcN6P or ribavirin, and 2-DG was used as positive control. The viability of HEK-293 and U2OS cells was monitored over 72 h (Fig. 12). In both cell lines, untreated control cells showed continuous proliferation throughout the experiment. As expected, combined treatment of 2-DG and Rot/AA almost completely abolished the cell growth, resulting in minimal cell viability after 72 h. Rot/AA alone slightly inhibited cell growth in both HEK-293 and U2OS cells, reducing the final viability to approximately 75–80% of the untreated control. Addition of GlcN6P to Rot/AA showed almost no difference compared with Rot/AA alone, and the growth curves remained largely similar throughout the experiment. In contrast, combining ribavirin and Rot/AA resulted in a substantially greater suppression of cell growth than Rot/AA alone in both cell lines. The viability decrease became more pronounced over time and was clearly evident after 48 hours. At the 72-hour endpoint, the Rot/AA-ribavirin combination reduced cell viability to approximately 30–40% of the untreated control, representing a significantly greater reduction than Rot/AA alone. However, compared with ribavirin alone, the Rot/AA-ribavirin combination only led to a slightly greater decrease (by 3.5%) in viability in U2OS cells but negligible difference (∼1.5%) in HEK-293 cells. This suggested that the combination did not synergistically reduced cell viability.

**Figure 12:**
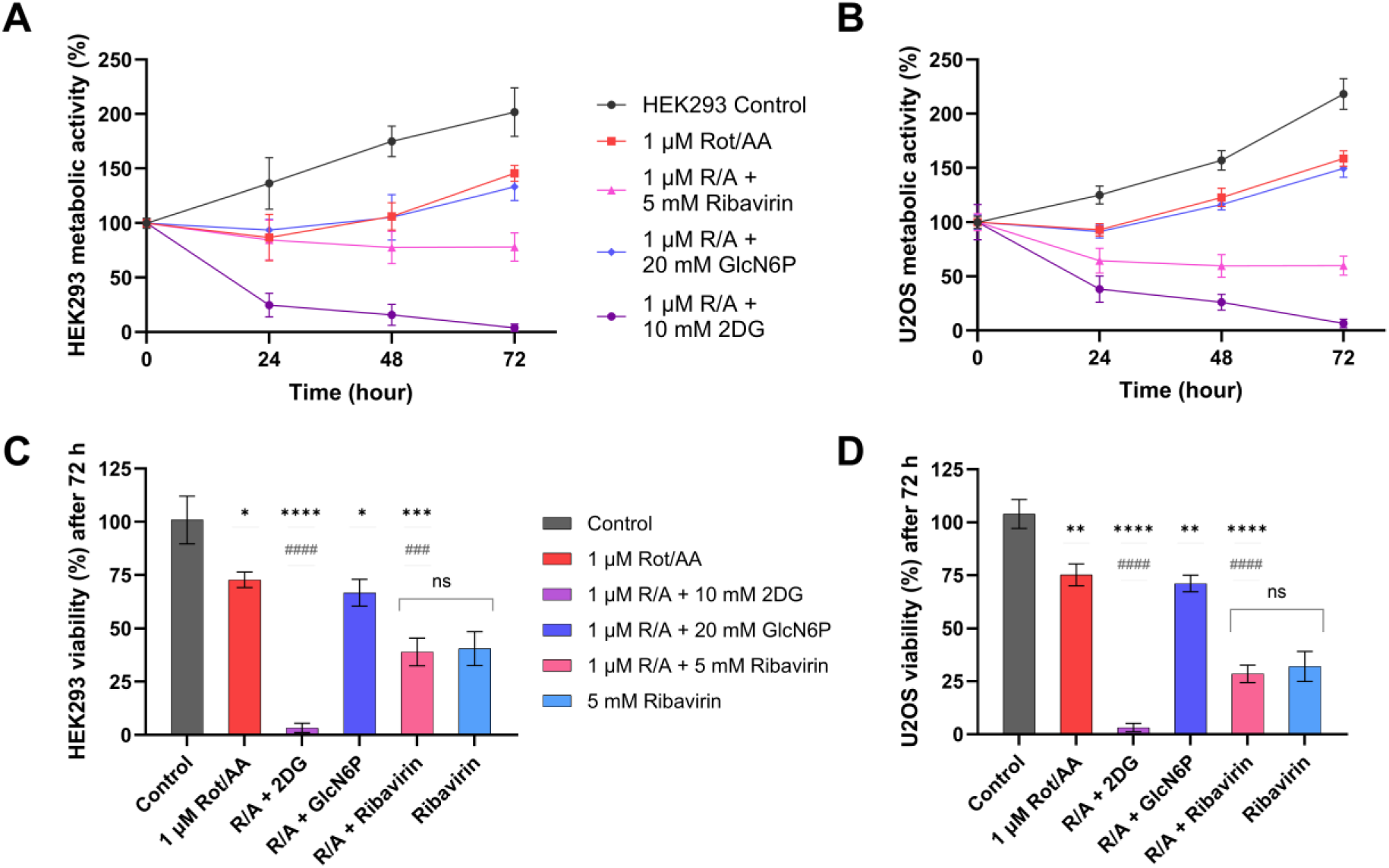
Combined effects of glycolytic and mitochondrial inhibition on the viability of HEK-293 and U2OS cells. Relative metabolic activity of HEK-293 (A) and U2OS (B) cells measured at 0, 24, 48, and 72 hours following treatment with Rot/AA (R/A) alone or in combination with ribavirin, GlcN6P, or 2-DG. The lower panels summarize the relative cell viability of HEK-293 (C) and U2OS (D) after 72 h of treatment with the indicated compounds, including ribavirin alone. Cell viability was determined using the PrestoBlue assay and normalized to the pretreatment values at each time point. Data represent the mean ± SD of six replicates (n = 6) per condition. Statistical significance was determined by one-way ANOVA using GraphPad Prism: Comparisons with the control are indicated as: ns (P ≥ 0.05); * (P < 0.05); **(P < 0.01); ***(P < 0.001); ****(P < 0.0001). Comparisons with the R/A-treated group are indicated as: ^##^(P < 0.01); ^###^(P < 0.001); ^####^(P < 0.0001). Brackets indicate comparisons between the Rot/AA + ribavirin and ribavirin treatment groups: ns (P ≥ 0.05).

**Figure 13:**
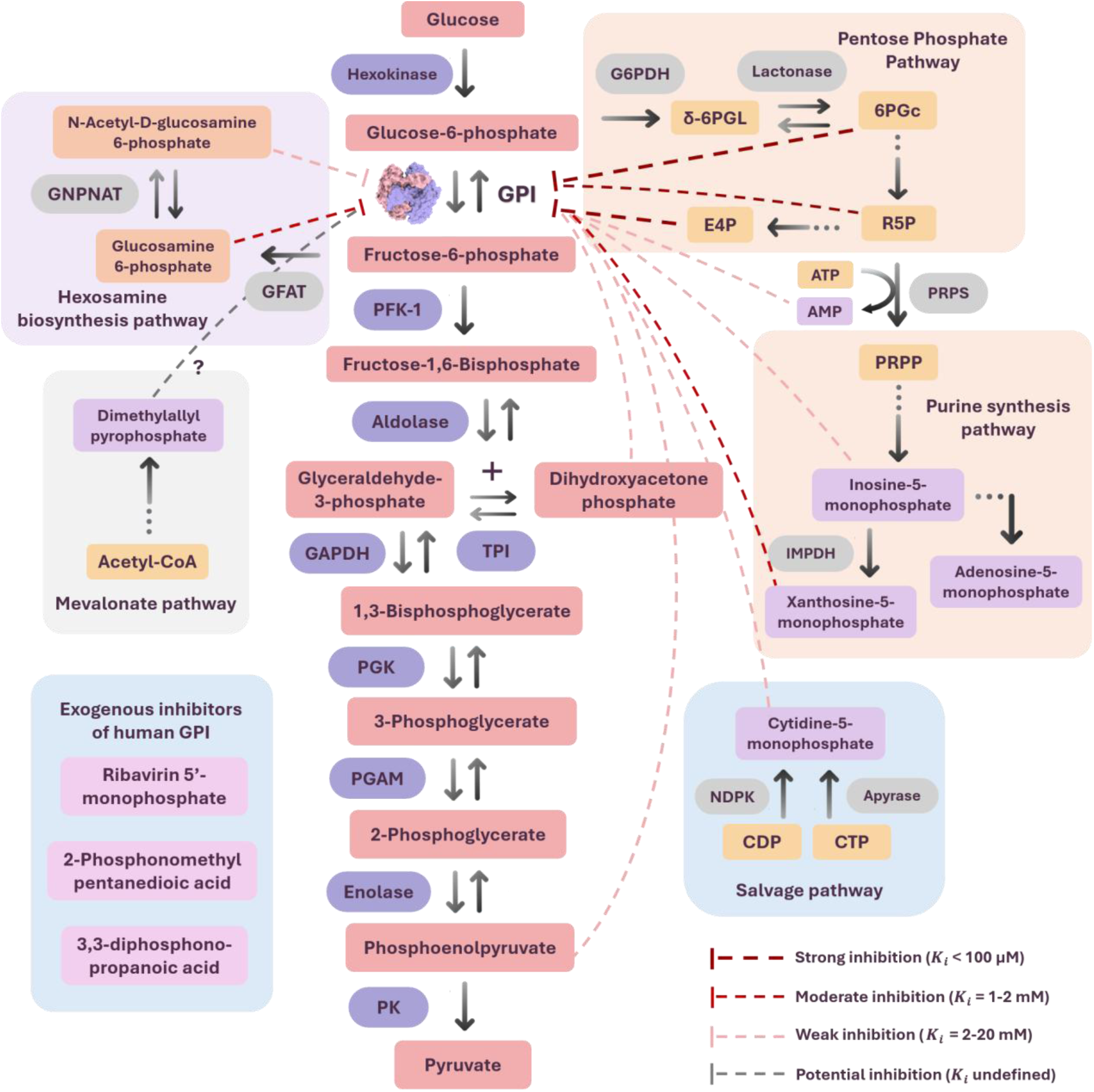
Representative cytosolic metabolites and exogenous compounds that inhibit human glucose-6-phosphate isomerase (GPI). Enzyme abbreviations: Glucosamine-phosphate N-acetyltransferase (GNAPNAT); Fructose-6-phosphate amidotransferase (GFAT); Phosphofructokinase-1 (PFK-1); Triose phosphate isomerase (TPI); Glyceraldehyde-3-phosphate dehydrogenase (GAPDH); Phosphoglucokinase (PGK); Phosphoglycerate mutase (PGAM); Pyruvate kinase (PK); Glucose-6-phosphate dehydrogenase (G6PDH); Phosphoribosyl pyrophosphate synthetase (PRPS); Inosine 5’-monophosphate dehydrogenase (IMPDH); Nucleoside-diphosphate kinase (NDPK). Metabolite abbreviations: δ-6-phosphogluconolactone (δ-6PGL); Phosphoribosyl pyrophosphate (PRPP); Cytidine diphosphate (CDP); Cytidine triphosphate (CTP).

## Discussion

### Virtual screening identified 11 GPI modulators and additional putative binders

The docking of 1.1 million compounds using Schrodinger Glide yielded 1480 putative binders. After re-scoring by Prime MM-GBSA, 193 compounds were retained for further curation, and 32 of them were experimentally validated. Among the tested, 11 compounds were confirmed to be true GPI modulators, whereas the remaining were false positives. All the confirmed hits were hydrophilic compounds, containing scaffolds such as nucleoside or sugar chain attached to a phosphoryl or phosphate-like group, while the false-positives are mostly aromatic, with a few being phosphate-based. Of the untested candidates, 138 share structural features with the previously reported GPI inhibitors, representing putative binders for further evaluation. These compounds fall into three major groups: 1) phosphorylated carbohydrates and sugar derivatives (39), 2) mono-and diphosphorylated nucleotides (48), and 3) compounds containing phosphate mimetic groups (51). Ligand interaction diagrams of representative compounds from the 138 candidates revealed a high degree of similarity to known GPI ligands in their predicted binding modes (Suppl. Fig. S9), interacting with active-site residues via conserved H-bonding and electrostatic interactions.

### Structural features underlying effective GPI binding and inhibition

SiteMap (Schrodinger) analysis predicted the GPI active site to be potentially druggable, with a large pocket volume (374 Å³) and a SiteScore of 1.14. The pocket can indeed accommodate ligands spanning broad molecular weight range (114 to 520 Da), as demonstrated in this study and previous report [16]. Nevertheless, it is dominated by conserved polar, charged residues, making it difficult to target with drug-like scaffolds. The enzyme has evolved to recognize its natural substrates and stabilize the high-energy, enediolate-like transition state formed during the isomerization. Throughout catalysis, the substrate undergoes structural changes, involving conversion from an open-ring G6P (aldose form), to a partially planar cis-enediol intermediate, and an open-ring F6P (ketose form) [13]. The highest-energy state is generally thought to be enediolate-like, either closely resembling the cis-enediolate intermediate or adopting a structure that lies between the open-chain sugar and the cis-enediolate. The most potent inhibitors of human GPI reported to date are structural mimics of the cis-enediolate intermediate, exhibiting potency in the lower micromolar range (K_i_ = 0.1-20 µM). The synthesis of GPI transition-state analogs has been described by Achmad et al. (2020) [19]. Other strong inhibitors of human GPI, such as arabinose 5-phosphate (A5P; K_i_ = 3 µM) [19,20], and mammalian GPI inhibitors, including 6PG (K_i_ = 42 µM) [21] and erythrose-4-phosphate (E4P; K_i_ = 0.7 µM) [22]; are structurally related to the open-chain forms of GPI substrates and also exhibit sub-micromolar inhibitory potency. In this study, the K_i_ of 6PG was estimated at ∼106 µM (Suppl. Fig. S10); about 12-fold higher than that reported for GPI from human erythrocytes (K_i_ = 9 µM), likely due to differences in assay conditions. The previous study used pH 8.0, 37 °C, and 0.03-1 mM F6P, while our assay was conducted at pH 7.4 and 25 °C, with 0.05-4 mM F6P.

The inhibitors identified in this study were relatively weak inhibitors, with R5P showing the strongest inhibitory activity (K_i_ = ∼0.6 mM). R5P predominantly exists in its cyclic form in solution (> 99.5%) [23,24], and therefore, the measured K_i_ should reflect the inhibitory potency of the cyclic conformer. RMP and nucleotide monophosphate (NMP) inhibitors are based on the cyclic ribose-phosphate scaffold, which likely serves as the core binding moiety. This was particularly evident from crystallographic structures of GPI in complex XMP, IMP, and AMP, in which electron density was observed only for the ribose-phosphate moiety, suggesting that the nucleobases were poorly coordinated within the enzyme active site. Moreover, the presence of triazole in RMP and nucleobase in the NMPs did not enhance inhibitory activity, and even led to reduced inhibition as seen in CMP and AMP. This suggested that the heterocyclic substituents contribute little to binding affinity, or could introduce unfavorable contacts and steric clashes that impair the binding. Likewise, despite sharing high structural similarity with 6PG, GlcN6P and NAG6P exhibited approximately 20-and 40-fold weaker inhibitory activity than 6PG, respectively. All three compounds contain a six-carbon sugar-phosphate scaffold, but they differ in the substituents at the C1 and C2 positions. In 6PG, the C2 hydroxyl group and C1 carboxylate can form multiple H-bonding and electrostatic interactions with the active-site residues (Suppl. Fig. S11). In comparison, GlcN6P contains an amino group at C2 and an aldehyde at C1, whereas NAG6P carries an N-acetyl group at C2. These substituents can alter the charge distribution, H-bonding patterns, and steric properties of the ligands, thereby limiting their ability to reproduce the interactions made by 6PG.

Central to GPI ligand recognition is the phosphate-binding pocket, where the dianionic phosphoryl group is coordinated by residues including Lys211, Thr212, and Thr215. These interactions anchor the ligand, orient the sugar moiety for catalysis, and contribute substantially to binding affinity. The importance of the phosphoryl group in GPI inhibition has been highlighted in numerous studies. Inorganic phosphate alone can act as a weak GPI inhibitor with K_i_ of 1.7 mM, whereas fructose and glucose have shown no detectable GPI inhibition [25,26]. Kinetic study on *Trypanosoma brucei* GPI demonstrated that the removal of phosphoryl group from potent inhibitors such as D-arabinonhydroxamic acid-5-phosphate (5PAH; IC_50_ = 0.4 µM) and D-arabinoate-5-phosphate (5PAA; IC_50_ = 0.9 µM) reduced their inhibitory activity by 10,000–15,000-fold, resulting in IC_50_ values of 6.0 mM and 9.5 mM, respectively [27]. Similarly, in our studies, glucosamine, ribose, and ribavirin, all of which lack a phosphoryl group, showed no apparent inhibitory activity against human GPI over the tested concentration range of 0.5–24 mM (data not shown).

Beyond the above, stereochemical factors may also influence GPI inhibition, particularly for linear phosphorylated sugars. This was exemplified by D-xylulose-5-phosphate (Xu5P), which is structurally related to the potent GPI inhibitor E4P, but showed no apparent GPI inhibition nor binding when tested over a broad concentration range (1 µM–40 mM). Zig-zag and Fischer projections of the linear phosphorylated sugar GPI inhibitors and Xu5P revealed differences in their relative configuration of the two hydroxyls closest to the phosphoryl group (Suppl. Table S4). In the zig-zag representation, the two hydroxyl groups of GPI inhibitors consistently point in opposite directions, whereas those of Xu5P are oriented in the same direction. Conversely, in the Fischer projections, the two hydroxyl groups are positioned on the same side in the GPI inhibitors but on opposite sides in Xu5P. Given its presence among the phosphorylated linear sugar inhibitors but absence in Xu5P, this relative configuration of the hydroxyl groups likely contributes to GPI recognition and inhibition.

### Bioisosteres as potential phosphate replacements

Phosphate bioisosteres are valuable tools in ligand design, offering a means to replace the phosphoryl group while preserving its key binding interactions. This is particularly relevant for enzymes like GPI, where the phosphoryl group is critical for ligand recognition. In this study, two phosphate bioisosteres, MDP and PAE, were shown to have binding affinities similar to that of inorganic phosphate. Furthermore, the PAE-and MDP-based compounds, 2PMPA and 33DP, exhibited weak inhibition of GPI activity. Building on this proof of concept, phosphoryl groups in the potent transition-state analogues [19] were replaced with PAE or MDP, followed by evaluation with SP Glide docking and Prime MM-GBSA analysis. The docking poses showed that both MDP-and PAE-substituted analogues could potentially retain the key interactions observed for the parent inhibitors, including conserved contacts within the phosphate-binding region. Moreover, their Glide and MM-GBSA scores were within the range predicted for the potent inhibitors (Suppl. Table S5), further supporting the potential of MDP and PAE as phosphate replacements for GPI ligand optimization.

### GPI interacts with metabolites across multiple pathways in the cytosol

The strongest endogenous GPI inhibitors reported so far, E4P and 6PG, are intermediates of the pentose phosphate pathway, which occurs within the cytosol. In a previous study, we reported several weak metabolite inhibitors of human GPI, including dihydroxyacetone phosphate (DHAP) and phosphoenolpyruvate (PEP), both of which are cytosolic metabolites derived from glycolysis [16]. The metabolite inhibitors identified in this study also represent intermediates from several cytosolic metabolic pathways (Fig. 12). R5P belongs to the same pathway as E4P and 6PG, and serves as a precursor for nucleotide biosynthesis. IMP and XMP are central intermediates in *de novo* purine biosynthesis, where they are converted into adenine and guanine nucleotides [28]. Likewise, AMP and CMP are continuously interconverted within the cellular adenine and pyrimidine nucleotide pools, depending on the cellular energy and nucleotide status. On the other hand, GlcN6P and NAG6P are intermediates of the hexosamine biosynthetic pathway that ultimately give rise to UDP-GlcNAc – a key precursor for protein and lipid glycosylation [29]. In addition, DMAPP is an intermediate of the mevalonate/isoprenoid biosynthesis pathway and serves as a precursor for isoprenoids, sterols, and protein prenylation [30].

Whether such weak interactions have physiological significance under normal cellular conditions remains unclear. However, the concentrations of individual metabolites can fluctuate substantially in response to altered metabolic demand, nutrient availability, or genetic perturbations. Under pathological conditions, mutations in metabolic enzymes or their dysregulation can lead to the accumulation or depletion of pathway intermediates, potentially increasing the likelihood of weak metabolite–enzyme interactions. For instance, activity impairment of ribose-5-phosphate isomerase (RPI) [31] and transketolase (TKT) [32] has been shown to cause R5P accumulation. Moreover, mutations in enzymes of purine synthesis, such as superactivity of phosphoribosyl pyrophosphate synthetase (PRPS) can increase the intracellular pools of IMP, XMP, and AMP [33], whereas deficiency of inosine-5’-monophosphate dehydrogenase (IMPDH) can lead to accumulation of IMP, and downregulation of guanine monophosphate synthetase (GMPS) may result in elevated levels of XMP [34]. Furthermore, overexpression of uridine/cytidine kinase (UCK) in pyrimidine salvage pathway may enhance intracellular CMP levels [35]. Similarly, perturbations of the hexosamine biosynthetic pathway may affect the concentrations of GlcN6P and NAG6P. While gain-of-function mutation in glutamine-fructose-6-phosphate transamidase (GFPT1) may cause GlcN6P to accumulate [36], upregulation of glucosamine-phosphate N-acetyltransferase 1 (GNPNAT1) [37] and/or impairment of phosphoglucomutase 3 (PGM3) can lead to increased NAG6P levels [38]. Under such conditions, transient interactions with GPI that are negligible under physiological metabolite concentrations could become more pronounced and potentially influence glycolytic activity.

### GPI inhibitors caused partial glycolytic inhibition in HEK-293 and U2OS cells

Glycolytic flux assay was performed to determine whether the confirmed GPI inhibitors suppress glycolytic flux in intact cells. RMP, ribavirin, XMP, and GlcN6P were assessed for their effects on cellular glycolytic flux. The four compounds reduced glycoPER to varying extents in both HEK-293 and U2OS cells, supporting their ability to interfere with cellular glycolysis. The observed glycolytic inhibition remained partial in both cell lines, with greatest glycoPER decrease of 40–50%. Interestingly, the glycoPER decrease was more pronounced in U2OS cells than in HEK-293 cells. While HEK-293 cells are generally considered more glycolytic than U2OS cells, the former appeared less sensitive to the glycolytic inhibitors under the tested conditions. One possible explanation is that the greater glycolytic capacity of HEK-293 cells, reflected by their higher basal and glycolytic ECAR levels (Suppl. Fig S12 & S13), allowed them to better maintain their glycolytic activity. Moreover, RMP and its non-phosphorylated form – ribavirin exhibited different effects on glycolytic flux. RMP consistently led to greater glycoPER decrease than ribavirin at comparable concentrations in both cell lines. This difference may reflect the need for intracellular phosphorylation of ribavirin before it can exert inhibitory activity on GPI.

### The metabolic fate of GPI inhibitors influence their long-term cellular effects

Glycolytic inhibition by the endogenous metabolites did not translate into substantial suppression of cell growth. The cell viability studies showed that RMP and ribavirin had greater cytotoxicity than the endogenous metabolites (XMP, GlcN6P and 6PG), suggesting differences in their metabolic fates that underly their effects on cell growth. In addition, there might be mechanisms beyond GPI inhibition contributed to the cytotoxic effects of ribavirin and RMP. Ribavirin is a synthetic guanosine analogue that enters cells mainly through equilibrative nucleoside transporters (ENTs), particularly ENT1 [39]. After entering cells, ribavirin may undergo sequential phosphorylation, becoming mono-, di-, or tri-phosphorylated. Ribavirin and its active metabolites can interact with multiple cellular targets to suppress cell growth. RMP, for instance, is well-known as a potent competitive inhibitor of human IMPDH – a rate limiting enzyme in guanine nucleotide biosynthesis [40]. Inhibition of IMPDH can suppress *de novo* guanine nucleotide biosynthesis and limiting nucleotide availability required for rapidly proliferating cells [40]. In contrast to RMP, the other three GPI inhibitors, GlcN6P, XMP, and 6PG, had only minimal effects on cell growth during the 72-h treatment period, despite showing moderate inhibition of glycolytic flux in HEK-293 and U2OS cells. These findings suggest that the glycolytic inhibition achieved by these metabolites are insufficient to impair cell proliferation. This is not unexpected, as all three compounds are endogenous metabolic intermediates that can be further metabolized intracellularly, potentially reducing their effective concentrations over time.

### Combining selected glycolytic inhibitors with mitochondrial inhibition did not further suppress cell growth

No substantial synergistic effects on cell growth were observed for the combination of Rot/AA and selected inhibitors. To examine if dual inhibition of glycolysis and mitochondrial respiration would lead to greater growth suppression, ribavirin and GlcN6P were combined with Rot/AA. While Rot/AA-ribavirin combination reduced cell viability more than Rot/AA alone, Rot/AA-GlcN6P showed no difference to the Rot/AA treatment. Moreover, compared with ribavirin alone, the Rot/AA–ribavirin combination produced only a slightly greater decrease (1.5-3.5%) in viability of HEK-293 and U2OS cells. These findings suggested that simultaneous inhibition of glycolytic and mitochondrial activities with the compound combinations did not produce additive or synergistic effect under the conditions tested.

### Multitarget mechanisms, metabolic activation, and toxicity of ribavirin

The pronounced inhibitory effects of ribavirin and RMP on HEK-293 and U2OS cell growth, despite their weak glycolytic inhibition, suggested that additional mechanisms contributed to their cytotoxic effects. Ribavirin is a broad-spectrum antiviral medication used in the treatment of many viral infections, including respiratory virus infections [41]. Beyond its established antiviral applications, ribavirin has attracted considerable interest as a candidate for anticancer drug repurposing, with preclinical studies and clinical trials showing antitumor effects across multiple malignancies, particularly acute myeloid leukemia [42]. Besides the cytotoxic activity of its phosphorylated metabolites, ribavirin itself has been demonstrated to inhibit proliferation of sarcoma cell lines, including U2OS, by downregulating protein arginine methyltransferases (PRMTs) 1 and 5 [43]. Moreover, ribavirin directly inhibits the oncogenic translation initiation factor eIF4E by competing with the mRNA cap structure, leading to reduced translation of growth-promoting transcripts such as MYC, cyclin D1, VEGF, and BCL2 [44]. Ribavirin treatment has also been shown to reduce EZH2 expression and histone methyltransferase activity in several cancer models, although current evidence suggested that these effects are likely indirect or cell type-dependent rather than arising from direct inhibition of EZH2 [44,45].

Despite its established clinical application, ribavirin treatment is associated with a well-characterized adverse effect, hemolytic anemia, which has been linked to the intracellular accumulation of phosphorylated ribavirin metabolites in erythrocytes [46]. This accumulation has been associated with reduced erythrocyte ATP levels, oxidative stress, membrane damage, and ultimately premature erythrocyte clearance and hemolysis [46,47]. Similar hematological toxicity has been observed in preclinical animal studies [48,49]. Metabolomic analyses of erythrocytes from ribavirin-treated patients further demonstrated cellular ATP depletion accompanied by decreased levels of glycolytic and PPP metabolites, supporting a link between intracellular ribavirin metabolism and impaired erythrocyte energy metabolism [50]. These findings illustrate that while intracellular phosphorylation enhances the pharmacological activity of ribavirin, prolonged accumulation of its phosphorylated metabolites can contribute to toxicity, including glycolysis inhibition.

### Limitations of the study

Schrödinger Glide is a molecular docking method widely used for hit screening from large compound libraries. It has been extensively validated across diverse receptor targets and generally provides good enrichment of active compounds over decoys, making it a valuable tool for hit identification [18,51]. Glide offers two scoring functions: GlideScore, which is commonly used for ranking chemically diverse compounds, and Emodel, which is force-field-weighted and optimized for pose selection. Although not designed for enrichment, Emodel outperformed GlideScore in distinguishing GPI binders from non-binders, ranking nearly all actives (24 of 26) within the top 2.9%, above the 800 inactives. But when applied to virtual screening, Emodel showed modest performance, with a notable number of false positives enriched among the top-ranked compounds. Moreover, incorporation of explicit water molecules and H-bond constraints, which were thought to favor the interaction with GPI ligands, did not improve the enrichment. A more comprehensive evaluation of alternative water sites and H-bond constraint combinations would be required to fully assess their impact on the enrichment performance.

Generally, docking studies are often followed by molecular dynamics (MD) simulations and free-energy perturbation (FEP) calculations, which can better account for protein flexibility and solvent effects but are computationally intensive. Here, Prime MM-GBSA was subsequently employed to re-rank the Glide SP-docked poses. Although this method incorporates solvent effects and receptor relaxation, it cannot fully capture the dynamic behavior of protein-ligand interactions. More importantly, the selection of hits for experimental validation was influenced by practical considerations, including compound synthetic cost and commercial availability at the time of the study, rather than solely relying the docking and MM-GBSA scores. Only 32 out of 193 compounds were available for testing. This curation process introduced bias into the final hit selection and may have affected the assessment of the overall predictive performance of the virtual screening protocol.

Both the Glide Emodel and Prime MM-GBSA scoring functions tended to assign more favorable scores to larger compounds. The potent endogenous GPI inhibitor, E4P, is a relatively small compound (198.1 Da) and had a Emodel score of –110 kJ/mol, whereas the larger nucleotide-based sugar phosphates (320–500 Da) received more favorable scores (–120 to –150 kJ/mol) despite their weak inhibitory activity. Structurally related intermediates from the same pathway, such as ribulose-5-phosphate (Ru5P; CID439184) and sedoheptulose 7-phosphate (S7P; CID165007), had docking scores similar to E4P and share the same relative configuration of the two hydroxyl groups nearest the phosphoryl group. Notably, both compounds have been reported to inhibit GPI from non-mammalian species such as *Dictyostelium discoideum* [52] and may possess inhibitory activity on the human GPI. However, they were not evaluated in this study due to the practical limitations described above.

The *in vitro* cellular assays also had limitations that should be considered. These assays did not establish whether the observed changes in glycolytic flux and cell viability were resulted from direct inhibition of GPI within the cells, as many cytosolic enzymes are known to bind phosphorylated nucleotides and sugar derivatives. Many endogenous metabolites serve as allosteric regulators or substrates for multiple enzymes within glycolysis and interconnected metabolic pathways. For example, GlcN6P has been reported to inhibit human hexokinase in addition to GPI. Moreover, as discussed earlier, RMP has been shown to inhibit IMPDH besides its proposed interaction with GPI, and further phosphorylation by kinases generate metabolites capable of interacting with other cellular targets. Therefore, the observed decrease in glycolytic flux and cell viability may reflect the combined effects of these metabolites on multiple metabolic targets rather than GPI inhibition alone.

A further limitation is that the combination of glycolytic and mitochondrial inhibitors was evaluated using only a limited number of glycolytic inhibitors and a single test concentration for each inhibitor. Therefore, the absence of synergy should not be generalized beyond the assay conditions used in this study. Future studies employing broader concentration ranges and alternative compound combinations would be essential to better evaluate potential synergistic interactions. In addition, the cellular exposure of compounds represents another limitation, as intracellular uptake and compound concentrations were not determined in this study. Consequently, the observed effects on glycolytic flux and cell viability may be influenced by factors such as membrane permeability and intracellular stability, particularly for highly polar phosphate-containing compounds. Lastly, the receptor(s) responsible for cellular uptake of most of these compounds were not identified, and it is unclear whether their uptake is mediated by common receptor(s).

## Conclusion

Human GPI is an attractive target in cancer studies because of its role in glycolysis and extracellular function as autocrine motility factor, but it remains underexplored, possibly due to challenges in identifying selective inhibitors for highly conserved, polar active sites. Here, we combined structure-based virtual screening with biochemical, biophysical, and structural validation to discover new GPI inhibitors. Screening of 1.1 million compounds identified eleven competitive inhibitors, representing previously unreported GPI ligands. While their potencies are unlikely to be sufficient for therapeutic applications, these inhibitors provide valuable insight into the ligand recognition by the GPI active site, demonstrating its ability to accommodate a broad range of phosphorylated compounds. Seven of the identified GPI inhibitors are endogenous metabolites from central metabolic pathways in the cytosol. Since these pathways share a metabolic environment with glycolysis, their intermediates can directly interact with GPI, enabling a form of pathway crosstalk that links energy production to biosynthetic demand. Beyond the validated hits, virtual screening revealed dozens of putative active-site binders that warrant further investigation, including Ru5P and compounds bearing phosphate-mimicking groups. Notably, two inhibitors featuring phosphate bioisosteres provided proof of concept that phosphate replacement is compatible with GPI binding and may be considered for ligand optimization. In addition, the identification of RMP as a competitive GPI inhibitor broadens the known target profile of ribavirin, while the inhibitory effects of ribavirin and RMP on glycolytic flux and cell viability revealed glycolysis inhibition as a potential contributor to their anticancer activity alongside other established mechanisms.

## Methods

### Materials

All chemicals for buffer preparation were bought from Sigma-Aldrich (St. Louis, MO, USA) and Merk (Darmstadt, Germany). Bacterial culture components include Yeast extract and Casein peptone were purchased from Neogen (Lincoln, NE, USA); Bacto agar from Becton, Dickinson (Holdrege, NE, USA); L-Rhamnose, isopropyl b-D-1-thiogalactopyranoside (IPTG), Ampicillin, and Chloramphenicol from AppliChem GmbH (Darmstadt, Germany). *Leuconostoc mesenteroides* glucose-6-phosphate dehydrogenase (*Lm*G6PDH) for enzyme-coupled assay was purchased from Sigma-Aldrich. Enzymes substrates, G6P and F6P, were derived from Sigma and NADPH from GoldBio (St. Lous, USA). The Protein Thermal Shift™ Dye Kit (No. 4461146) was purchased from Thermo Fisher Scientific (Waltham, MA, USA). FrameStar® 96-well semi-skirted PCR plates (ABI® FastPlate style) and qPCR adhesive seals were obtained from Azenta (Burlington, VT, USA). Crystallization precipitant (PEG 3350) was derived from Molecular Dimensions (Sheffield, UK). Crystallization plates (24-well), 22 mm cover slips, and silicone grease were obtained from Hampton Research (Aliso Viejo, CA, USA). Crystal harvesting tools and dewar (cryoshipper) were from Molecular Dimensions. Test compounds for GPI assays include: Phosphonoacetic acid, Methylphosphonic acid, Inosine 5’-monophosphate, Xanthosine 5’-monophosphate, and N-Acetyl-a-D-glucosamine 1-phosphate were purchased from Sigma-Aldrich; Adenosine 5’-monophosphate, 2-Phosphonomethyl-pentanedioic acid, S-(N-Methylcarbamoyl)-glutathione, 6-Phosphogluconic acid, and PMPA were supplied by Santa Cruz Biotechnology (Dallas, Texas, USA); Ribavirin, Ribavirin monophosphate, Cytidine 5’-monophosphate, Glucosamine-6-phosphate, and L-Ascorbic acid 2-phosphate were from MedChemExpress (New Jersey, USA); N-acetyl glucosamine-6-phosphate, 3,3-diphosphonopropanoic acid, and other bisphosphonate compounds were ordered from Mcule (Budapest, Hungary); Dimethylallyl diphosphate was obtained from Fluorochem Limited (Derbyshire, UK). Clear-bottom 384-well microplates for activity assay readouts were purchased from Greiner Bio-One (Frickenhausen, Germany). Human embryonic kidney 293 (HEK-293; CRL-1573.3™) and human bone osteosarcoma epithelial (U2OS; HTB-96™) cell lines were obtained from ATCC (Manassas, VA, USA). Agilent Seahorse XFe96/XF Pro FluxPak, DMEM assay media, cell culture supplements (glucose, glutamine, pyruvate), and Glycolytic Rate Assay Kit were purchased from Agilent Technologies (Santa Clara, CA USA).

### Human GPI expression & purification

Human GPI was expressed in *E. coli*, purified, and characterized as previously described [16]. Protein purity (>97%) was assessed by SDS-PAGE analysis. The protein samples were stored at −80 °C in buffer containing 100 mM Tris, pH 7.4, 150 mM NaCl. For crystallization experiments, the protein solution was buffer-exchanged into 25 mM Tris-HCl, pH 7.4, containing 50 mM NaCl, passed through a 0.22 µm Millipore filter, and stored at −80 °C.

### Virtual screening

Glide docking and MM-GBSA from Schrödinger Suite were used for library screening and post-docking analysis. Since rigid docking was used as the screening strategy, docking grids generated from multiple human and mammalian GPI structures, in the presence or absence of crystallographic waters, were compared to select for the most suitable grid for docking. For grid generation, the GPI structures were prepared using the Protein Preparation Wizard in Maestro, with all water molecules removed or retained during the process. Docking grids were generated by centering the grid box on residues within GPI active site, with grid box dimensions set to 12 × 12 × 12 Å. First, cross-docking was performed to evaluate the accuracy of the grids in pose prediction, and results are summarized in figures S14 and S15 in supplementary data. Overall, hydrated grids yielded slightly lower RMSD values compared to dry grids (Suppl. Fig. S16), indicating improved pose prediction accuracy. However, visual inspection revealed that the predicted ligand orientations and interactions with key residues were largely consistent between the two conditions. Moreover, in both cases, the docked poses closely resemble the cognate poses in crystal structures, suggesting that inclusion of water molecules had a little impact on the overall binding mode.

Enrichment analysis was performed for both dry and hydrated grids to evaluate their ability in distinguishing between GPI active-site binders and non-binders. The active set comprised 24 phosphorylated ligands and 3 non-phosphorylated inhibitors of GPI (Suppl. Fig. S17), whereas the inactive set consisted of 800 metabolites previously screened against GPI with no observed effect on its activity [16]. Notably, the inactive set also included multiple phosphorylated compounds, ensuring that the enrichment was not biased towards phosphate-containing compounds. The enrichment results are shown in Suppl. Table S6 and S7. No significant correlation was observed between the docking accuracy (RMSD) and early enrichment performance (indicated by BEDROC) of the docking grids (Suppl. Fig. S18). Overall, the dehydrated grids performed better at retrieving GPI actives than the hydrated ones, suggesting that inclusion of water molecules may introduce unfavorable steric constraints in rigid docking. Furthermore, the Emodel scoring function showed better performance than Glide gscore in ranking the actives and was therefore used as the primary metric for prioritizing potential binders. The dry grid derived from the human GPI structure (PDB ID: 6XUH) was used for subsequent library screening study, as it showed the best performance among the grids, with a receiver operational characteristic (ROC) value of 0.99, Boltzmann-Enhanced Discrimination of ROC (BEDROC) values of 0.996 (α = 160.9) and 0.950 (α = 20.0). All 24 phosphorylated inhibitors were ranked within the top 2.9% of compounds, above all inactives, while two fragment inhibitors were ranked within the top 6%, and one fragment within the top 28%, indicating strong discrimination between actives and inactives. Adding explicit water molecules and H-bond constraints to this grid did not improve enrichment (Suppl. Table S8 & S9), thus, no further modification was considered.

Compound libraries for virtual screening were sourced from following databases: Zinc15/20, Enamine, ChEMBL, PubChem, Mcule, and eMolecules. The compound libraries were curated to represent two distinct chemical spaces: drug-like and hydrophilic. The drug-like screening library was filtered according to Lipinski’s Rule of Five, selecting compounds with a molecular weight ≤ 500 Da, a partition coefficient (logP) ≤ 5, no more than five hydrogen bond donors (HBD), no more than ten hydrogen bond acceptors (HBA), ten or fewer rotatable bonds, and a polar surface area (PSA) ≤140 Å². Following filtering, 130,322 drug-like compounds were retained from an initial set of 237,572 compounds obtained from Zinc15/20 libraries (Biogenic/Metabolite, Natural Products, and FDA-approved) and Enamine collections (DDS-10, DDS-50, Bioactive Bioreference, Natural Product-Like, and Phenotypic Screen libraries). The hydrophilic screening library comprised compounds from ChEMBL, ZINC20, PubChem, MCule, eMolecules, and Enamine collections. 5,085,920 compounds were filtered based on an AlogP range of –6 to –1.5 and Mw of 100–550 Da, yielding a final screening library of 988,321 compounds. To reduce redundancy, duplicate compounds across the libraries were removed.

Prior to docking, the ligands were prepared using LigPrep, to apply the OPLS force-field for energy minimization, and to optimize the structures and add hydrogen atoms. Furthermore, Epik was used to assign the protonation states at pH 7.4 ± 2 and possible tautomers were generated for each molecule by determining their chirality from 3D structures. For initial screening, compound libraries were docked into the GPI active site in HTVS mode, with flexible ligand sampling. The top 20% of ligands from the HTVS stage (ranked based on Emodel scores) were re-docked into the same binding site using the standard precision (SP) docking mode. A hit selection threshold was then applied based on the Glide Emodel scores of known GPI binders (−161.5 to −104.0 kJ/mol). Compounds with Emodel scores ≤ −104.0 kJ/mol were retained as putative binders for further analysis by Prime MM-GBSA (Molecular Mechanics-Generalized Born Surface Area), allowing side-chain flexibility in the GPI active site. Following MM-GBSA analysis, the compounds were re-ranked by their binding free energy scores (MM-GBSA dG bind). The hit selection threshold was established using the MM-GBSA dG bind scores of known GPI binders, which ranged from −41.6 to −5.8 kJ/mol. Compounds with MM-GBSA binding scores ≤ −5.8 kJ/mol were selected for further curation. Because the hydrophilic screen generated a large number of putative hits with favorable docking scores, additional filtering was performed to reduce the number of candidates while preserving chemical diversity. To this end, selected compounds from SP docking stage were clustered by Fingerprint Similarity using dendritic fingerprints with Daylight invariant atom types and bonds distinguished by bond order. Compound similarity was evaluated using the Tanimoto coefficient. Top 20% of representatives from each cluster were subjected to binding free energy calculations using Prime MM-GBSA and curated based on their MM-GBSA dG bind scores.

### Enzyme activity assays

Human GPI activity was measured in gluconeogenesis direction (F6P to G6P) using a coupled assay with *Lm*G6PDH as the coupling enzyme, which reduces NADP⁺ to NADPH. Absorbance measurements were carried in 384-well plates and NADPH formation was monitored at 340 nm using an iD5 multimodal microplate reader (Molecular Devices, UK). All enzymes and reagents were solubilized in assay buffer containing 100 mM Tris pH 7.4, 150 mM NaCl. For GPI activity measurement, the final assay concentrations of GPI and *Lm*G6PDH were set to 5 ng/mL of dimeric GPI and 50 µg/mL *Lm*G6PDH, respectively. In dose-response assays, 1 mM of F6P, 5 mM of NADP+ were applied, and ligands were tested at different concentrations. To determine Michaelis-Menten kinetics of GPI, the concentration of F6P was varied from 0.05 to 4 mM, with all other assay components unchanged and ligand concentrations kept fixed. In counter-screen assay, *Lm*G6PDH activity was measured with an assay mixture containing 5 µg/mL *Lm*G6PDH, 1 mM G6P, and 5 mM NADP+. For negative control, where GPI is 100% active (no inhibition), the assay buffer was utilized instead of test compounds, and for positive control, 6-phophogluconate, a competitive GPI inhibitor was employed. In all measurements, the enzyme was pre-incubated with test compounds at 25 °C for 30 min, and substrates were subsequently added, and activity measured at 25 °C, over 3 min at 12 sec intervals.

### MST binding assay

Binding interactions between GPI and the identified hits were studied using microscale thermophoresis (MST) on a Monolith NT.LabelFree instrument with NT.LabelFree capillaries from NanoTemper Technologies GmbH (Munich, Germany). All test compounds were prepared in MST buffer (100 mM Tris, pH 7.4, 150 mM NaCl). Prior to measurement, the target enzyme was centrifuged at 5,000 ×g for 3 min and added to the compound solutions prepared in the same buffer. The mixtures were then centrifuged at 5,000 ×g for 3 min and equilibrated at room temperature for 30 min. MST measurements were performed at 25 °C using 20% excitation-and 40% MST power, with on-time response set to at least 10 s. The enzyme concentration was kept constant at 0.2 µM (dimeric form) of GPI. For K_d_ determination, two-or three-fold serial dilutions of test compounds were titrated against a constant enzyme concentration (0.2 µM), with ligand concentrations ranging from 0.01 to 50 mM. All compounds were tested in at least duplicate in full 12-point titration experiments. In the event of variation in initial fluorescence and photobleaching rate, SDS denaturation tests were performed to distinguish between binding-specific and ligand-induced changes in target fluorescence, following a previously described protocol [16].

### DSF thermal shift assay

Differential scanning fluorometry (DSF) thermal shift assay was employed as validation step to confirm the ligand-enzyme interaction. The DSF assay was carried out in 96-well qPCR plates with a final assay volume of 20 µL. Fluorescence measurements were performed using a QuantStudio™ 3 Real-Time PCR System (Thermo Fisher Scientific) and ROX dye (Thermo Fisher Scientific) as the fluorescent reporter, with excitation and emission wavelengths of 580 ± 10 nm and 623 ± 14 nm, respectively. Assay mixtures consisted of ∼0.5 µM of dimeric GPI, 0-40 mM test compound, MST buffer supplemented with 0.001% (w/v) Triton X-100, and 1× ROX dye. Each ligand was tested at 2–3 concentrations, with each condition measured in triplicate. Additionally, 8-point titration was performed for selected ligands in duplicate. Enzyme-ligand mixtures containing varying ligand concentrations (0–45 mM) were incubated at 4 °C for 10 min prior to dye addition, followed by centrifugation at 3000 ×g and 4 °C for 3 min. Before measurements, the assay mixtures were equilibrated at 25 °C for 2 min in the PCR instrument. Fluorescence changes were then monitored over a temperature range of 25–90°C with an increment rate of 0.05 °C/s. No background fluorescence signal was detected when the test compounds were assayed alone. Melt curve data were analyzed in GraphPad Prism by calculating the first derivative of the relative fluorescence units (RFU) as a function of temperature. To allow clearer comparison between control and ligand-treated samples, the first-derivative curves were normalized and smoothed in GraphPad Prism.

### Protein X-ray crystallography

Co-crystallization of human GPI with ligands was performed using the hanging-drop vapor diffusion method. All protein-, ligand-, and buffer solutions were filtered through 0.22 µm membrane filters prior to use. The GPI stock solution was prepared in buffer containing 25 mM Tris, pH 7.4 and 50 mM NaCl. Before co-crystallization, 8 mg/mL of GPI protein was incubated with ligand (15-25 mM) at 4°C for 1 hour. Protein crystals were obtained by mixing 1.5 µL of the GPI-ligand mixture with 0.5 µL of crushed, diluted GPI crystal seeds and 1.5 µL of precipitation solution comprised of reservoir solution (23.5% PEG3350, 58 mM HEPES pH 7.4) and corresponding ligand at a defined concentration. Crystals were grown at ambient temperature for 4-5 days prior to harvesting. For cryoprotection, crystals were briefly transferred into a cryoprotectant solution consisting of mother liquor supplemented with 18-20% w/v glycerol. Crystals were snap-frozen in liquid nitrogen and stored in a dewar for transport to synchrotron facilities. X-ray diffraction data were collected at beamline P11 at PETRA III (DESY, Hamburg, Germany) and beamline BioMAX at MAX IV (Lund, Sweden). Diffraction data were manually processed using XDS. Molecular replacement was performed with Phaser (CCP4 suite) using a previously reported human GPI structure (PDB ID: 9FHF) as the reference model. Structure refinement was carried out using phenix.refine (Phenix suite), and manual model building was performed in Coot.

### Metabolic flux measurement

The human embryonic kidney 293 (HEK-293) and human bone osteosarcoma epithelial (U2OS) cell lines were obtained from ATCC. All culture and assay media, and buffer reagents were pre-warmed to 37°C before use. Phosphate Buffered Saline (PBS) buffer was filtered, autoclaved, and then sterile-filtered inside a laminar flow cell-culture hood. All cells were grown in Gibco™ DMEM medium containing GlutaMAX™, glucose, and pyruvate, supplemented with 1% penicillin/streptomycin and 10% v/v Gibco™ fetal bovine serum (FBS); all from Thermo Fisher Scientific. For recovery, frozen cells were thawed in 37°C water bath for 2 min, and the cell suspension was transferred into the complete medium and centrifuged at 250 × g for 5 min. After removal of the supernatant, the cell pellet was resuspended in the complete growth medium and seeded into T25 flasks. The initial seeding densities for HEK-293 and U2OS cells were 2.0×10^5^ cells/mL and 1.5×10^5^ cells/mL, respectively. Cell cultures were maintained at 37°C in a 5% v/v CO_2_ incubator with constant humidity. Prior to flux measurement, the cells were passaged for 4 or 5 times upon reaching between 80-85% confluency and with 95-98% viability. For detachment, the cells were rinsed with 1× PBS buffer and treated with trypsinization solution containing 0.25% (w/v) trypsin and 0.025% (w/v) EDTA. Viability of the cells was monitored by trypan blue exclusion using automated cell counter – Countess II (Thermo Scientific, US). Two days before flux assay, the HEK-293 cells were seeded at ∼1.0×10^4^ cells/100 µL and U2OS cells ∼5.000 cells/100 µL to ensure 80-90% confluency on the day of measurement.

After seeding, the cells were allowed to rest inside the laminar flow cell-culture hood at ambient temperature for an hour before placing into the incubator. 24h before the assay, sensor plates were hydrated with the Agilent Seahorse Calibrant buffer and incubated at 37 °C, without CO_2_. On the day of measurement, 7 μL of PrestoBlue reagent (Thermo Fisher) was added to each well of the culture plate, followed by incubation for 1 h at 37 °C, 5% CO₂. The medium was subsequently removed, and absorbance was measured for cell viability assessment and normalization. Cells were gently washed twice with 100 μL of XF assay medium (Agilent DMEM Running Medium supplemented with 5 mM HEPES, pH 7.4, 10 mM glucose, 1 mM pyruvate, and 2 mM glutamine). Thereafter, 180 μL of XF assay medium was added to each well, and the plate was incubated at 37 °C under CO₂-free condition for 1 h to allow equilibration. The medium was then replaced with fresh XF assay medium before the flux measurements.

All test compounds and assay inhibitors were reconstituted in XF assay medium, sterile-filtered, and loaded into the sensor cartridge ports as follows: port A contained 2–8 mM ribavirin monophosphate (RVP), 5–20 mM xanthosine monophosphate (XMP), or 5–20 mM glucosamine-6-phosphate (GlcN6P); port B loaded with 5 μM rotenone/antimycin A (Rot/AA); and port C contained 50 mM 2-deoxy-D-glucose (2-DG). This configuration enabled sequential injection of the test compounds and assay inhibitors during the measurement. For control, the XF assay medium was added into port A. The flux measurement was performed using Agilent Seahorse XFe96 Analyzer, which had been pre-equilibrated at 37 °C overnight. Following sensor cartridge calibration, the assays were run at 37 °C with 5-7 measurements collected after each injection; five or six replicates were used per condition. Cell viability was assessed using PrestoBlue reagent (Invitrogen™, Carlsbad, CA, USA) by measuring absorbance at 570 and 600 nm as described in the manufacturer’s protocol. Proton efflux rate (PER), calculated from the measured extracellular acidification rate (ECAR), was used to approximate the rate of glycolytic flux.

### Cell viability assay

The effect of test compounds on cell viability was assessed using the PrestoBlue assay. Cell viability assessment was undertaken using the same cell lines as those employed in glycolytic flux measurements. All cells were cultured in complete DMEM media (described above) in Nunclon Delta-Treated 96-well microplates (ThermoFisher, USA) and incubated at 37°C in a 5% v/v CO_2_ incubator. To avoid excessive cell confluency during the measurement period, HEK-293 cells were seeded at ∼1.0 ×10^5^ cells/mL, whereas U2OS cells were seeded at ∼4.0×10^4^ cells/mL, in a final volume of 100 µL per well. The cell viability was monitored for every 24 hours over a course of four days, with measurements performed 24 h after each daily medium replacement (100 µL per well). On day 1 (24 h after seeding), all cells received fresh complete medium without inhibitors. On days 2-4, cells in the treatment groups were supplied with fresh inhibitor-containing medium, whereas control cells received fresh complete medium.

The inhibitors were tested at the following concentrations: RMP (2–8 mM), XMP (5–20 mM), GlcN6P (5–20 mM), CMP (5–20 mM), 2-deoxyglucose (2-DG; 10–20 mM), 6-phosphogluconate (6PG; 10–20 mM), ribavirin (5–20 mM), and rotenone/antimycin A (Rot/AA; 1.0 µM). Before the viability measurement, the cells were treated with PrestoBlue (7 µL per well) and incubated for 1 hour at 37°C with 5% v/v CO_2_. The PrestoBlue-treated media were then transferred to 96-well microplates for absorbance measurement at 570 and 600 nm. After removal of the medium, U2OS cells were gently washed three times with 180 µL of 1× PBS and replenished with 100 µL of fresh medium. For HEK-293 cells, the wells were washed thrice with 180 µL of fresh complete medium (instead of PBS) before replenishment with 100 µL of the complete medium, using slow pipetting to minimize cell detachment. During all washing steps, approximately 20 µL of residual medium or PBS buffer was left in each well to prevent drying of the cells.

## Acknowledgements

We acknowledge MAX IV Laboratory for beamtime on Beamline BioMAX under Proposal 20250347, and Research conducted at MAX IV, a Swedish national user facility, is supported by the Swedish Research council under contract 2018-07152, the Swedish Governmental Agency for Innovation Systems under contract 2018-04969, and Formas under contract 2019-02496. We acknowledge DESY (Hamburg, Germany), a member of the Helmholtz Association HGF, for the provision of beamtime and research carried out at PETRA III P11 under the proposal BAG-20230676. Snædís Ragnarsdóttir at the Department of medicine at the University of Iceland is gratefully thanked for providing the human cell lines. Óttar Rolfsson at the Department of medicine at the University of Iceland is specially thanked for the access to Seahorse XFe96 instrument. Funding Sources. Financial support from the Icelandic Research Fund (project 239695-051) and the Icelandic Cancer Society is gratefully acknowledged.

## Author contributions

**YYJ** planned and conducted the virtual screening and experimental work, performed the data analysis, and wrote the manuscript. **JGH** contributed to theoretical and experiment planning; provided reagents and access to facilities, and critically reviewed the manuscript. **KT** provided access to Schrödinger software license and technical advice on virtual screening protocols, and critically review the manuscript.

Competing interest: The authors state no competing interest.

## Abbreviations

2-DG: 2-deoxy-glucose;
2PMPA: 2-phosphonomethyl pentanedioic acid;
33DP: 3,3-diphosphono-propanoic acid;
ALDO: aldolase;
AMP: adenosine 5′-monophosphate;
ASU: asymmetric unit;
BEDROC: Boltzmann-enhanced discrimination of ROC;
CMP: cytidine 5′-monophosphate;
DMAPP: dimethylallyl pyrophosphate;
DSF: differential scanning fluorometry;
EC_50_: half-maximal effective concentration;
ECAR: extracellular acidification rate;
ENO: enolase;
F_norm_: normalized fluorescence change (F_hot_/F_cold_);
G6PDH: glucose-6-phosphate dehydrogenase;
GAPDH: glyceraldehyde-3-phosphate dehydrogenase;
GlcN6P: D-glucosamine 6-phosphate;
Glide: Grid-based Ligand Docking with Energetics;
GPI: glucose-6-phosphate isomerase;
GlycoPER: glycolytic proton efflux rate;
HK: hexokinase;
IC_50_: half-maximal inhibitory concentration;
IMP: inosine 5′-monophosphate;
K_i_: inhibition constant;
MDP: methylene diphosphonic acid;
MM-GBSA: molecular mechanics/generalized Born surface area;
MOA: mode/mechanism of action;
MST: microscale thermophoresis;
NAG6P: N-acetyl-D-glucosamine 6-phosphate;
PAE: phosphonoacetic acid;
PER: proton efflux rate;
PFK-1: phosphofructokinase-1;
PGK: phosphoglycerate kinase;
PGM: phosphoglycerate mutase;
PK: pyruvate kinase;
R5P: D-ribose 5-phosphate;
RMP: ribavirin 5′-monophosphate;
ROC: receiver operating characteristic;
AUC: area under the curve;
RMSD: root-mean-square deviation;
Rot/AA: rotenone/antimycin;
TPI: triosephosphate isomerase;
XMP: xanthosine 5′-monophosphate.

