## Supplementary material for "Structure-based virtual screening reveals new metabolite interactions and modulators of human glucose-6-phosphate isomerase"

**The file contains the following contents:**

1. Supplementary tables for Tab. S1-S9
2. Supplementary figures Fig. S1-S18

### Supplementary tables

#### Screening of drug-like libraries against GPI active site

Table S 1: Confirmed GPI hit - dimethylallyl pyrophosphate (DMAPP) and false positives from the drug-like screen

| | | | |
|---|---|---|---|
| 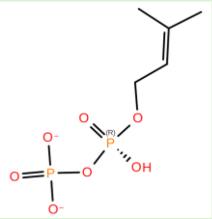<br>(DMAPP) | 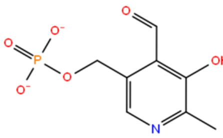 | 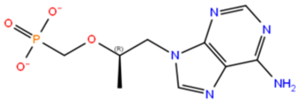 | 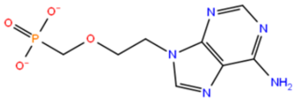 |

| | | | |
|---|---|---|---|
| 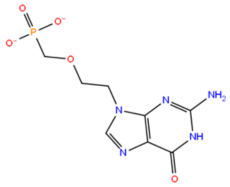 | 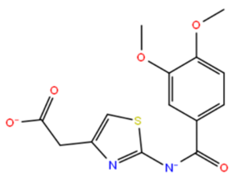 | 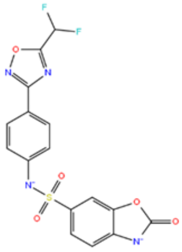 | 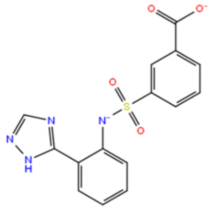 |
| 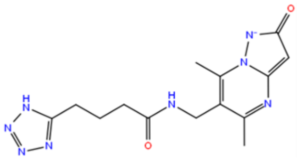 | 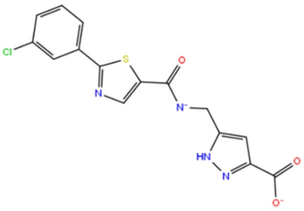 | 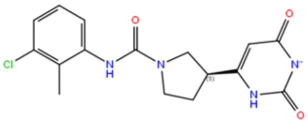 | 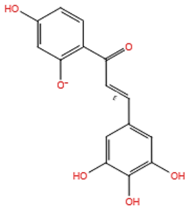 |

#### Screening of hydrophilic-compound libraries against GPI active site

Table S 2: Confirmed GPI inhibitors (highlighted in green) and false positives from the screening of hydrophilic compound libraries

| | | | | |
|---|---|---|---|---|
| 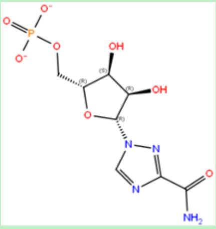 | 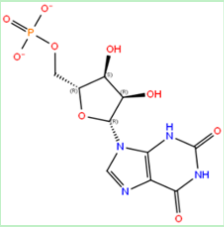 | 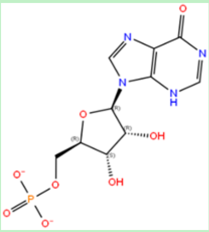 | 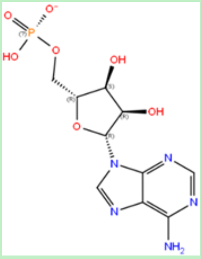 | 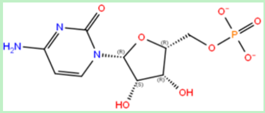 |
| 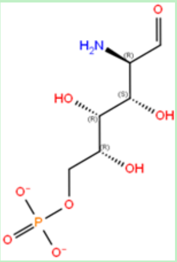 | 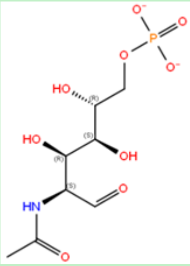 | 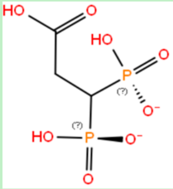 | 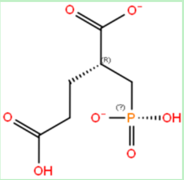 | 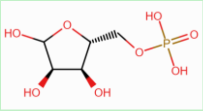 |
| 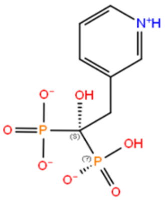 | 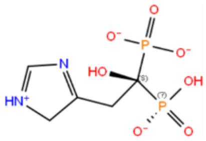 | 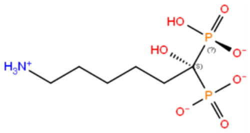 | 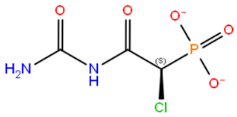 | 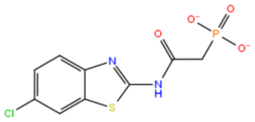 |

| | | | | |
|---|---|---|---|---|
| 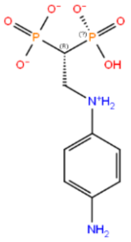 | 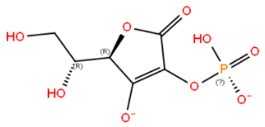 | 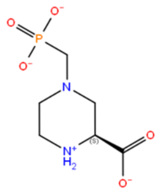 | 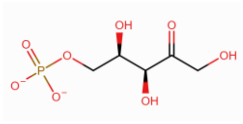 | 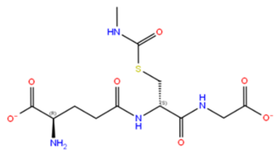 |

Table S 3: RMSD between docked poses (grey) and crystallographic poses of human GPI ligands: RMP (pink), XMP (purple), CMP (green), GlcN6P (orange), NAG6P (yellow), and DMAPP (turquoise).

| | | |
|---|---|---|
| 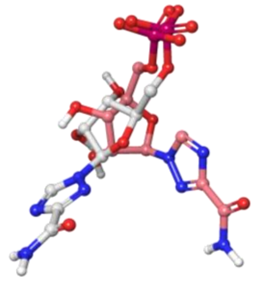 | 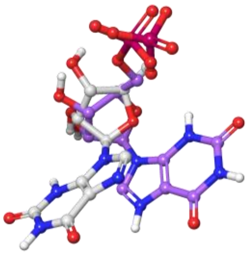 | 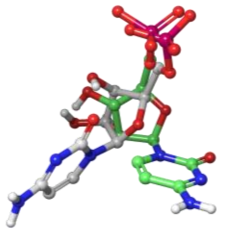 |
| **RMP**<br>**RMSD = 5.767 Å** | **XMP**<br>**RMSD = 4.732 Å** | **CMP**<br>**RMSD = 4.962 Å** |
| 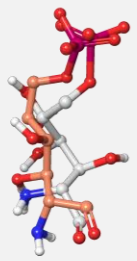 | 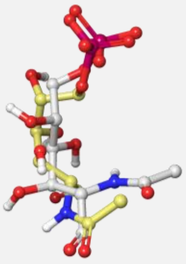 | 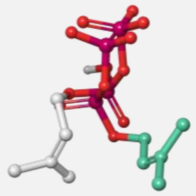 |
| **GlcN6P**<br>**RMSD = 2.413 Å** | **NAG6P**<br>**RMSD = 3.501 Å** | **DMAPP**<br>**RMSD = 4.865 Å** |

Table S 4: Zig-zag and Fischer projections of GPI ligands and false-positive hit

| | Zig-zag projections | Fischer projections |
|---|---|---|
| **D-Xylulose 5-phosphate (False-positive)** | 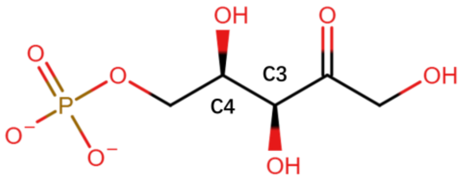 | 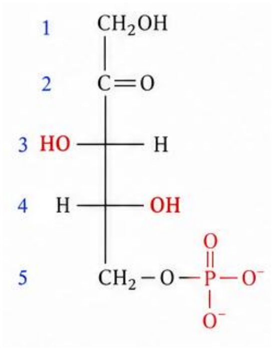 |
| **D-Erythrose 4-phosphate** | 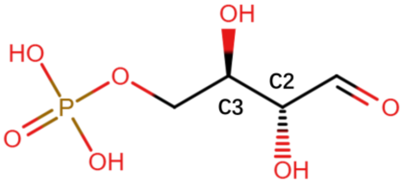 | 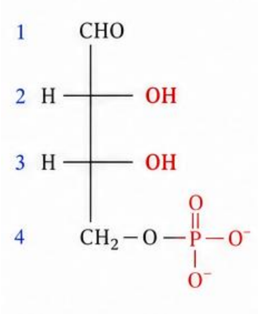 |
| **6-Phosphogluconate** | 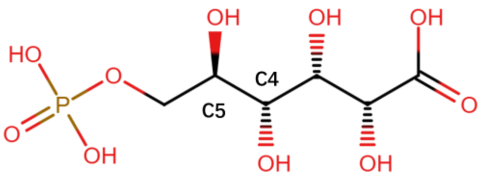 | 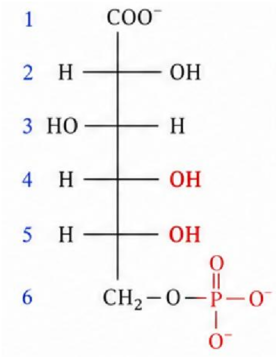 |
| **D-Arabinose 5-phosphate** | 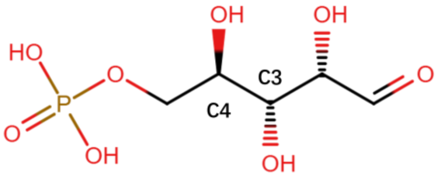 | 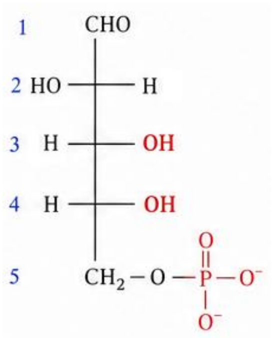 |
| **D-Ribose 5-phosphate** | 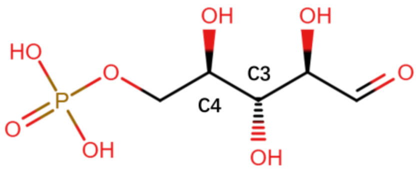 | 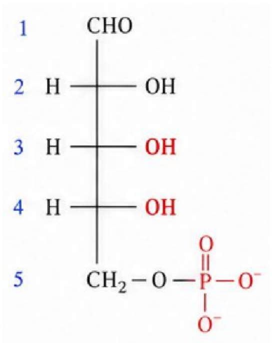 |

D-Glucosamine 6-phosphate

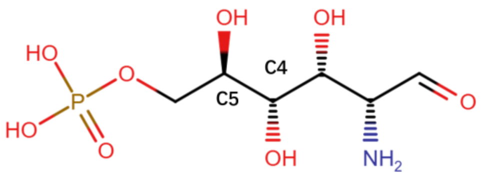


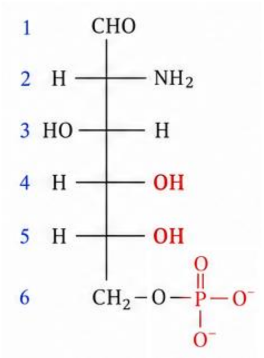

D-Sorbitol 6-phosphate

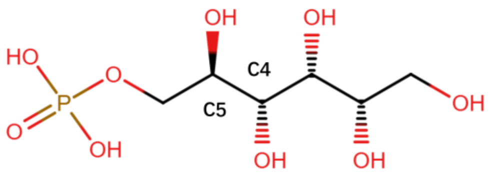


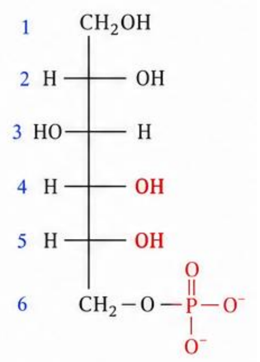

D-Fructose 6-phosphate

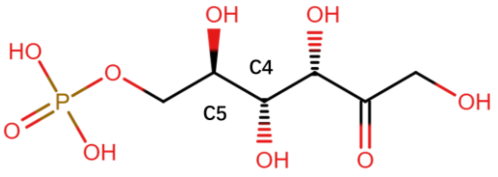


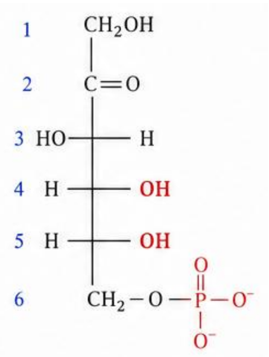

D-Glucose 6-phosphate

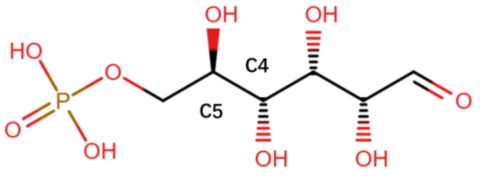


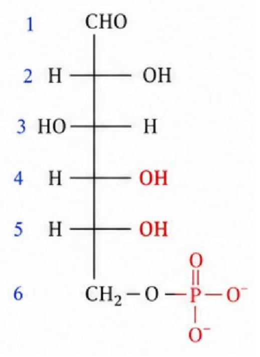

Table S 5: Ligand interaction diagrams showing GPI ligands before and after replacement of phosphoryl group with methyl diphosphonate (MDP) or phosphonoacetate (PAE). Glide (Emodel) docking and MM-GBSA scores are shown in kJ/mol.

| **Original ligand** | **Replacement with MDP** | **Replacement with PAE** |
|---|---|---|
| 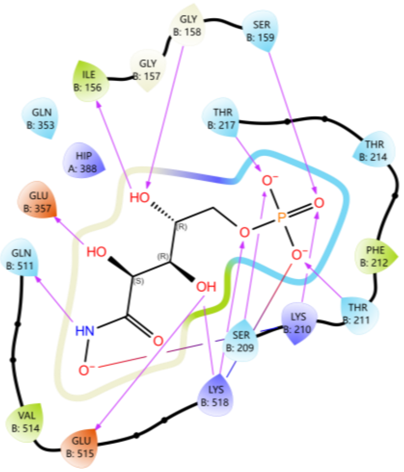<br>Glide Emodel = -141.79<br>MM-GBSA = -22.38 | 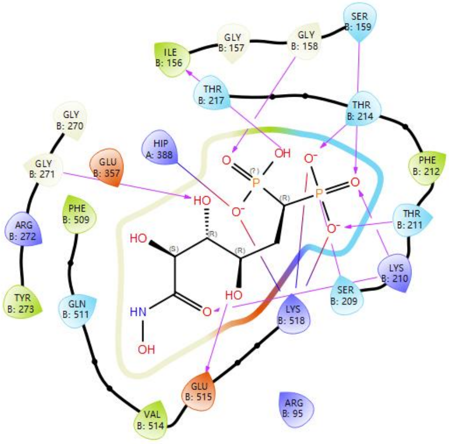<br>Glide Emodel = -145.28<br>MM-GBSA = -20.11 | 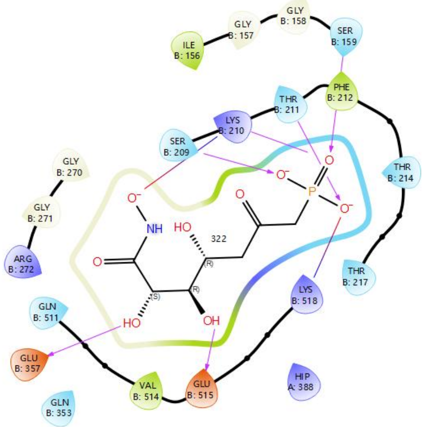<br>Glide Emodel = -134.82<br>MM-GBSA = -18.28 |
| 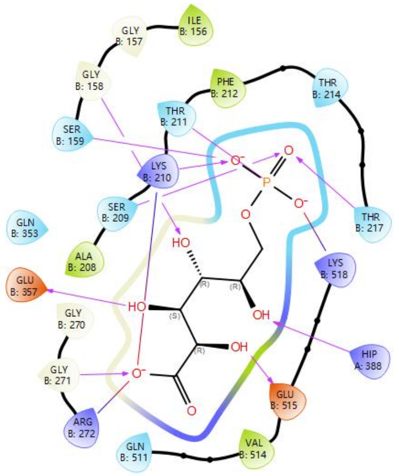<br>Glide Emodel = -139.60<br>MM-GBSA = -21.70 | 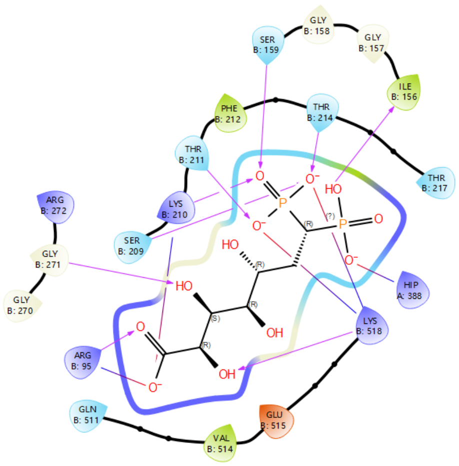<br>Glide Emodel = -155.45<br>MM-GBSA = -14.63 | 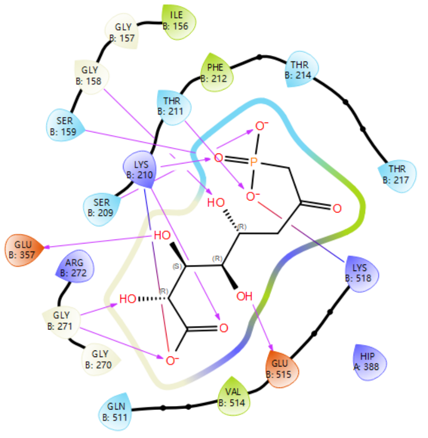<br>Glide Emodel = -127.98<br>MM-GBSA = -15.61 |

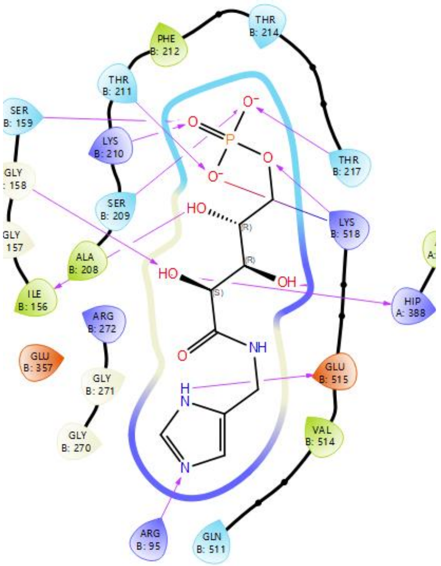

Glide Emodel = -153.22

MM-GBSA = -22.16

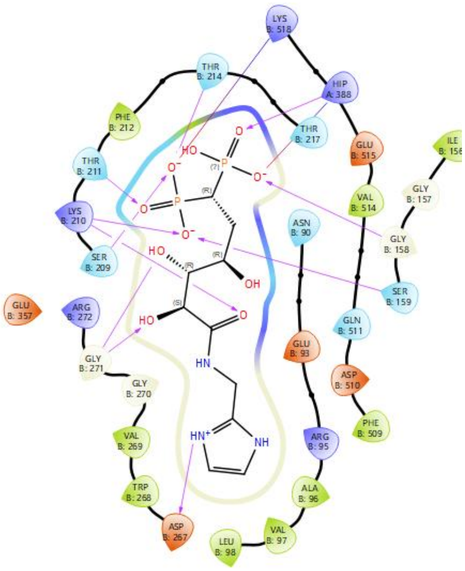

Glide Emodel = -155.45

MM-GBSA = -14.63

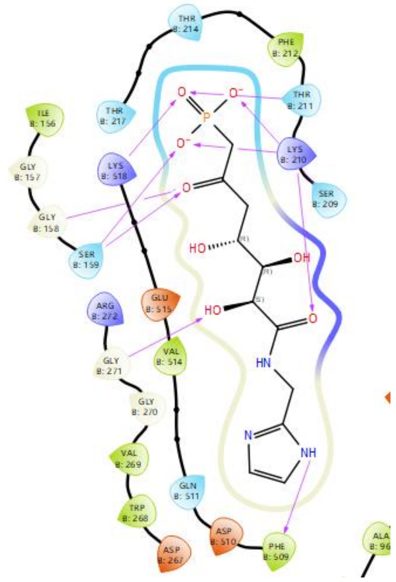

Glide Emodel = -143.43

MM-GBSA = -21.38

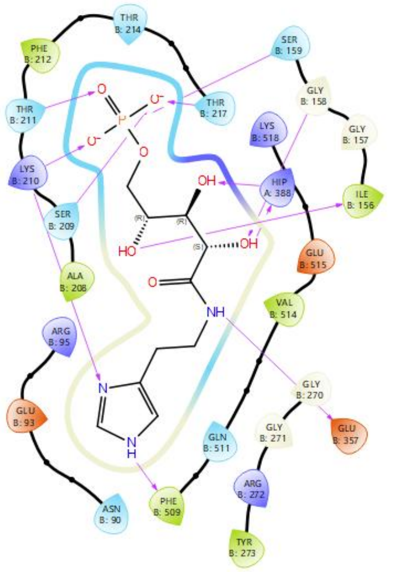

Glide Emodel = -141.59

MM-GBSA = -24.53

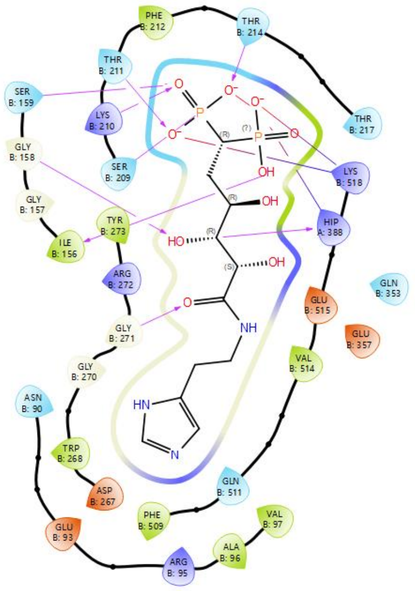

Glide Emodel = -165.09

MM-GBSA = -18.85

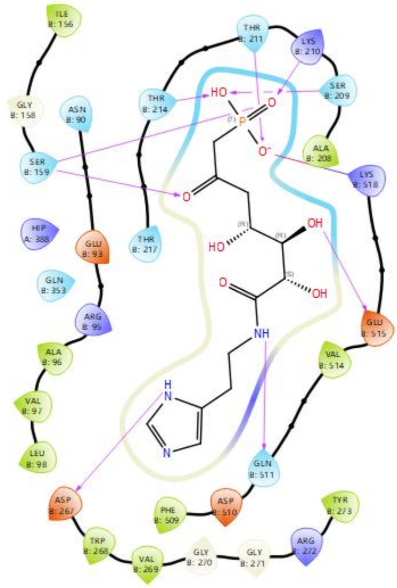

Glide Emodel = -145.63

MM-GBSA = -21.90

| | | |
|---|---|---|
| 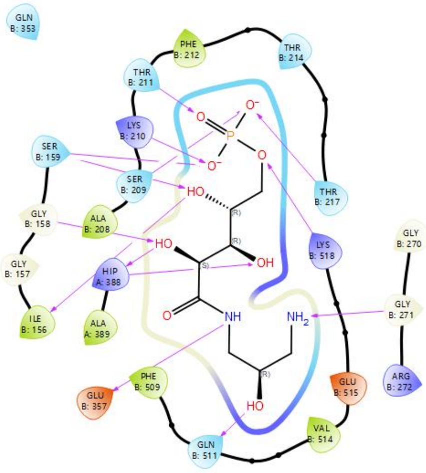 | 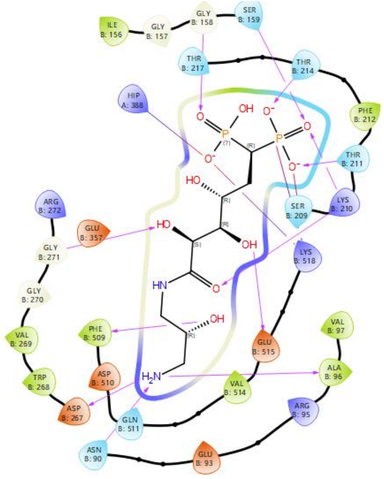 | 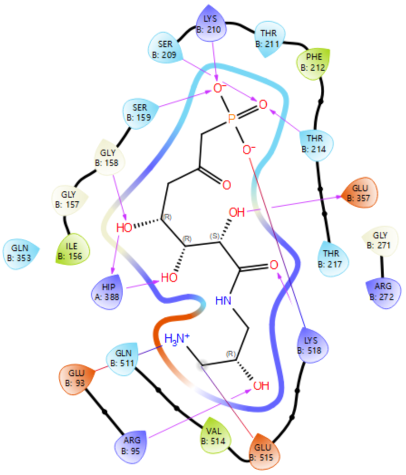 |
| Glide Emodel = -148.59 | Glide Emodel = -167.92 | Glide Emodel = -126.53 |
| MM-GBSA = -27.39 | MM-GBSA = -16.82 | MM-GBSA = -17.46 |

**Table S 6: Enrichment analysis of hydrated GPI grids using GlideScore and Emodel as scoring functions. Enrichment was evaluated using a set comprising known GPI binders as active compounds and 800 non-binders as decoys. The ability of GlideScore and Emodel to prioritize GPI actives was assessed by the number of actives recovered within the top 3% of ranked compounds and the enrichment metrics BEDROC, ROC-AUC, and RIE.**

| | | GlideScore | | | | Emodel | | | |
|---|---|---|---|---|---|---|---|---|---|
| **PDB CODE** | Average RMSD | BEDROC (α = 160.9) | # actives in top 3% | ROC | RIE | BEDROC (α = 160.9) | # actives in top 3% | ROC | RIE |
| **1XTB** | 1.806 | 0.003 | 0 | 0.81 | 1.92 | 0.909 | 12 | 0.91 | 8.67 |
| **2CXP** | 1.790 | 0.000 | 0 | 0.76 | 1.76 | 0.851 | 10 | 0.89 | 8.87 |
| **2CXR** | 1.921 | 0.035 | 4 | 0.83 | 2.76 | 0.519 | 16 | 0.94 | 8.88 |
| **6XUH** | 2.214 | 0.054 | 3 | 0.92 | 3.53 | 0.817 | 20 | 0.95 | 8.66 |
| **6XUI** | 2.327 | 0.021 | 2 | 0.90 | 5.85 | 0.894 | 18 | 0.96 | 8.89 |
| **1HOX** | 2.650 | 0.374 | 4 | 0.75 | 2.76 | 0.476 | 12 | 0.88 | 6.94 |
| **1IRI** | 2.634 | 0.000 | 0 | 0.70 | 0.45 | 0.359 | 9 | 0.87 | 6.26 |
| **1KOJ** | 1.976 | 0.000 | 0 | 0.85 | 1.38 | 0.784 | 20 | 0.96 | 10.22 |
| **1U0F** | 1.974 | 0.049 | 2 | 0.88 | 2.88 | 0.084 | 4 | 0.84 | 3.29 |

BEDROC (Boltzmann-Enhanced Discrimination of ROC); ROC (Receiver Operating Characteristic); RIE (Robust Initial Enhancement).

**Table S 7: Enrichment analysis of dry GPI grids using GlideScore and Emodel as scoring functions. Enrichment was evaluated using a set comprising known GPI binders as active compounds and 800 non-binders as decoys. The ability of GlideScore and Emodel to prioritize GPI actives was assessed by the number of actives recovered within the top 3% of ranked compounds and the enrichment metrics BEDROC, ROC-AUC, and RIE.**

| | | GlideScore | | | | Emodel | | | |
|---|---|---|---|---|---|---|---|---|---|
| PDB CODE | Average RMSD | BEDROC (α = 160.9) | # actives in top 3% | ROC | RIE | BEDROC (α = 160.9) | # actives in top 3% | ROC | RIE |
| **1XTB** | 2.398 | 0.001 | 1 | 0.84 | 1.45 | 0.440 | 13 | 0.91 | 11.29 |
| **2CXP** | 2.535 | 0.087 | 3 | 0.94 | 7.89 | 0.993 | 23 | 0.97 | 13.69 |
| **2CXR** | 2.152 | 0.014 | 1 | 0.89 | 3.92 | 0.628 | 15 | 0.97 | 11.87 |
| **6XUH** | **2.287** | **0.027** | **3** | **0.95** | **6.72** | **0.996** | **24** | **0.99** | **13.95** |
| **6XUI** | 2.430 | 0.146 | 1 | 0.90 | 3.21 | 0.930 | 19 | 0.98 | 13.00 |
| **1HOX** | 2.892 | 0.000 | 0 | 0.62 | 0.40 | 0.028 | 3 | 0.85 | 3.69 |
| **1IRI** | 2.291 | 0.002 | 1 | 0.69 | 1.14 | 0.036 | 3 | 0.89 | 4.10 |
| **1KOJ** | 2.397 | 0.000 | 0 | 0.86 | 1.91 | 0.919 | 20 | 0.98 | 13.12 |
| **1U0F** | 2.474 | 0.000 | 0 | 0.57 | 0.46 | 0.164 | 5 | 0.69 | 5.83 |
| **9FHF** | 2.605 | 0.174 | 8 | 0.94 | 7.09 | 0.758 | 20 | 0.98 | 12.66 |
| **9FCW** | 2.688 | 0.000 | 0 | 0.84 | 1.52 | 0.443 | 15 | 0.96 | 11.11 |

BEDROC (Boltzmann-Enhanced Discrimination of ROC); ROC (Receiver Operating Characteristic); RIE (Robust Initial Enhancement). Dry grid created from the active-site of 6XUH showed the best enrichment performance among all the evaluated docking grids.

**Table S 8: Conserved active-site residues involved in ligand binding in GPI crystal structures – highlighted in blue**

| | DHAP | PEP | MAE | CTA | 6PG | F6P | G6P | S6P | PA5 | E4P | A5P | O1B |
|---|---|---|---|---|---|---|---|---|---|---|---|---|
| **Arg95** | | | | | | | | | ■ | | | |
| **Ile156** | | | | | | | | | | ■ | | |
| **Gly157** | ■ | | | | | | | | | | | |
| **Gly158** | | | | | ■ | ■ | ■ | ■ | ■ | ■ | | ■ |
| **Ser159** | | | ■ | ■ | | ■ | ■ | | | | ■ | ■ |
| **Ser209** | ■ | ■ | ■ | ■ | ■ | ■ | | ■ | ■ | ■ | | ■ |
| **Lys210** | ■ | | | ■ | ■ | | ■ | ■ | ■ | ■ | ■ | ■ |
| **Thr211** | ■ | ■ | ■ | ■ | ■ | ■ | ■ | ■ | ■ | ■ | ■ | ■ |
| **Thr214** | ■ | | | ■ | ■ | ■ | | ■ | | ■ | | |
| **Arg272** | | | | | | | | ■ | | | | |
| **Gln353** | | | | | | | ■ | | ■ | | ■ | |
| **Glu357** | | | | | | | | | ■ | | ■ | ■ |
| **His388** | ■ | | | | ■ | ■ | ■ | ■ | ■ | ■ | ■ | ■ |
| **Lys518** | | | | | | | | | ■ | | | ■ |

The impact of water molecules and H-bond constraints on docking performance was examined using selected crystallographic waters and H-bond interactions that are consistently observed across multiple GPI crystal structures. Since rigid docking was employed, the inclusion of water molecules may have introduced steric clashes, there resulting in unfavorable poses and poorer enrichment. While docking with at least one H-bond constraint maintained comparable enrichment, applying two or more H-bond constraints reduced performance.

**Table S 9: Docking test on 6XUH grid with hydrogen-bond constraint or conserved water molecules**

| 6XUH grid | BEDROC (α = 160.9) | # actives in top 3% | ROC | RIE | Average RMSD |
|---|---|---|---|---|---|
| **At least 1 out of 4 H-bond constraints *** | 0.994 | 23 | 0.97 | 13.74 | 2.509 |
| **At least 2 out of 4 H-bond constraints *** | 0.614 | 19 | 0.93 | 10.88 | 2.835 |
| **At least 4 out of 10 H-bond constraints **** | 0.210 | 4 | 0.9 | 5.82 | 2.669 |
| **5 conserved H2O in active site***** | 0.436 | 17 | 0.92 | 8.52 | 2.335 |

* Most frequent H-bonding residues: Ser209, Lys210, Thr211, and His388.

** Residues forming H-bonds with ⩾2 ligands: Gly158, Ser159, Ser209, Lys210, Thr211, Thr214, Gln353, Glu357, His388, and Lys518.

*** Active-site water molecules that form H-bonds with ligands in at least four GPI crystal structures.

#### Supplementary figures

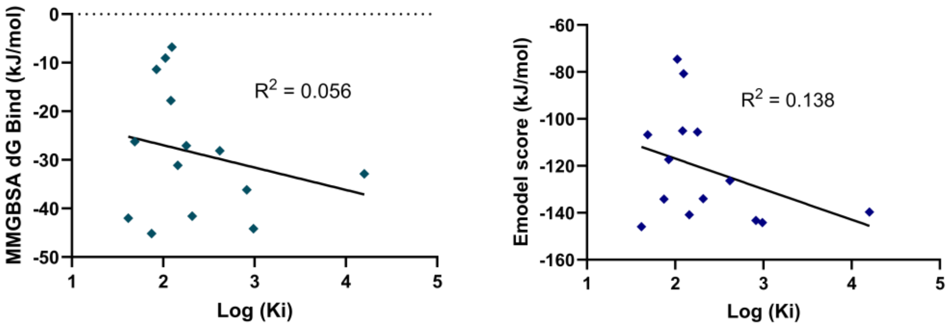


**Figure S 1: Relationship between docking/MM-GBSA scores and experimental activity (Log Ki).** Scatter plots showing the correlation between (left) GlideScore and logarithm of Ki and (right) Emodel scores and logarithm of Ki for nine identified GPI inhibitors and five previously reported inhibitors: 6-phosphogluconate, dihydroxyacetone phosphate, phosphoenolpyruvate, citraconate, and maleate. $R^2$ (Gscore) = 0.138; $R^2$ (Emodel) = 0.056.

#### Differential Scanning Fluorometry (DSF) – Thermal Shift Assay Results

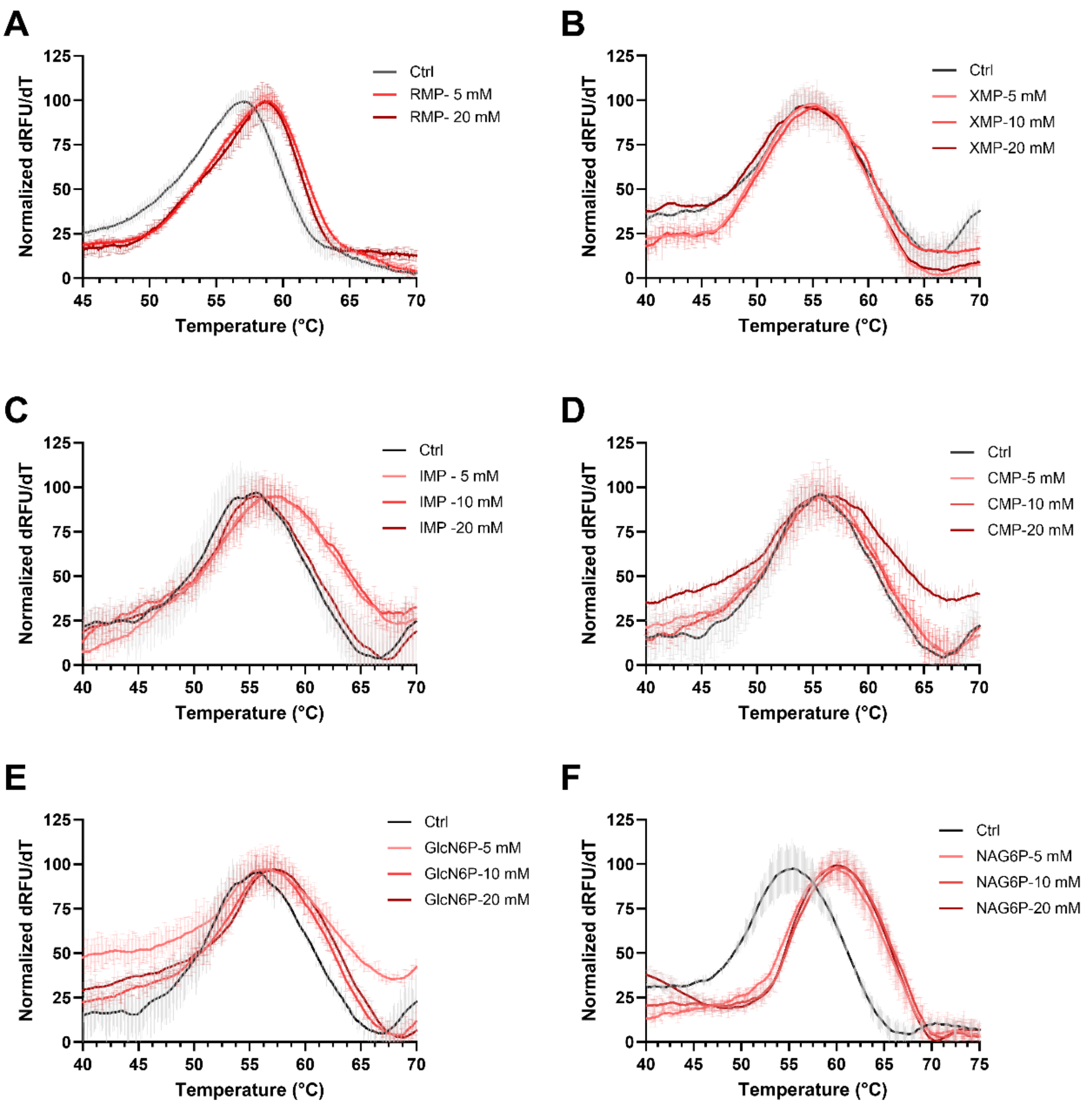


**Figure S 2: Differential scanning fluorometry (DSF) analysis of human glucose-6-phosphate isomerase (GPI) in the absence and presence of ligands**. (A) Ribavirin 5'-monophosphate (RMP). (B) Xanthosine 5'-monophosphate (XMP). (C) Inosine 5'-monophosphate (IMP). (D) Cytidine 5'-monophosphate (CMP). (E) D-Glucosamine 6-phosphate (GlcN6P). (F) N-Acetyl-D-glucosamine 6-phosphate (NAG6P). Protein thermal denaturation profiles are shown as the first derivative of the normalized fluorescence signal (normalized dRFU/dT) as a function of temperature (°C). Each curve represents the mean ± SD of three replicates.

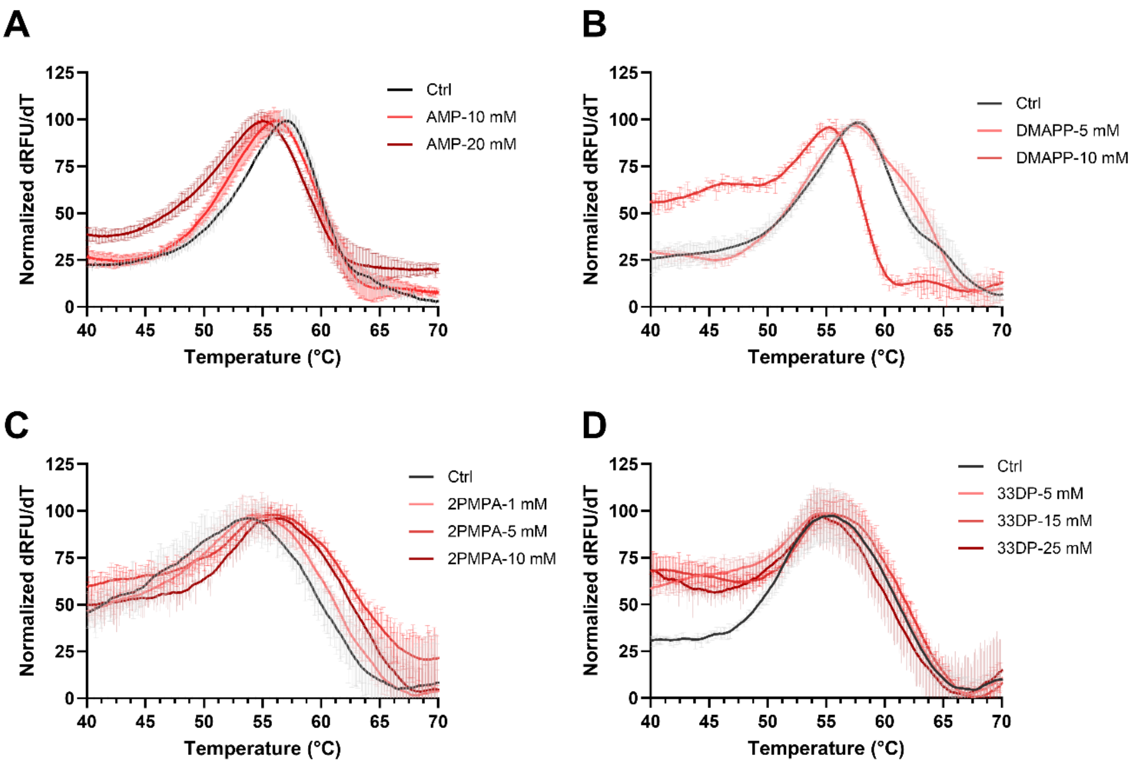


Figure S 3: Differential scanning fluorometry (DSF) analysis of human glucose-6-phosphate isomerase (GPI) in the absence and presence of ligands. (A) Adenosine 5'-monophosphate (AMP). (B) Dimethylallyl pyrophosphate (DMAPP). (C) 2-Phosphonomethyl pentanedioic acid (2PMPA). (D) 3,3-diphosphono-propanoic acid (33DP). Protein thermal denaturation profiles are shown as the first derivative of the normalized fluorescence signal (normalized dRFU/dT) as a function of temperature (°C). Each curve represents the mean ± SD of three replicates.

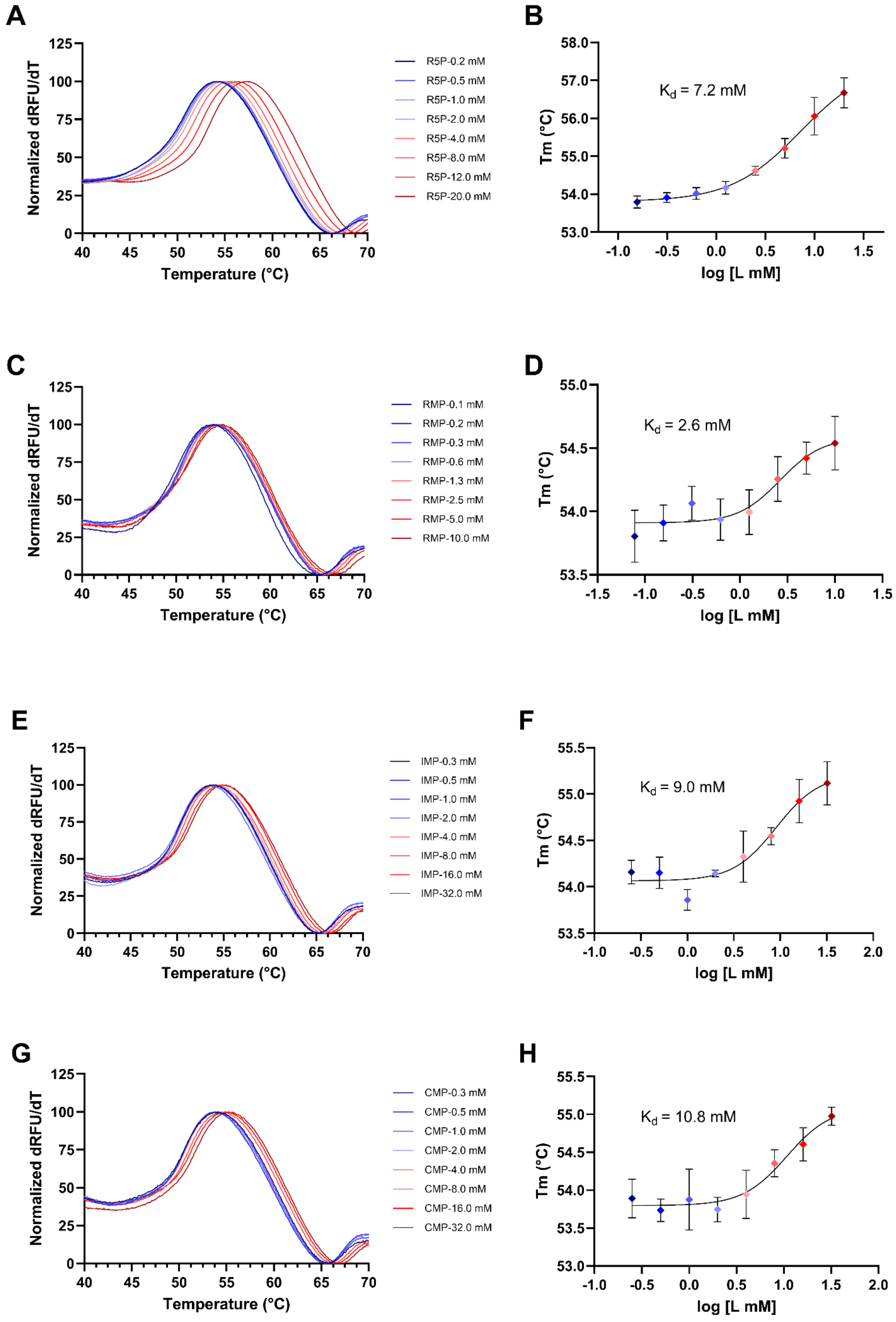

A
Normalized dRFU/dT
Temperature (°C)
R5P-0.2 mM
R5P-0.5 mM
R5P-1.0 mM
R5P-2.0 mM
R5P-4.0 mM
R5P-8.0 mM
R5P-12.0 mM
R5P-20.0 mM
B
Tm (°C)
log [L mM]
Kd = 7.2 mM
C
Normalized dRFU/dT
Temperature (°C)
RMP-0.1 mM
RMP-0.2 mM
RMP-0.3 mM
RMP-0.6 mM
RMP-1.3 mM
RMP-2.5 mM
RMP-5.0 mM
RMP-10.0 mM
D
Tm (°C)
log [L mM]
Kd = 2.6 mM
E
Normalized dRFU/dT
Temperature (°C)
IMP-0.3 mM
IMP-0.5 mM
IMP-1.0 mM
IMP-2.0 mM
IMP-4.0 mM
IMP-8.0 mM
IMP-16.0 mM
IMP-32.0 mM
F
Tm (°C)
log [L mM]
Kd = 9.0 mM
G
Normalized dRFU/dT
Temperature (°C)
CMP-0.3 mM
CMP-0.5 mM
CMP-1.0 mM
CMP-2.0 mM
CMP-4.0 mM
CMP-8.0 mM
CMP-16.0 mM
CMP-32.0 mM
H
Tm (°C)
log [L mM]
Kd = 10.8 mM

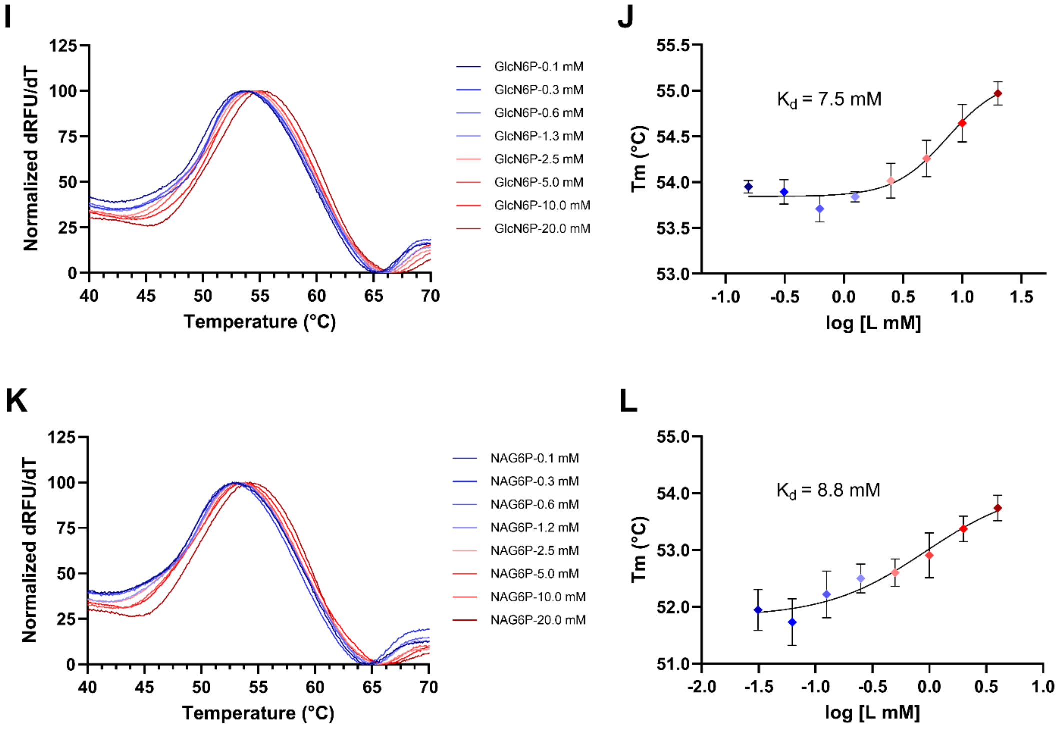


Figure S 4: Differential scanning fluorometry (DSF) titration of human glucose-6-phosphate isomerase (GPI) with selected ligands: (A, B) Ribose-5-phosphate (R5P). (C, D) Ribavirin 5'-monophosphate (RMP). (E, F) Inosine 5'-monophosphate (IMP). (G, H) Cytidine 5'-monophosphate (CMP). (I, J) D-Glucosamine 6-phosphate (GlcN6P). (K, L) N-acetyl glucosamine-6-phosphate (NAG6P). Protein thermal denaturation was monitored in the presence of an 8-point ligand concentration series. Thermal curves are shown as normalized first-derivative fluorescence signals (dRFU/dt) plotted against temperature (°C). Data represent the mean ± SD of two replicates. Melting temperature ($T_m$) were used for estimation of binding affinity ($K_d$).

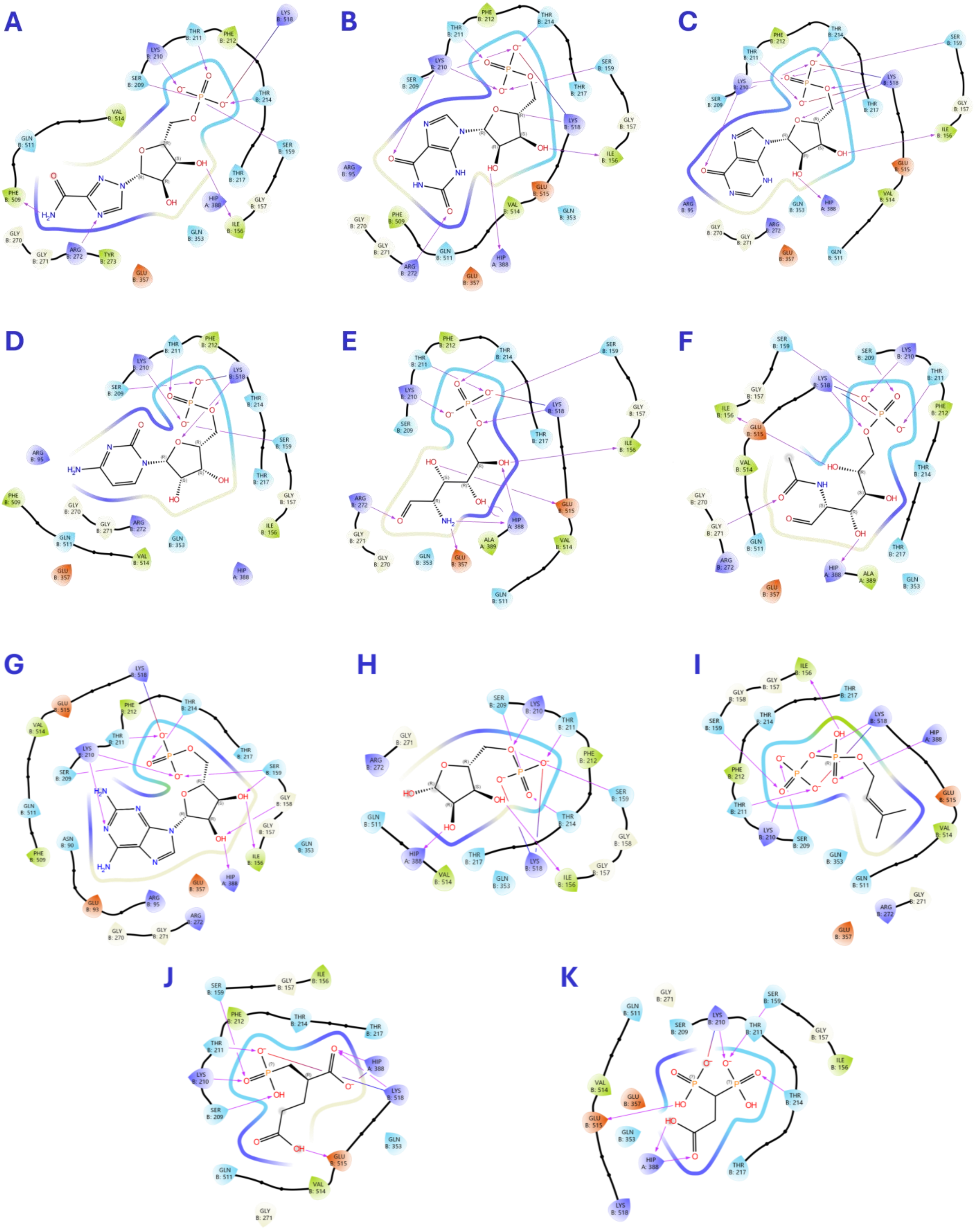


**Figure S 5: Ligand interaction diagrams showing the predicted binding modes of 11 confirmed hits docked into GPI active site.** (A) Ribavirin 5'-monophosphate (RMP). (B) Xanthosine 5'-monophosphate (XMP). (C) Inosine 5'-monophosphate (IMP). (D) Cytidine 5'-monophosphate (CMP). (E) D-Glucosamine 6-phosphate (GlcN6P). (F) N-Acetyl-D-glucosamine 6-phosphate (NAG6P). (G) Adenosine 5'-monophosphate (AMP). (H) D-ribose-5-phosphate. (I) Dimethylallyl pyrophosphate (DMAPP). (J) 2-Phosphonomethyl pentanedioic acid (2PMPA). (K) 3,3-

diphosphono-propanoic acid (33DP).

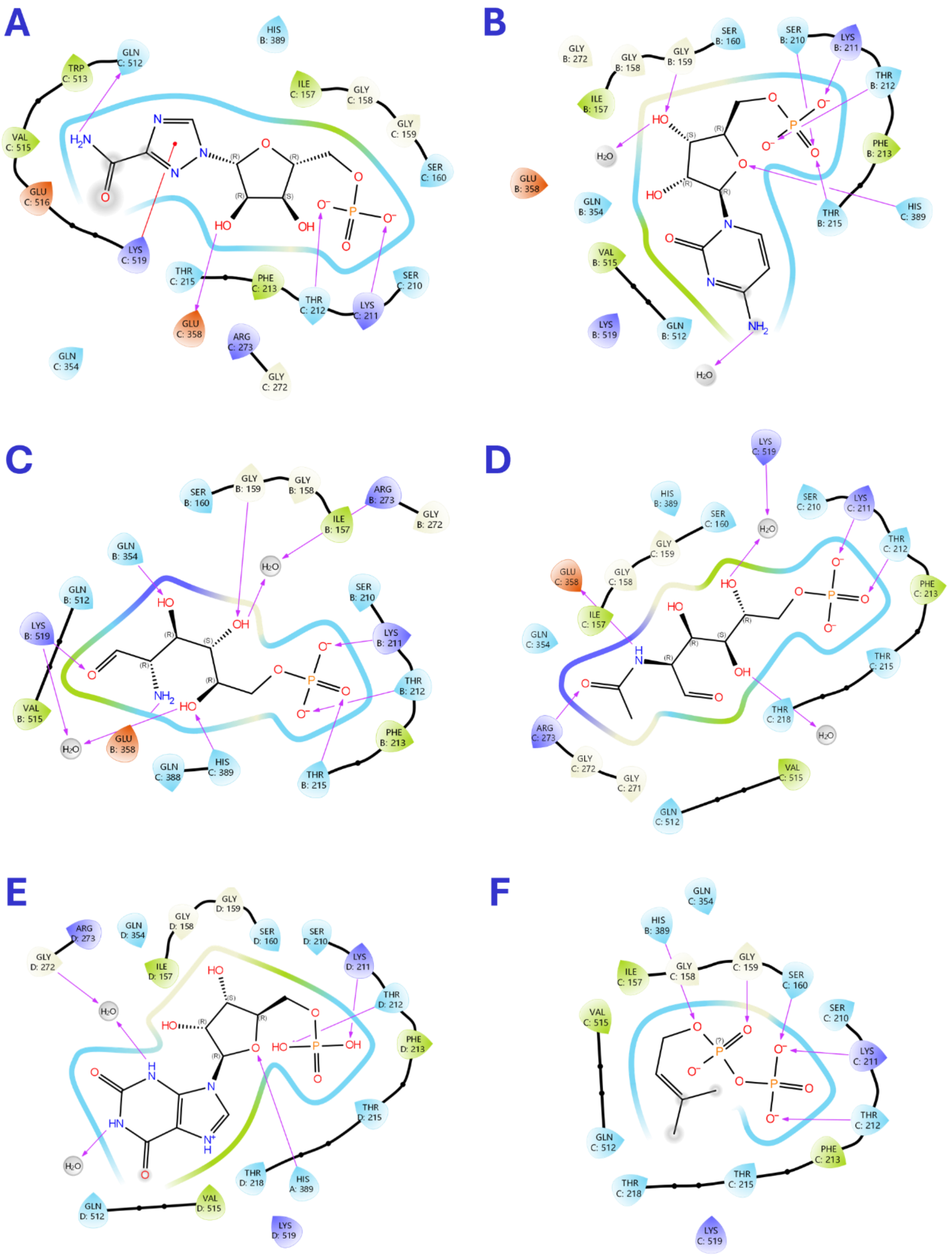


**Figure S 6: Ligand interaction diagrams showing the binding of ligands to human glucose-6-phosphate isomerase in crystal complexes**. (A) Ribavirin 5'-monosphosphate (RMP; PDB: 32BF). (B) Cytidine 5'-monosphosphate (CMP; PDB: 32JZ). (C) D-glucosamine-6-phosphate (GlcN6P; PDB: 32BE). (D) N-acetyl-glucosamine-6-phosphate (NAG6P; PDB: 32JQ). (E) Xanthosine 5'-phosphate (XMP; PDB: 32YR). (F) Dimethylallyl pyrophosphate (DMAPP; PDB: 32XF). The interaction diagrams were generated in Schrödinger Maestro.

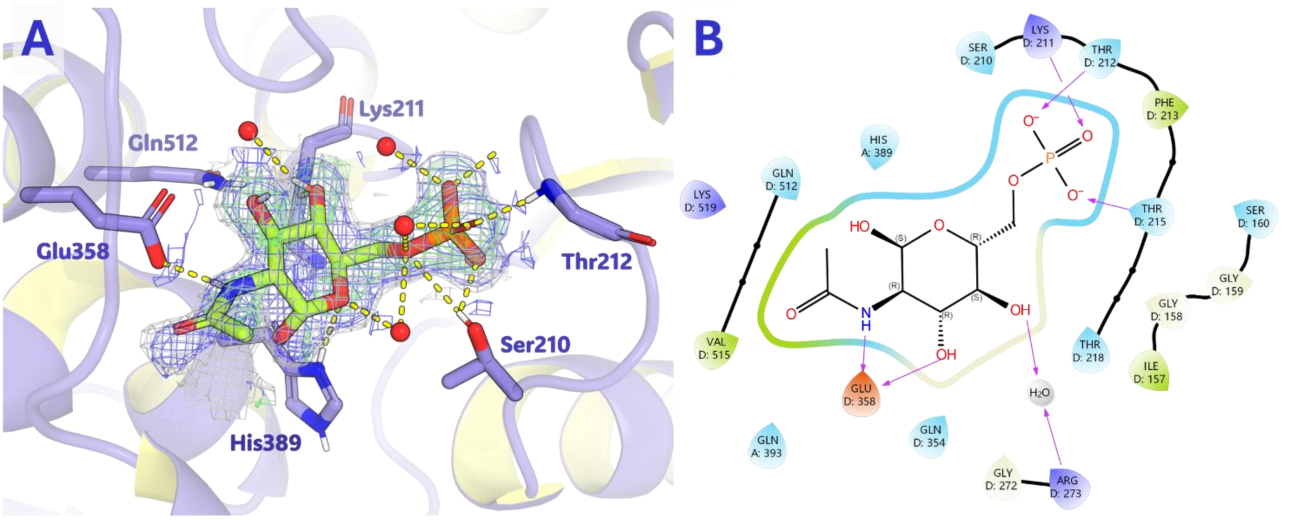


Figure S 7: Binding of N-acetyl-glucosamine-6-phosphate (NAG6P; PDB: 32JQ) – cyclic form to human glucose-6-phosphate isomerase (GPI). (A) Binding pose of NAG6P within the active site of GPI. (B) Ligand interaction diagram showing the contacts formed between NAG6P and GPI active-site residues.

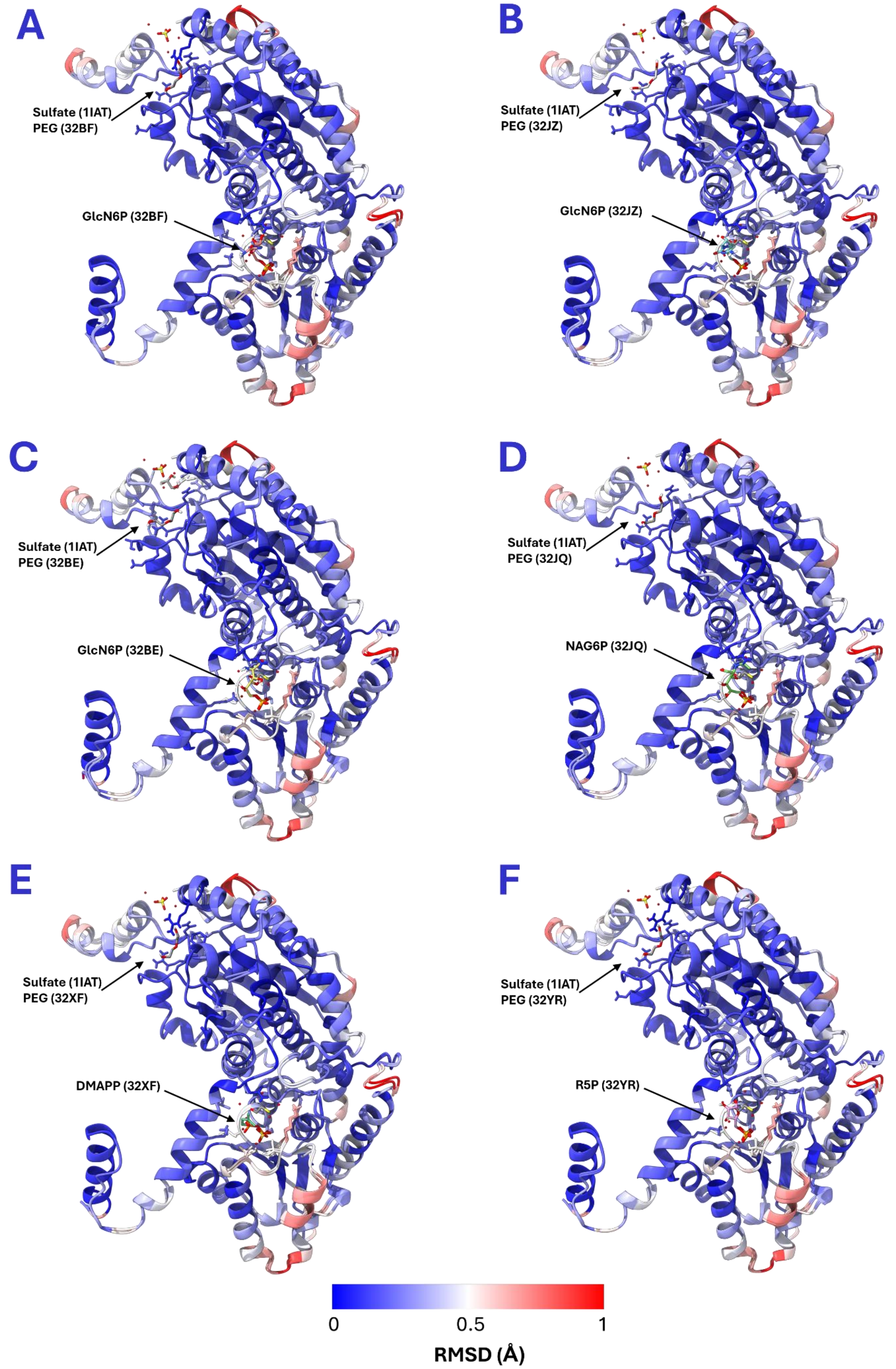


**Figure S 8: RMSD-based color rendering of aligned human glucose-6-phosphate isomerase (GPI).** The apo human GPI structure (PDB: 1IAT) was superimposed with chain B of different GPI–ligand complexes containing (A) Ribavirin 5'-

monosphosphate (RMP; PDB: 32BF). (B) Cytidine 5'-monosphosphate (CMP; PDB: 32JZ). (C) D-glucosamine-6-phosphate (GlcN6P; PDB: 32BE). (D) N-acetyl-glucosamine-6-phosphate (NAG6P; PDB: 32JQ). (E) Dimethylallyl pyrophosphate (DMAPP; PDB: 32XF). (F) Xanthosine 5'-phosphate (XMP; PDB: 32YR). Structural alignments, RMSD calculations, and figure generation were performed using UCSF Chimera.

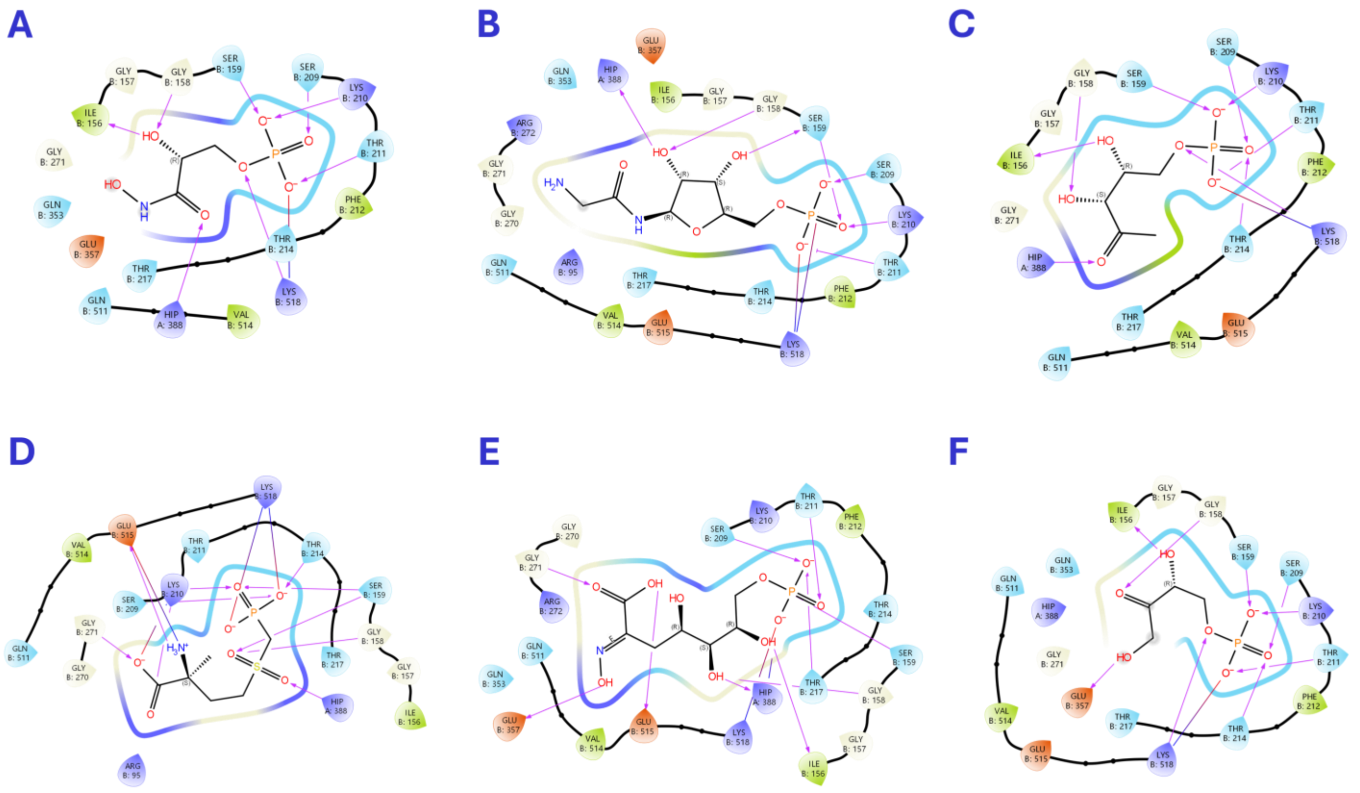

Figure S 9: Examples of virtual screening hits that showed good promises as GPI active-site binders but were not available for testing. (A) CID85778437: -106.72 kJ/mol. (B) MCULE-8005768824: -130.02 kJ/mol. (C) ZINC01532265: -113.99 kJ/mol. (D) ZINC06482071: -123.65 kJ/mol. (E) ZINC584905133: -121.48 kJ/mol. (F) ZINC0100897693: -104.71 kJ/mol.

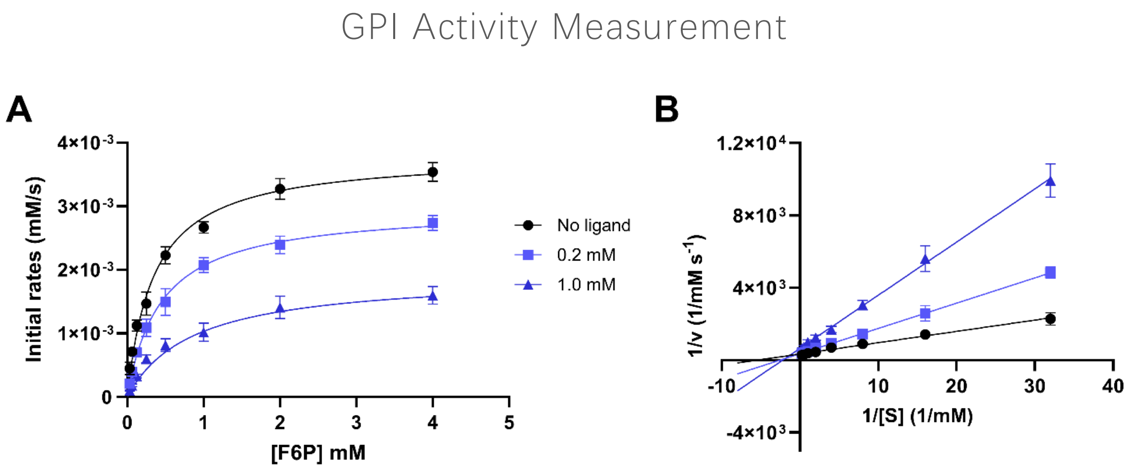


Figure S 10: Michaelis–Menten kinetics (A) and Lineweaver–Burk (B) analysis of human glucose-6-phosphate

**isomerase (GPI) inhibition by 6-phosphogluconate (6PG).** Inhibitory potency ($K_i$) estimated to be approximately 106 µM. Data were fitted using Michaelis–Menten nonlinear regression and linear regression in GraphPad Prism. Each data represent the mean of duplicate measurements, and error bars indicate the standard deviation (± SD).

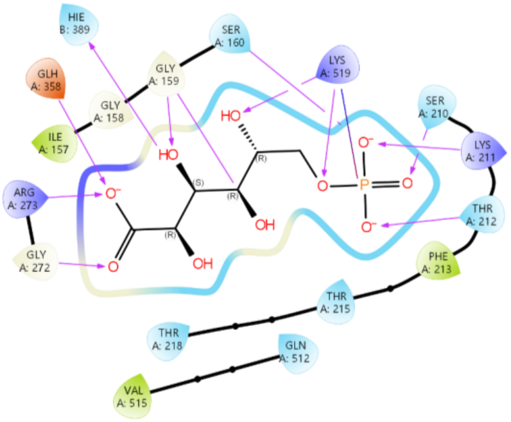


**Figure S 11: Ligand interaction diagram of 6-phosphogluconate (6PG) bound in the active site of mouse glucose-6-phospahte isomerase** (PDB: 2CXR). The interaction diagram was generated in Schrödinger Maestro.

Extracellular Flux Measurement Results

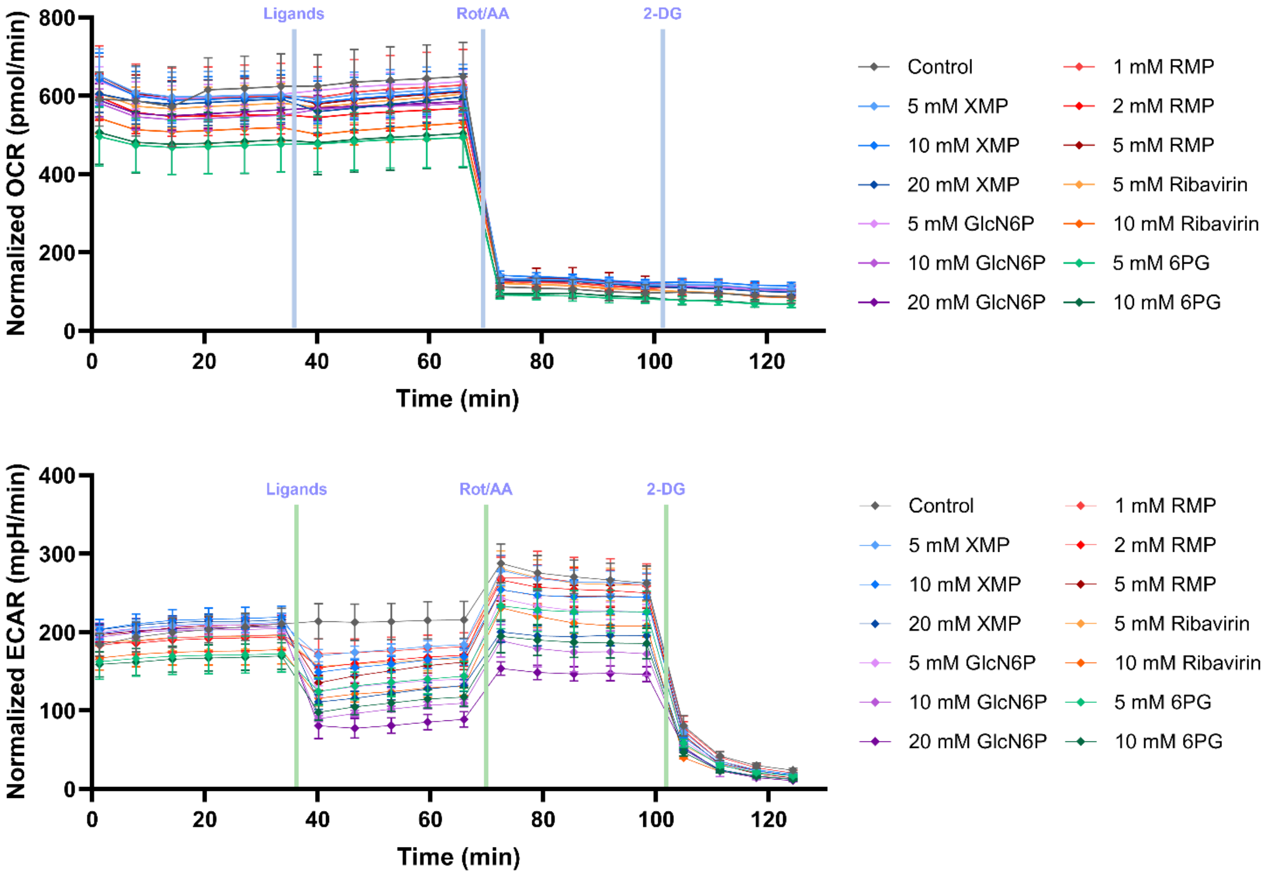

**Figure S 12: Real-time measurement of HEK-293 oxygen consumption rate (OCR; upper panel) and extracellular acidification rate (ECAR; lower panel)** at 37°C, before and after the injection of effectors and inhibitors. The OCR and ECAR values were normalized to relative viable cells in each well, quantified by Prestoblue staining. Each data point represents the mean ± SD of five replicates.

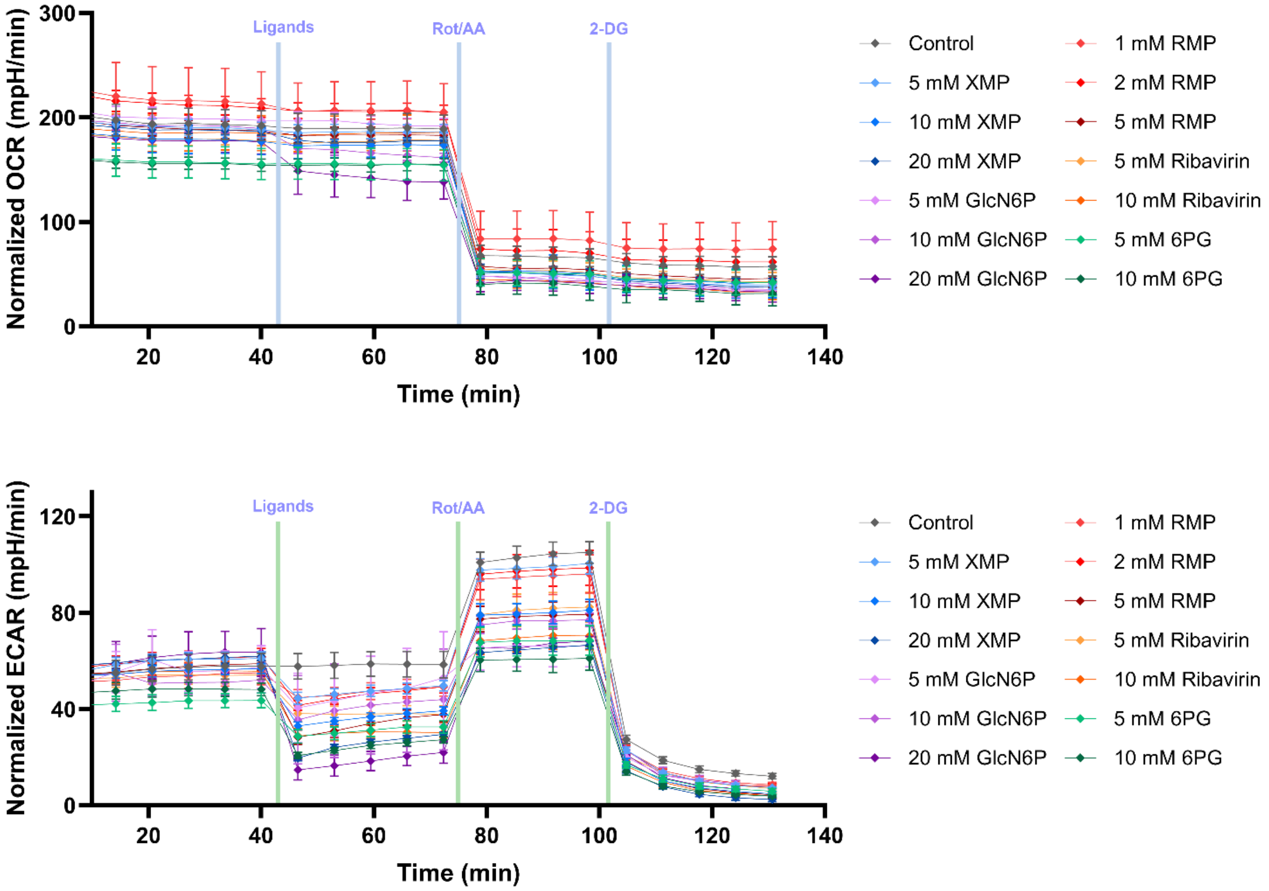


**Figure S 13: Real-time measurement of U2OS oxygen consumption rate (OCR; upper panel) and extracellular acidification rate (ECAR; lower panel)** at 37°C, before and after the injection of effectors and inhibitors. The OCR and ECAR values were normalized to relative viable cells in each well, quantified by Prestoblue staining. Each data point represents the mean ± SD of five replicates.

#### Pre-docking validation

##### Cross-docking results for hydrated GPI grids

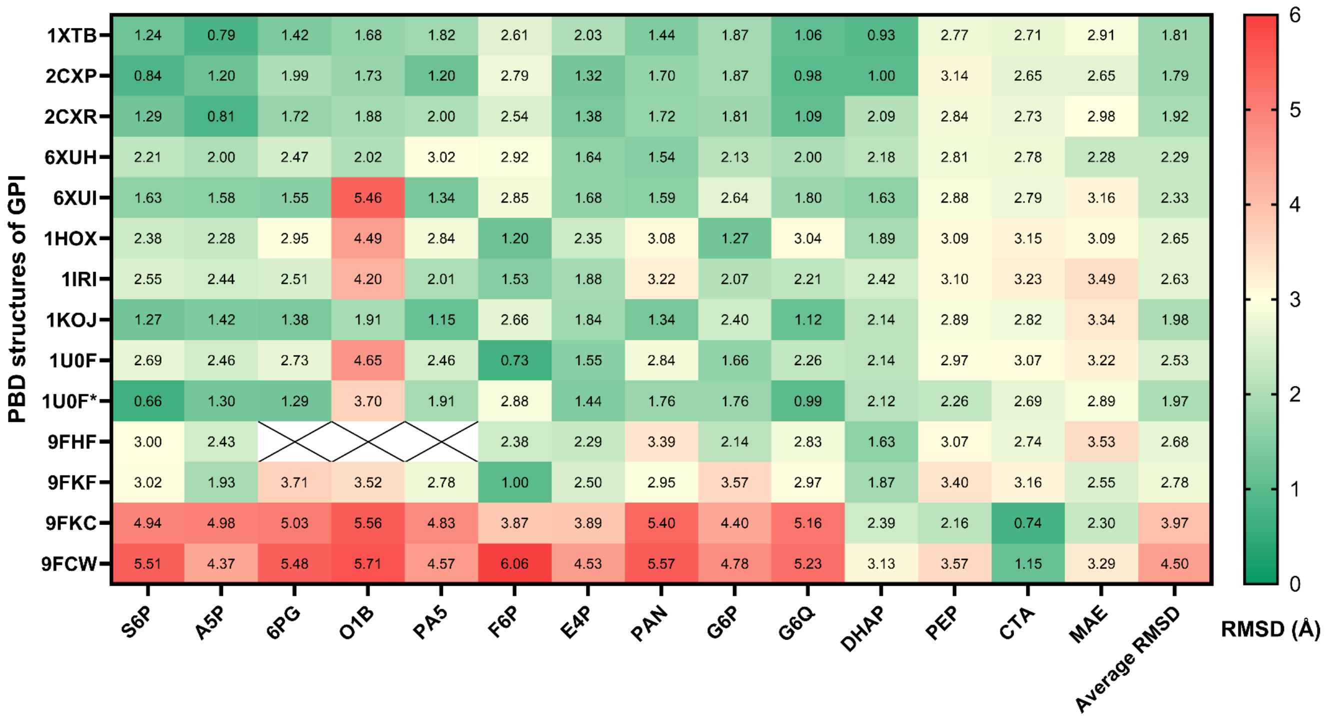


**Figure S 14: Cross-docking study performed for known structures of human and mammalian GPI in the presence of water molecules.** Active-site grids were generated with crystallographic waters retained. Cognate ligands were docked into the hydrated grids using Schrödinger Glide in Standard Precision (SP) mode with flexible ligand sampling to evaluate the accuracy of the docking protocol. The root-mean-square deviation (RMSD, Å) between the docked and crystallographic ligand poses is shown in each cell.

##### Cross-docking results for dry GPI grids

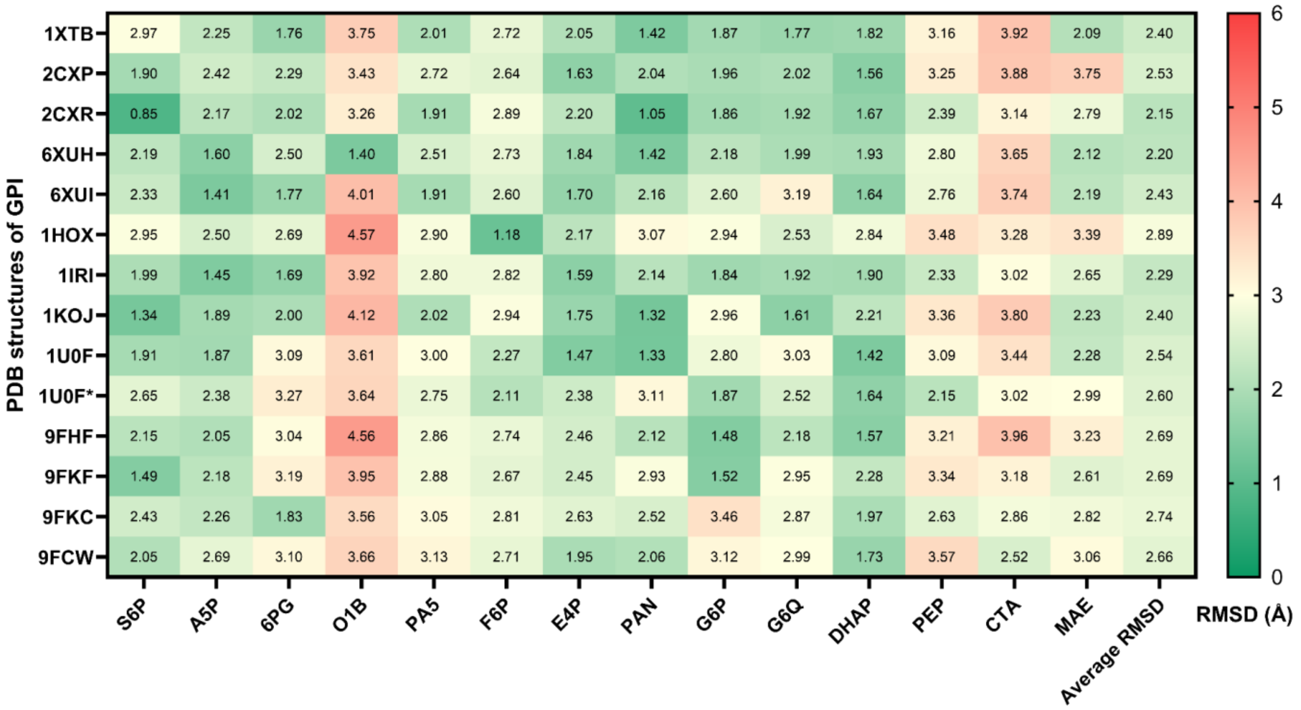


Figure S 15: Cross-docking study performed for known structures of human and mammalian GPI in the absence of water molecules. Active-site grids were generated after removal of crystallographic waters. Cognate ligands were docked into the dry grids using Schrödinger Glide in Standard Precision (SP) mode with flexible ligand sampling to evaluate the accuracy of the docking protocol. The root-mean-square deviation (RMSD, Å) between the docked and crystallographic ligand poses is shown in each cell.

##### Cross-docking result comparison

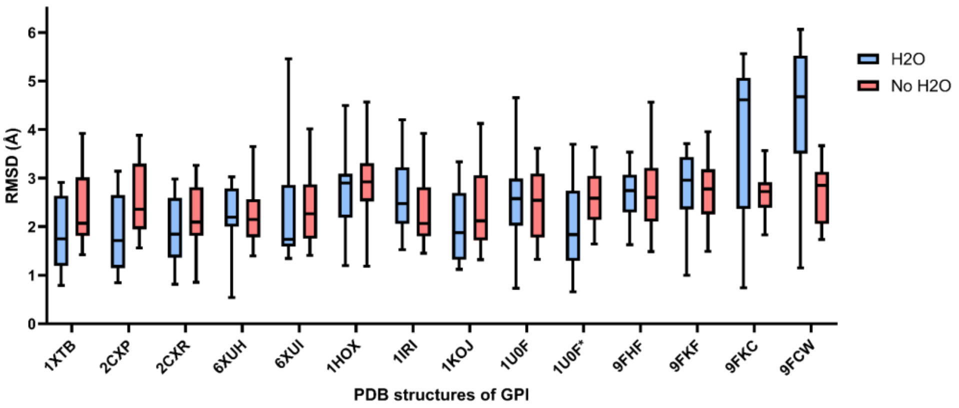


Figure S 16: Comparison of cross-docking performance using hydrated and dry GPI active-site grids. Docking accuracy indicted by the RMSD (Å) between the docked and crystallographic ligand poses for hydrated and dry grids.

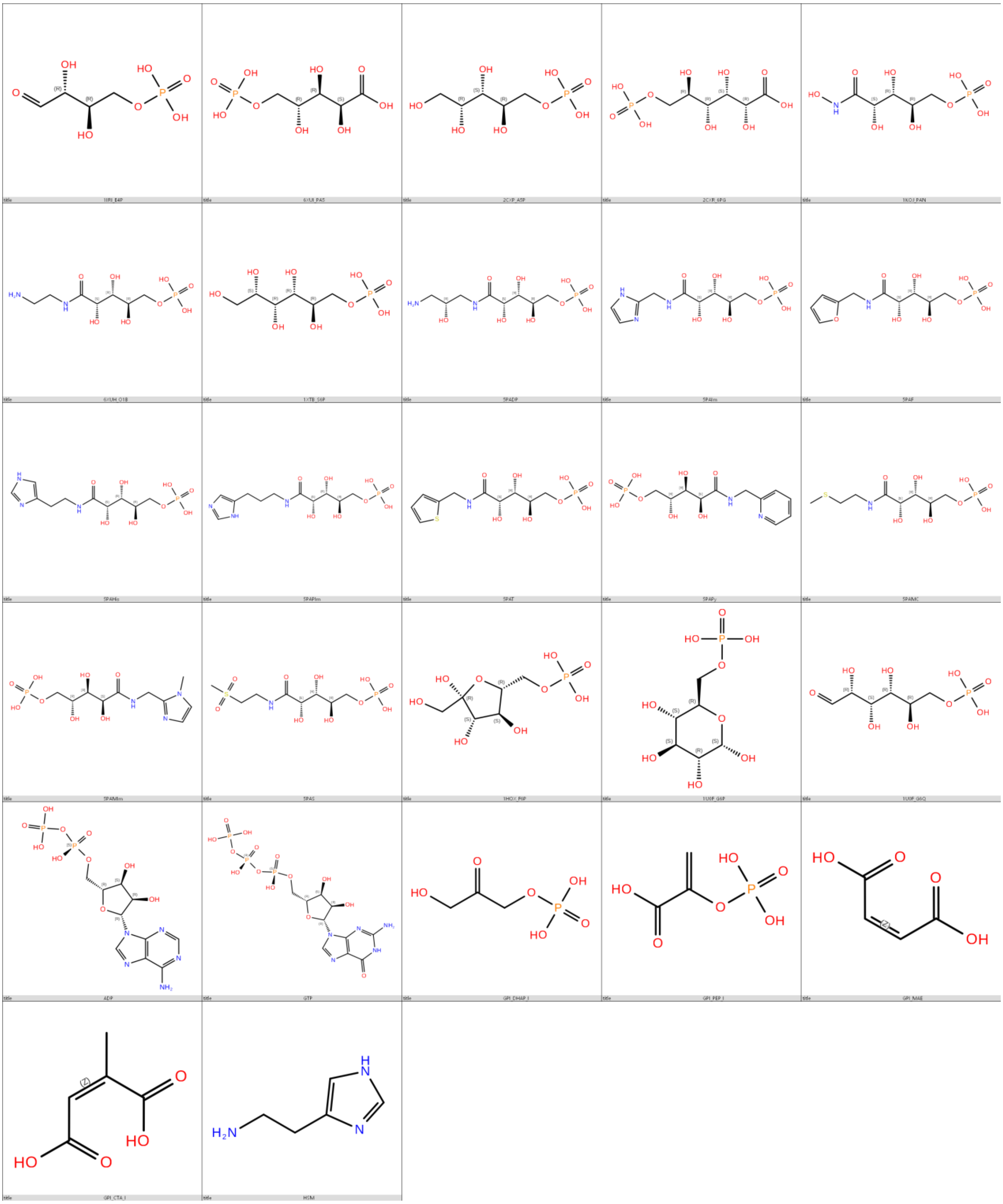

**Figure S 17: Chemical structures of previously reported human and mammalian GPI inhibitors**. Detailed information for each compound, including physicochemical properties and docking scores, is provided in the supplementary excel file "Supplementary_A_known-inhibitors".

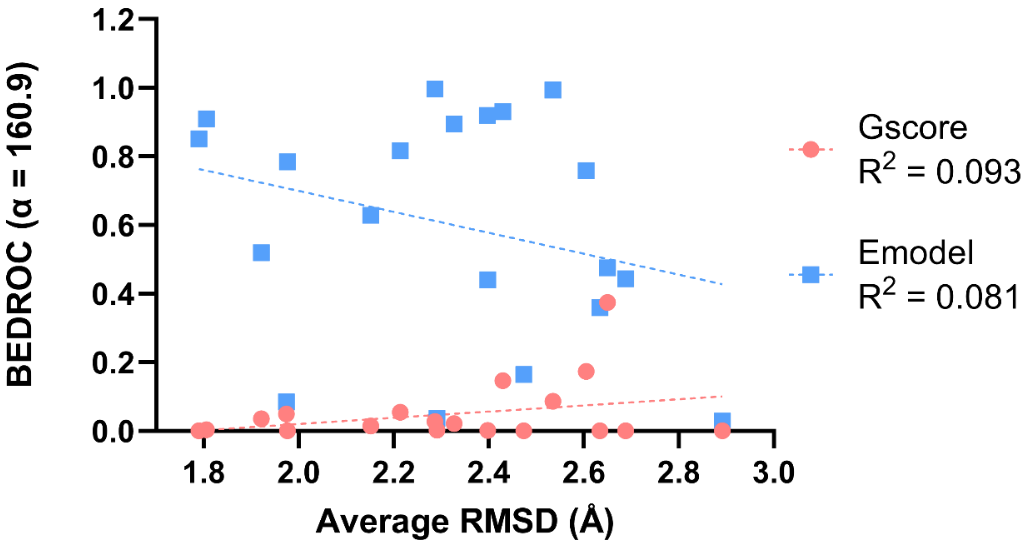


Figure S 18: Relationship between docking scores and docking accuracy. Scatter plots showing the correlation between (pink) GlideScore and RMSD and (blue) Emodel and RMSD for all cross-docking poses. RMSD (Å) values were calculated relative to the crystallographic ligand poses. $R^2$ (Gscore) = 0.093; $R^2$ (Emodel) = 0.081

**Summary**: No correlation was observed between pose RMSD and early enrichment performance - BEDROC (α = 160.9), as indicated by low $R^2$ values for both Gscore ($R^2$ = 0.093) and Emodel ($R^2$ = 0.081), suggesting that accurate pose prediction does not necessarily translate into improved early enrichment. Although not optimized for enrichment, the Emodel scoring function outperformed the Gscore function in active compound enrichment across different docking grids, regardless of the presence of water molecules. Better performance of Emodel over Gscore scoring function likely arises from its composite nature, as it integrates multiple energetic and geometric terms (e.g., pose quality and ligand strain), making it more effective in distinguishing active compounds during ranking-based enrichment compared to the Gscore function, which has been designed for database enrichment and binding affinity prediction.
